# Native-like soluble E1E2 glycoprotein heterodimers on self-assembling protein nanoparticles for hepatitis C virus vaccine design

**DOI:** 10.1101/2025.05.16.654559

**Authors:** Linling He, Yi-Zong Lee, Yi-Nan Zhang, Maddy L. Newby, Benjamin M. Janus, Fabrizio G. Gonzalez, Garrett Ward, Connor DesRoberts, Shr-Hau Hung, Erick Giang, Joel D. Allen, Liudmila Kulakova, Eric A. Toth, Thomas R. Fuerst, Mansun Law, Gilad Ofek, Max Crispin, Jiang Zhu

**Affiliations:** Department of Integrative Structural and Computational Biology, The Scripps Research Institute, La Jolla, CA 92037, USA; Department of Cell Biology and Molecular Genetics, University of Maryland, College Park, MD, 20742, USA; Institute for Bioscience and Biotechnology Research, University of Maryland, Rockville, MD, 20850, USA; School of Biological Sciences, Highfield Campus, University of Southampton, Southampton, SO17 1BJ, UK; Department of Immunology and Microbiology, The Scripps Research Institute, La Jolla, CA 92037, USA

## Abstract

Hepatitis C virus (HCV) is a leading cause of chronic liver disease, cirrhosis, and hepatocellular carcinoma worldwide. E1E2-based HCV vaccine development has been hindered by the challenge of producing a soluble E1E2 (sE1E2) antigen that faithfully recapitulates the native glycoprotein heterodimer found on virions. Based on available cryo-electron microscopy (cryo-EM) structures, we rationally engineered sE1E2 for genotype 1a H77 by truncating the E1 and E2 stems (Cut_1_), removing a putative fusion peptide (pFP)-containing region in E1 (Cut_2_), and stabilizing the E1-E2 interface with diverse heterodimeric scaffolds. All H77 sE1E2.Cut_1+2_ scaffolds showed native-like E1-E2 association and robust binding to the broadly neutralizing antibody (bNAb) AR4A. A genotype 1a HCV-1 sE1E2.Cut_1+2_ variant scaffolded by a modified SpyTag/SpyCatcher (SPYΔN) was selected for in vitro, structural, and immunogenic characterization. The structure of this sE1E2 scaffold in complex with bNAbs was analyzed by cryo-EM and negative-stain EM (nsEM), with an nsEM-based approach developed for antibody epitope mapping. HCV-1 sE1E2.Cut_1+2_.SPYΔN was displayed on self-assembling protein nanoparticles (SApNPs) to enhance immunogenicity. HCV-1 sE1E2.Cut_1+2_.SPYΔN heterodimer and SApNPs with wildtype and modified glycans were tested in mice, revealing the beneficial effects of multivalent display and oligomannose enrichment. Our study provides a rigorous foundation for next-generation HCV vaccine development.

**ONE-SENTENCE SUMMARY:** Rational design, characterization, and in vivo assessment of HCV soluble E1E2 heterodimer and nanoparticles will inform vaccine development.

## INTRODUCTION

According to the World Health Organization (WHO), an estimated 50 million people worldwide are chronically infected with hepatitis C virus (HCV), with approximately 1 million new infections occurring annually (*1*). In 2022, about 240,000 deaths were reported, primarily due to cirrhosis and hepatocellular carcinoma, making HCV one of the leading causes of liver-related morbidity and mortality (*1*). In the United States, the growing opioid and injection drug use has contributed to more than 70,000 overdose-related deaths in 2019 (*2*) and a sharp rise in acute HCV infections (*3*). Over the past decade, direct-acting antiviral (DAA) therapies have demonstrated remarkable success in treating chronic HCV infection and achieving sustained virologic response (SVR) rates (*4–6*). Although DAAs have been shown to improve liver function in patients with decompensated cirrhosis (*7, 8*), they do not prevent HCV reinfection or reduce the risk of liver cancer (*9–11*). Finally, despite improvements in diagnostic technologies (*12*), early detection of asymptomatic HCV infections remains a challenge, and treatment is often not initiated until individuals have already developed liver damage or cirrhosis (*13*). Thus, an effective prophylactic HCV vaccine is urgently needed to achieve the WHO’s goal of hepatitis elimination by 2030 (*14, 15*).

HCV belongs to the *Hepacivirus* genus in the *Flaviviridae* family, which comprises small, enveloped, positive-sense single-stranded RNA viruses that infect rodents, canines, primates, and other species (*16–18*). The HCV genome encodes a single polyprotein that is processed into seven non-structural (NS) proteins and three structural proteins, including the capsid-forming core protein and two envelope glycoproteins (Env) anchored in the viral membrane (*19–21*). HCV entry into host hepatocytes is mediated by the Env heterodimer, E1E2 (*22, 23*), in which E2 binds to host cellular receptors CD81 and scavenger receptor class B member 1 (SR-B1) (*24*), as well as low-density lipoprotein receptor (LDLr) as a co-receptor (*25*), while E1 tightly associates with E2 to facilitate E1E2 formation and membrane fusion and participates in various steps of the viral life cycle (*26, 27*). The development of effective HCV therapeutics and vaccines has been hampered by the virus’s inherent genetic diversity, driven by low polymerase fidelity and rapid replication, as evidenced by the existence of eight genotypes and 93 subtypes (*28*). Upon HCV infection, rapid mutation of E1E2 and NS5A helps establish a population of related but distinct quasispecies (*29*), enabling viral evasion from host neutralizing antibodies (NAbs) and leading to resistance to DAAs (*30*). Nonetheless, HCV infection can spontaneously clear in 20-25% of cases, with both T cells and NAbs contributing to effective virus control (*31*). However, a recent phase 1/2 trial evaluating recombinant viral vectors encoding an NS-based antigen (NSmut) failed to prevent chronic HCV infection, underscoring the limitations of T cell-focused vaccine strategies. Early NAb studies (*32–38*) demonstrated the critical role of broadly neutralizing antibodies (bNAbs) in protection against HCV, a finding that has been increasingly appreciated in vaccine design (*39–43*).

Structural studies of antibody-bound Env proteins have revealed how the humoral immune system recognizes sites of HCV vulnerability, forming the basis for rational vaccine design against this highly mutable virus (*44–46*). Peptide/NAb complex structures (*47–51*) have provided insights into key linear neutralizing epitopes on E1 and E2, enabling epitope grafting onto non-viral protein scaffolds, including nanoparticles (NPs) (*52–54*). The crystal structures of two E2 ectodomains, an E2 core (E2c) from the H77 isolate (genotype 1a) stabilized by bNAb AR3C, and a truncated E2 from the J6 isolate (genotype 2a) bound to the non-NAb 2A12, marked a milestone in HCV research (*55, 56*). Overall, the two E2 structures shared a similar fold despite differences in variable region 3 (VR3) and disulfide bonding patterns (*57*). The crystal structure of a more complete E2 offered further insight into this viral glycoprotein (*58*). Stable E2c constructs enabled the structural characterization of human bNAbs and the definition of major E2 epitopes (*46, 59, 60*), including antigenic site 412-423 (AS412), antigenic site 434-446 (AS434, part of the E2 front layer [FL]), and antigenic region 3 (AR3). A conserved E2 surface comprising most of the FL and the CD81 binding loop (CD81bl) forms a neutralizing face (NF) targeted by antibodies during infection (*59, 61*). Notably, AR3-directed bNAbs preferentially use the V_H_1-69 gene in humans and equivalent heavy-chain variable (V_H_) genes in nonhuman primates (NHPs) (*62–65*), supporting a germline-targeting strategy for AR3-focused HCV vaccine design (*40*). Additionally, variability in epitope conformations and antibody binding modes has been associated with NF recognition by VH1-69–derived NAbs (*62, 65, 66*). The growing body of structural data on E2 has guided the optimization of soluble E2 (sE2) and E2c constructs to improve epitope-specific NAb responses and advance E2-based NP vaccines (*67–69*). While E2 remains a major vaccine target, recent high-resolution structures of E1E2 determined by cryo-electron microscopy (cryo-EM) have, for the first time, revealed atomic-level details of the E1 structure and the E1–E2 interface (*70–72*), signaling a shift toward E1E2-based HCV vaccines (*39, 73–75*). In the full-length E1E2 structure, the E1 and E2 stems form an extended interface positioned above the viral membrane (*71*). Scaffolding the E1 and E2 ectodomains with a heterodimeric coiled coil produced the first soluble E1E2 (sE1E2) reactive with bNAb AR4A (*32*)—which requires a native-like E1−E2 interface (*76, 77*)—and enabled the determination of a 3.65 Å resolution cryo-EM structure for another sE1E2 scaffold (*70*). The recently reported dimer-of-dimer E1E2 structure (*72*) added further complexity to HCV vaccine design. Nonetheless, these structural advances have spurred efforts to develop E1E2 NP vaccines (*75, 78, 79*). Still, E1E2-based HCV vaccine design remains in its early stages (*73*).

In our previous studies, we established a rational vaccine design strategy for class I fusion viruses (*80, 81*) by combining antigen optimization, protein NP display, and glycan modification. As demonstrated for the HIV-1 Env (*82–84*), filovirus glycoprotein (*85, 86*), SARS-CoV-1/2 spike (*87*), and RSV fusion protein (*88*), we first identified the causes of metastability through structural analysis and stabilized the glycoproteins with targeted mutations. We then displayed these antigens on multilayered single-component self-assembling protein NPs (SApNPs), which were engineered from 60-meric bacterial proteins E2p and I3-01, as multivalent vaccine candidates (*67, 82, 83, 85–87, 89–91*). Our recent work also demonstrated the benefits of glycan trimming and oligomannose enrichment on HIV-1 and filovirus vaccines, respectively (*82, 85*). Here, we adapted this strategy to HCV, a virus with a noncanonical fusion mechanism. First, using available E1E2 structures (*70, 71*) as templates, we truncated E1 at H312 and E2 at Y701 and deleted an unstructured E1 region encompassing the putative fusion peptide (pFP) in genotype 1a H77. The resulting sE1E2.Cut_1_ and sE1E2.Cut_1+2_ constructs were scaffolded using four heterodimeric leucine zippers of different sizes and an N-terminally truncated SpyTag/SpyCatcher (*92*), termed SPYΔN. All H77 sE1E2.Cut_1+2_ scaffolds bound with high affinity to bNAb AR4A (*32*), indicating a native-like E1−E2 interface. We next designed a His_6_-tagged sE1E2.Cut_1+2_.SPYΔN construct for the genotype 1a strain HCV-1 and characterized it biochemically, biophysically, and antigenically. Cryo-EM analysis was performed on this HCV-1 sE1E2 scaffold in complex with bNAbs AR3C (or HEPC74 (*58*)) and AR4A, and a negative-stain EM (nsEM) approach was evaluated for epitope mapping. HCV-1 sE1E2.Cut_1+2_.SPYΔN was displayed multivalently on ferritin 24-mer and I3-01v9a 60-mer (*90*) as virus-like particle (VLP) vaccines. Finally, we immunized mice to assess NAb responses to these rationally designed sE1E2 vaccines with and without glycan modification. The results showed that NP display and oligomannose enrichment positively influenced NAb elicitation. Together, these findings support a promising strategy for developing effective E1E2-based HCV vaccines.

## RESULTS

### Rational design of HCV sE1E2 scaffolds with a native-like E1**−**E2 interface

HCV E1E2 is a noncanonical fusion glycoprotein with an unusual heterodimeric architecture (*93*). Cryo-EM analysis of full-length E1E2 bound to three NAbs provided the first complete view of this elusive viral glycoprotein and paved the way for rational vaccine design (*73*). Although an adjuvanted membrane-bound E1E2 vaccine has been tested in humans (*94*), an sE1E2 construct with a native-like E1−E2 interface would, in principle, offer a more tractable antigen for vaccine development. Recently, heterodimeric coiled coils were used to scaffold HCV E1E2 ectodomains, resulting in measurable AR4A binding (*76*) and enabling the first cryo-EM structure of an sE1E2 heterodimer (*70*). However, AR4A coexpression was required to stabilize the E1−E2 interface and improve folding for full-length E1E2 and a scaffolded sE1E2 antigen in structural studies (*70, 71*), suggesting intrinsic metastability within the E1E2 ectodomains that must be addressed.

Here, we designed stable HCV sE1E2 scaffolds using a rational approach. Superposition of two cryo-EM structures (*70, 71*) revealed a similar E1E2 architecture despite different sequence backbones (**Fig. 1a**), with the E1 and E2 stems unresolved in the scaffolded sE1E2 (*70*). The E1E2 of H77, a prototype genotype 1a isolate, was used to validate the sE1E2 construct design and heterodimeric scaffolds (**Fig. 1b**). For HCV Env, we hypothesized that truncation of the flexible E1 and E2 stems would enable precise structural control of sE1E2 and stabilization of the E1−E2 interface, and that deletion of the unstructured pFP-containing E1 loop would reduce sE1E2 aggregation. To test this hypothesis, we truncated E1 at H312 and E2 at Y701 (termed “Cut_1_”) and replaced the E1 region L264-F293 with a glycine (termed “Cut_2_”) (**Fig. 1b**). For scaffolds, we hypothesized that diverse heterodimeric coiled coils could be used to accommodate sE1E2 and that a covalently linked heterodimer may provide the optimal protein scaffold to stabilize sE1E2 in an irreversible form. In addition to the previously reported LZ (*76*), a 40-aa human c-Fos/c-Jun leucine zipper (*95*), and SZ (*70*), a 45-aa synthetic leucine zipper (*96*), we identified a 30-aa GCP (*97*) and a 21-aa IAAL (*98*), which are designed leucine zippers composed of acidic and basic chains. Previously, we used SpyTag/SpyCatcher (*92*) to attach SARS-CoV-1/2 receptor-binding domains (RBDs) to protein NPs (*87*). Here, we removed its unstructured N-terminus to create a covalently linked heterodimeric scaffold, termed SPYΔN. A total of 10 constructs were designed to present H77 sE1E2.Cut_1_ and sE1E2.Cut_1+2_ on five scaffolds (LZ, SZ, GCP, IAAL, and SPYΔN), with a furin cleavage motif between sE1 and sE2 to promote native-like E1−E2 assembly (**Fig. 1c** and **Fig. S1a**). An LZ scaffold presenting H77 E1E2 ectodomains, termed sE1E2.LZ (**Fig. S1a**), and the original LZ-scaffolded H77 sE1E2 provided by Fuerst and co-workers (*76*), termed sE1E2.LZorg, were included as controls. All in-house-designed sE1E2 scaffolds had a C-terminal His_6_ tag to facilitate purification by immobilized metal affinity chromatography (IMAC).

**Fig. 1.**
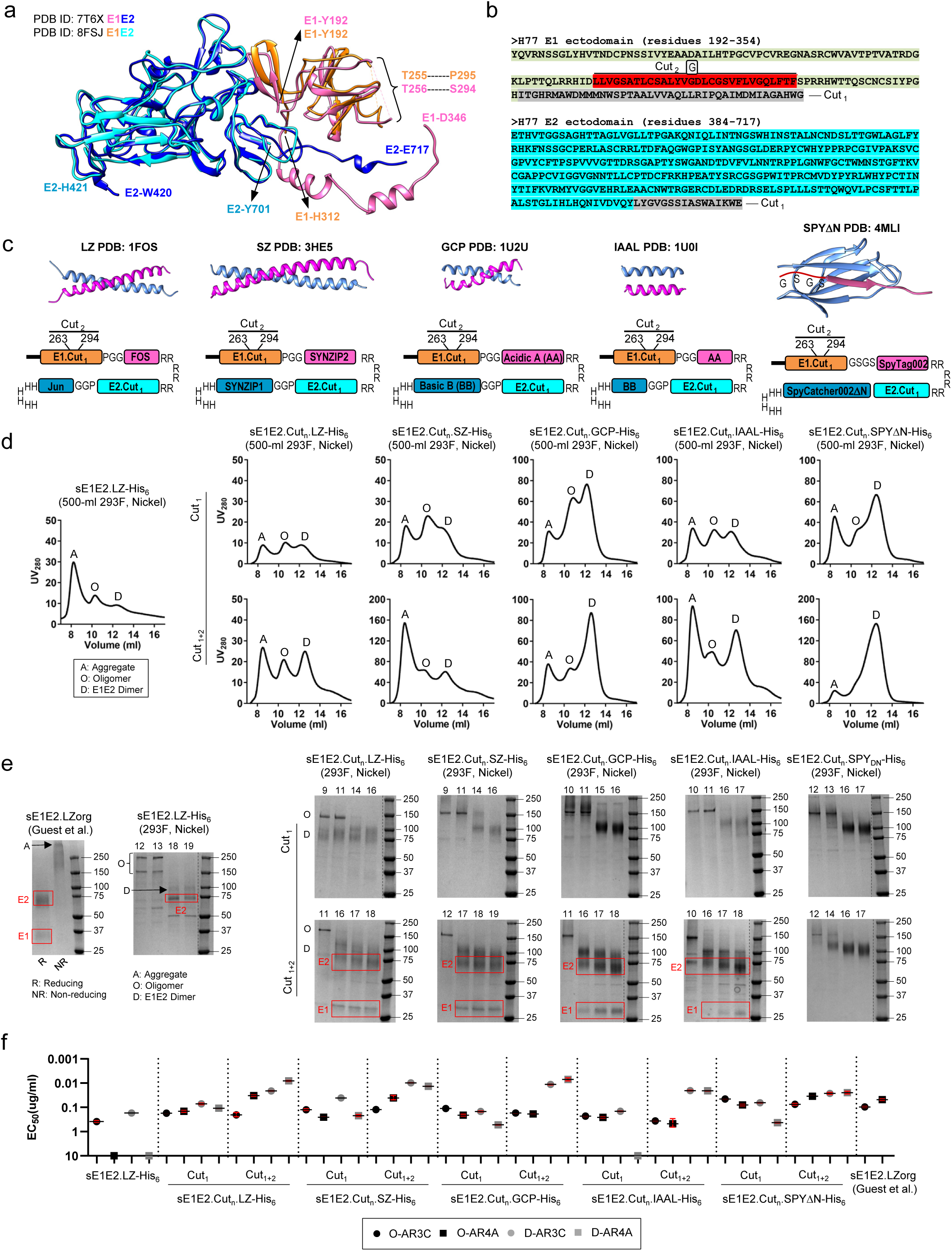
Rational design and in vitro characterization of genotype 1a H77 sE1E2 scaffolds. (**a**) Superimposition of cryo-EM structures of full-length E1E2 (PDB ID: 7T6X) and scaffolded sE1E2 (PDB ID: 8FSJ), both shown in colored ribbon representation. E1 and E2 in the full-length E1E2 are colored magenta and blue, and in the scaffolded sE1E2, orange and cyan, respectively. The structurally resolved N- and C-termini are labeled for both structures, along with the N- and C-terminal anchoring residues of the unstructured, putative fusion peptide (pFP)-containing region in E1. (**b**) Amino acid sequences of genotype 1a H77 E1 and E2 ectodomains, with stem truncation (Cut_1_) and replacement of the pFP-containing E1 loop with a glycine (Cut_2_) indicated. (**c**) Left to right: ribbon representation of a heterodimeric scaffold (top) and schematic representation of the sE1E2.Cut_1_ (or sE1E2.Cut_1+2_) scaffold design (bottom). The two scaffold subunits are colored hot pink and sky blue, except for SPYΔN, in which SpyTag002 shown in maroon. In the schematic, the furin cleavage motif (RRRRRR), linkers (e.g., PGG and GSGS), and His_6_ tag are labeled. (**d**) SEC profiles of H77 sE1E2 scaffolds following transient expression in HEK293F cells and purification using a nickel column. Leftmost: sE1E2.LZ-His_6_; left to right: sE1E2.Cut_1_ (top) and sE1E2.Cut_1+2_ (bottom) on the LZ, SZ, GCP, IAAL, and SPYΔN scaffolds. Peaks corresponding to aggregate (A), oligomer (O), and dimer (D) species are labeled. (**e**) Non-reducing SDS-PAGE analysis of various H77 sE1E2 scaffolds. Leftmost: gels for sE1E2.LZorg, provided by Fuerst and colleagues (Guest et al., PNAS 2021), and SEC fractions of sE1E2.LZ-His_6_ produced in-house without AR4A coexpression; left to right: gels for selected SEC fractions of sE1E2.Cut_1_ (top) and sE1E2.Cut_1+2_ (bottom) on the LZ, SZ, GCP, IAAL, and SPYΔN scaffolds. Dissociated sE1 and sE2 bands are outlined with red line boxes. (**f**) ELISA-derived EC_50_ (µg/ml) values of H77 sE1E2 scaffolds binding to bNAbs AR3C and AR4A. If absorbance at 450 nm is less than 0.5 at the starting and highest concentration (10 μg/ml), the antigen is considered to have negligible binding to the tested antibody, and the EC_50_ value is set to 10 μg/ml. For all in-house-produced sE1E2 scaffolds, oligomer (O) and dimer (D) species from SEC were tested.

Eleven designed sE1E2 scaffolds were transiently expressed in 500-ml HEK293F cells, purified by IMAC using a nickel column, and analyzed by size-exclusion chromatography (SEC) on a Superdex 200 column (**Fig. 1d**). sE1E2.LZ exhibited an SEC profile exhibiting an aggregation peak at ∼8.2 ml and two small tailing peaks (**Fig. 1d**, leftmost). All sE1E2.Cut_1_ scaffolds, except for sE1E2.Cut_1_.LZ, showed improved yield with a high-molecular weight (MW) peak at ∼10.6-10.8 ml and a potential dimer peak at ∼12.1-12.4 ml (**Fig. 1d**, top). Among the four leucine zippers, GCP produced the highest yield and the most prominent dimer peak. SPYΔN showed a similar SEC profile to GCP, with a more pronounced dimer peak, suggesting that dimeric sE1E2 was the predominant form with this covalently linked scaffold. Deletion of the pFP-containing E1 region (L264-F293) exerted a positive effect on expression yield and composition. As indicated by the SEC profiles, a higher ratio of dimer to high-MW species was observed for all five sE1E2.Cut_1+2_ scaffolds, with GCP and SPYΔN showing the most visible improvement (**Fig. 1d**, bottom). Among these five scaffolds, sE1E2.Cut_1+2_.SPYΔN was the best performer, with a prominent dimer peak, only a trace of high-MW species, and a relatively small aggregation peak. Next, sodium dodecyl sulfate-polyacrylamide gel electrophoresis (SDS-PAGE) was performed to analyze selected SEC fractions for ten sE1E2.Cut_n_ scaffolds, with two sE1E2.LZ controls included for comparison (**Fig. 1e**). The control antigen reported by Guest et al. (*76*), sE1E2.LZorg, registered separate E1 and E2 bands—but not an E1E2 band—on the reducing gel, with the non-reducing gel displaying a large diffuse band at the top (**Fig. 1e**, leftmost). In contrast, sE1E2.LZ showed distinct bands on the non-reducing gel corresponding to multiple species (**Fig. 1e**, left). For all in-house−designed sE1E2 scaffolds, SEC fractions at ∼12.1-12.4 ml produced a ∼100 kDa band, consistent with the MW of a single sE1E2 calculated from its amino acid sequence and *N*-linked glycans (∼3 kDa per glycan), whereas fractions at ∼10.6-10.8 ml corresponded to a band of 150-250 kDa, suggesting an oligomeric sE1E2 species. SDS-PAGE also revealed a scaffold-specific difference between the sE1E2.Cut_1_ and sE1E2.Cut_1+2_ constructs: for the four leucine zippers, dissociated E1 and E2 bands were discernible on the reducing gels of sE1E2.Cut_1+2_ scaffolds, whereas for SPYΔN, no E1 and E2 bands were observed regardless of the E1 loop deletion.

We used bNAbs AR3C and AR4A in an enzyme-linked immunosorbent assay (ELISA) to validate the H77 sE1E2 scaffolds (**Fig. 1f** and **Fig. S1b**). AR3C targets a major epitope within the E2 NF formed by part of the FL (426–443) and the tip of the CD81bl (529–531) (*55*), and represents a large family of AR3-directed human bNAbs of the V_H_1-69 origin (*40*). AR4A is an E1E2-specific bNAb (*32*) that recognizes an E2 epitope near the native E1−E2 interface (*70, 71*). AR4A has been used as an interface probe in binding assays to validate sE1E2 designs (*76, 79*) and as a chaperone in coexpression with E1E2 or sE1E2 to promote native-like E1−E2 association (*70, 71, 76*). In the ELISA, the control sE1E2 antigen designed by Guest et al. (*76*), sE1E2.LZorg, bound favorably to AR3C and AR4A with half-maximal effective concentration (EC_50_) values of 0.096 and 0.048 µg/ml, respectively, whereas the nickel-purified sE1E2.LZ-His_6_ showed poor AR3C binding and was barely recognized by AR4A (**Fig. 1f** and **Fig. S1b**). Among the ten designed sE1E2 scaffolds, the single sE1E2 form—expected from the cryo-EM structure of E1E2 and scaffold design—is the preferred species for vaccine development; other forms likely represent unintended assembly states (e.g., a dimer of heterodimers). Truncation of the E1 and E2 stems (Cut_1_) markedly improved bNAb binding for all five scaffolds except IAAL, with EC_50_ in the range of 0.072-0.148 for AR3C and 0.109-0.529 µg/ml for AR4A (**Fig. 1f** and **Fig. S1b**). Compared to sE1E2.LZorg (*76*), the sE1E2.Cut_1_ scaffolds showed similar AR3C binding affinities, but their AR4A binding was less effective, with a 2- to 10-fold higher EC_50_ value. Further deletion of the pFP-containing E1 loop (Cut_2_) dramatically increased bNAb recognition, with EC_50_ values of the resulting sE1E2.Cut_1+2_ scaffolds in the range of 0.010-0.027 for AR3C and 0.007-0.025 µg/ml for AR4A (**Fig. 1f** and **Fig. S1b**). Based on AR4A binding alone, sE1E2.Cut_1+2_.GCP-His_6_ was the best performer, with a 7-fold higher affinity than sE1E2.LZorg (*76*). sE1E2.Cut_1+2_.SPYΔN-His_6_ bound to AR3C and AR4A with similar affinities, 3.6- and 2.0-fold higher than sE1E2.LZorg (*76*), thus providing a balanced profile. For the majority of sE1E2.Cut_1_ and sE1E2.Cut_1+2_ scaffolds, the oligomeric sE1E2 form showed reduced bNAb binding, likely due to steric hindrance.

Our results revealed critical elements for designing a stable and native-like sE1E2 antigen. Truncation of the sE1 and sE2 stems, along with deletion of the unstructured pFP-containing E1 loop, produced an optimal sE1E2.Cut_1+2_ construct for scaffolding. The beneficial effect of deleting the E1 loop may be explained by its hydrophobicity (∼50%), which can cause aggregation or form non-specific epitopes, and by cysteines C272 and C281, which may form disulfide-linked sE1E2 oligomers with occluded bNAb epitopes. sE1E2.Cut_1+2_.GCP-His_6_ showed superior AR4A binding, suggesting that “archetypal” coiled coils of ∼30 aa may be effective scaffolds for presenting HCV sE1E2. Overall, sE1E2.Cut_1+2_.SPYΔN provides the most promising antigen in terms of expression yield, folding, stability, and epitope presentation.

### In vitro characterization of select HCV sE1E2.Cut_1+2_.SPY**Δ**N antigens

To advance an HCV vaccine construct to clinical trials, it is a prerequisite that diverse strains be screened to select an optimal vaccine backbone and that the antigen be produced using the standard manufacturing systems at scale (*99–101*). Previously, two sE1E2 scaffolds derived from genotype 1a H77 and 1b 1b09 were produced in Expi293F cells for in vitro evaluation, cryo-EM analysis, and animal immunization (*70, 76, 77*). In our recent studies, rationally designed vaccine candidates were produced in a transient Chinese hamster ovary cell (CHO) line, ExpiCHO, for in vitro and in vivo characterization (*67, 82, 83, 85–90*), which may inform future development using stable CHO cell lines under Good Manufacturing Practice (GMP) conditions. To facilitate the transition to the CHO system, the five H77 sE1E2.Cut_1+2_ scaffolds were transiently expressed in 25-ml ExpiCHO cultures and purified by IMAC using a nickel column. Unexpectedly, the SEC profiles revealed an indiscernible dimer peak, along with an overall low yield and the presence of lower-MW species (**Fig. S1c**), suggesting an incompatibility between the H77 strain and the CHO system.

To identify a suitable vaccine backbone, we extended the sE1E2.Cut_1+2_.SPYΔN construct design to genotype 1a HCV-1 and two strains from a panel of HCV E1E2 sequences: genotype 3 UKNP3.1.2 and genotype 5 UKNP5.2.1 (*102*) (**Fig. S2a**). Notably, HCV-1 was used to develop the first HCV E1E2 vaccine, which was GMP-produced in CHO cells and evaluated for safety and immunogenicity in a phase 1 trial (*94*). Briefly, the three sE1E2.Cut_1+2_.SPYΔN-His_6_ constructs were transiently expressed in either 500-ml HEK293F or 25-ml ExpiCHO cultures, followed by nickel purification and SEC analysis (**Fig. 2a**). All three strains showed higher overall yield than H77, with HCV-1 sE1E2.Cut_1+2_.SPYΔN-His_6_ being the best performer regardless of the cell line used. Multiple production runs of this HCV-1 sE1E2 scaffold showed overlapping dimer and high-MW peaks in the HEK293F-derived SEC profiles, but a consistent pattern of high yield and a prevalent dimer peak at 13.1-13.3 ml in the ExpiCHO-derived profiles. Reducing SDS-PAGE confirmed that the fractions at 13.1-13.3 ml indeed corresponded to the sE1E2 heterodimer, with a band between 75 and 100 kDa on the gel (**Fig. S2b**). Despite the lower expression yield, the UKNP5.2.1 sE1E2 scaffold produced a distinct dimer peak in the SEC profile, suggesting it is a suitable genotype 5 vaccine strain. Next, we applied differential scanning calorimetry (DSC) to assess the thermostability of the three sE1E2.Cut_1+2_.SPYΔN-His_6_ constructs expressed in two cell lines, for a total of six antigens (**Fig. 2b**). The thermostability of SpyTag/SpyCatcher has been studied in the context of a cyclized form with β-lactamase (BLA), with a melting temperature (T_m_) of 85.4 °C reported for the reconstituted SPY domain (*103*). Based on this study, the sharp peak at 91°C observed in all six thermograms likely corresponds to rapid unfolding of SPYΔN, whereas the broad peaks in the 50-65 °C range may be associated with melting of sE1E2, sE1, and sE2. We also observed a unique pattern for the HCV-1 sE1E2 scaffold produced in ExpiCHO cells: it generated two low-T_m_ peaks (at 50.1 and 58.7 °C), whereas the UKNP3.1.2 and UKNP5.2.1 sE1E2 scaffolds displayed a single low-T_m_ peak, suggesting strain-specific thermostability profiles.

**Fig. 2.**
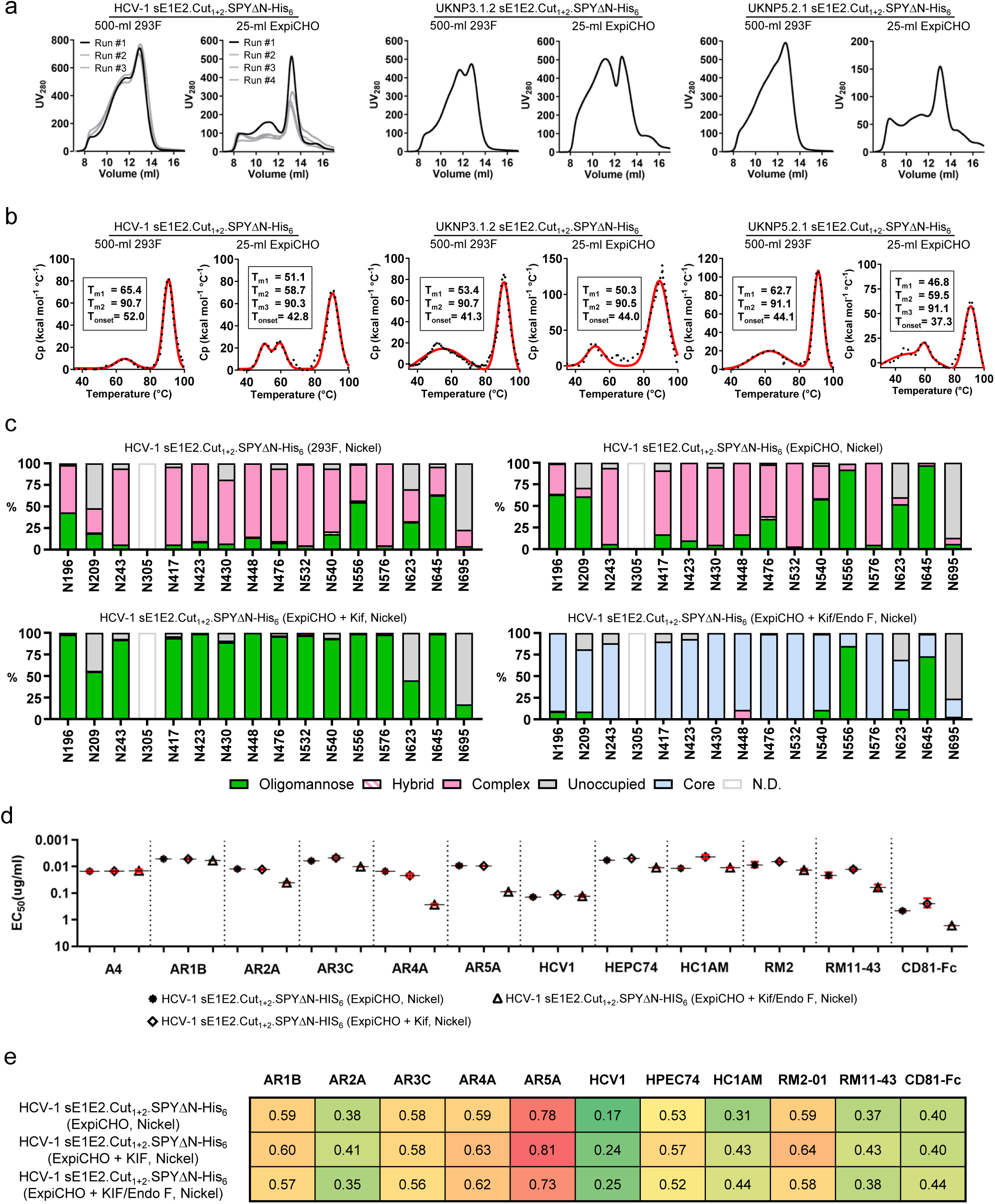
In vitro characterization of the HCV-1 sE1E2.Cut_1+2_.SPYΔN-His_6_ heterodimer. (**a**) SEC profiles of HEK293F- and ExpiCHO-produced genotype 1a HCV-1, genotype 3 UKNP3.1.2, and genotype 5 UKNP5.2.1 sE1E2.Cut_1+2_.SPYΔN-His_6_ antigens. All antigens were purified using a nickel column. Results are shown for three HEK293F and four ExpiCHO production runs. (**b**) DSC thermograms of HEK293F- and ExpiCHO-produced HCV-1, UKNP3.1.2, and UKNP5.2.1 sE1E2.Cut_1+2_.SPYΔN-His_6_ antigens. All antigens were purified using a nickel column and SEC. Experimental data and Gaussian fits are shown as black dots and red lines, respectively. Thermal denaturation midpoints (T_m_ or T_m1-3_) and onset temperatures (T_onset_) are labeled. (**c**) Site-specific glycan profiles for HCV-1 sE1E2.Cut_1+2_.SPYΔN-His_6_ antigens produced in two cell lines (HEK293F and ExpiCHO) and with two glycan modifications (Kif and Kif/endo F treatment). Kifunensine (Kif) was added to ExpiCHO cell cultures to convert N-glycans to oligomannose-type. Endo F1-3 then trimmed oligomannose-type glycans to GlcNAc cores. Glycan compositions are grouped into four types: complex (solid pink), hybrid (pink lines), oligomannose (solid green), and unoccupied (solid gray). Glycan sites that could not be determined are denoted “N.D.” (**d**) ELISA-derived EC_50_ (µg/ml) values of unmodified, Kif-treated, and glycan-trimmed HCV-1 sE1E2.Cut_1+2_.SPYΔN-His_6_ antigens binding to 11 NAbs and CD81-Fc. All antigens were produced in ExpiCHO cells and purified using a nickel column and SEC. If absorbance at 450 nm is less than 0.5 at the starting and highest concentration (10 μg/ml), the antigen is considered to have negligible binding, and the EC_50_ is set to 10 μg/ml. (**e**) BLI profiles of unmodified, Kif-treated, and glycan-trimmed HCV-1 sE1E2.Cut_1+2_.SPYΔN-His_6_ antigens binding to 10 NAbs (excluding the murine NAb A4) and CD81-Fc. Sensorgrams were obtained on an Octet RED96 using an antigen titration series of six concentrations (starting at 1000 nM, followed by 2-fold dilutions) and are shown in **Fig. S2g**. Peak responses at the highest concentration are displayed in a matrix, with cells colored from green (weak binding) to red (strong binding).

The HCV glycan shield consists of ∼15-16 glycans and masks conserved NAb epitopes on the E1E2 complex (*104, 105*). Site-specific glycan analysis has been performed for the bNAb-bound and unbound full-length AMS0232 E1E2 heterodimer using liquid chromatography-mass spectrometry (LC-MS) (*71*). Our recent studies have demonstrated the differential effects of glycan modification on NAb responses induced by HIV-1 Env and filovirus GP vaccines (*82, 85*). Here, we examined the HCV glycan shield on HCV-1 sE1E2.Cut_1+2_.SPYΔN-His_6_ produced in two cell lines and subjected to two glycan modifications. For the latter, kifunensine (Kif) was added to the ExpiCHO cell culture to enrich oligomannose content, and these *N*-glycans were further trimmed to GlcNAc stumps using endoglycosidase F1-3 (endo F1-3). Prior to glycan profiling, we assessed the quality of the glycan-modified antigens. In SEC analysis, Kif had minimal impact on sE1E2 folding, yield, or purity, whereas endo F shifted the dimer peak from ∼13.0 to ∼13.8 ml (**Fig. S2c**). In DSC analysis, both Kif- and Kif/endo F-treated antigens yielded similar thermograms, with T_m_ values nearly identical to the unmodified material (**Fig. S2d**). The double-peak (T_m1_/_Tm2_) pattern of sE1E2 unfolding appeared to be an intrinsic feature of this antigen—likely due to its strain and E1E2 sequence—when expressed in CHO cells (**Fig. 2B**, left and **Fig. S2d**). We next determined site-specific glycosylation and occupancy (**Fig. 2c** and **Fig. S2e**). Briefly, two aliquots of each antigen were digested separately with chymotrypsin and α-lytic protease to generate peptides and glycopeptides containing a single *N*-linked glycan site for LC-MS analysis. Both HEK293F- and ExpiCHO-produced sE1E2 displayed predominantly complex-type glycans, with a slightly higher abundance of oligomannose-type glycans in the ExpiCHO material. NeuGc and NeuAc glycans were absent in the ExpiCHO-produced antigen, but sialylated glycans were detected at every site in the HEK293F-produced sE1E2. In addition, high levels of fucosylation were observed in sE1E2 from both cell lines. As expected, Kif and Kif/endo F treatments transformed most *N*-glycan sites into oligomannose-typfe glycans and GlcNAc cores, respectively (**Fig. 2c**, bottom). Glycan N305, which forms a salt bridge with E655 in full-length E1E2 (*71*), could not be resolved by LC-MS analysis. Notably, Kif treatment creates the closest mimic of the native glycan shield on HCV virions, which is predominantly composed of oligomannose-type glycans (*71*).

HCV-1 sE1E2.Cut_1+2_.SPYΔN-His_6_ carrying unmodified, oligomannose-type (Kif-treated), and GlcNAc (Kif/endo F-treated) glycans was assessed against a panel of 11 NAbs, along with CD81 fused to the fragment crystallizable (Fc) region (CD81-Fc) (**Table S1**). These NAbs target diverse epitopes: one murine NAb binds an epitope on E1, while the others are directed against AR1-AR5 on E2 (*32, 65, 106*). ELISA revealed a differential effect of glycan modification on sE1E2 binding (**Fig. 2d** and **Fig. S2f**). Compared to the unmodified version, Kif treatment modestly improved sE1E2 binding to CD81-Fc and AR3-directed NAbs, with negligible impact on other epitopes. In contrast, Kif/endo F treatment resulted in varying levels of reduced binding to AR2-AR5. Glycan trimming likely affected NAb binding to the E2 NF and AR4 via weakened glycan interactions and local conformational changes, respectively. The E2 NF is surrounded by *N*-glycans N417, N423, N430, N532, and N540 (*59*). Crystal structures revealed that AR3C heavy and light chains (HC and LC) each form a hydrogen bond with the N430 GlcNAc core (*55, 67*), and HC1AM HC and LC interact with N423 and N532 GlcNAc cores through three hydrogen bonds (*66*). These intact GlcNAc cores largely mitigated the adverse effect of glycan trimming on AR3 recognition by the NAbs. AR4 is an E1E2-specific epitope near the noncanonical *N*-glycan N695, which contains a NXV motif with ∼25% occupancy (*71*). A hydrogen bond between N695 and AR4A was observed in the cryo-EM structure of a scaffolded sE1E2 (*70*), but not in the full-length E1E2 structure (*71*). Since removal of this glycan site had a minimal impact on E1E2 antigenicity (*71*), the 18-fold reduction in AR4A binding to endo F-treated HCV-1 sE1E2 is likely due to a conformational change at the E1-E2 interface. Results from biolayer interferometry (BLI) were consistent with the ELISA data (**Fig. 2e** and **Fig. S2g**). Overall, AR5A showed the highest binding signals with plateaued dissociation. Notably, glycan trimming caused some degree of abnormality in the BLI measurement at high antigen concentrations (**Fig. S2g**, left). Compared to the HEK293F-expressed H77 antigen (**Fig. 1f** and **Fig. S1b**), the ExpiCHO-expressed, unmodified or Kif-treated HCV-1 sE1E2.Cut_1+2_.SPYΔN-His_6_ bound to bNAbs AR3C and AR4A with up to 5.6- and 1.7-fold higher affinity, respectively, suggesting it is a suitable vaccine antigen.

HCV-1 sE1E2.Cut_1+2_.SPYΔN presented as a promising vaccine antigen. In addition to high yield and purity in CHO expression, it exhibited a desirable antigenic profile, with high affinities for AR4A, other NAbs targeting diverse epitopes on E1E2, and CD81. Its modest thermostability could be improved through stabilizing mutations. Most importantly, Kif treatment resulted in high oligomannose content—a key feature of the E1E2 glycan shield on native HCV virions (*71*). This Kif-treated construct warrants more detailed structural characterization.

### Electron microscopy (EM) analysis of HCV-1 sE1E2.Cut_1+2_.SPY**Δ**N bound to antibodies

While HCV E1E2 has resisted X-ray crystallography for decades, cryo-EM has enabled high-resolution structure determination of full-length E1E2 (*71*) and scaffolded sE1E2 (*70*), suggesting that cryo-EM may offer critical structural insight into this elusive antigen. We first conducted cryo-EM analysis of HCV-1 sE1E2.Cut_1+2_.SPYΔN-His_6_ in complex with the fragment antigen-binding (Fab) regions of AR4A and HEPC74. A total of 5,106 movies were collected on a Titan Krios 300 kV cryo-transmission electron microscope (cryo-TEM) equipped with a Gatan K3 camera. After movie processing in cryoSPARC (*107*), populations of sE1E2.Cut_1+2_.SPYΔN-His_6_ bound by both antibodies were observed in the dataset confirming sE1E2.Cut_1+2_.SPYΔN-His_6_ was structurally compatible for recognition by the two bNAbs. Selection of 36,896 particles for 3D classification enabled subsequent reconstruction of a density map to a modest resolution of 6.30 Å (**Fig. S3a** and **Table S2**). Rigid-body fitting of the NAb-bound sE1E2.SZ structure (PDB ID: 8FSJ (*70*)) into the density revealed a similar conformation, despite differences in the E1E2 strain and construct (1b09 sE1E2 *vs.* HCV-1 sE1E2.Cut_1+2_) and scaffold (SZ vs. SPYΔN) (**Fig. S3b**). We then performed cryo-EM analysis of HCV-1 sE1E2.Cut_1+2_.SPYΔN-His_6_ bound to AR4A and AR3C Fabs. Two different grid preparation platforms were used in sample vitrification to mitigate orientation bias (see Methods), resulting in two datasets with 19,080 and 5,248 movies. After cryoSPARC processing (*107*), two subsets (85,959 and 89,013 particles) were combined to generate a density map at a resolution of 4.97 Å (**Fig. S3c** and **Table S2**). This map enabled rigid-body fitting of the cryo-EM structure of sE1E2.SZ with AR4A (PDB ID: 8FSJ (*70*)) and the crystal structure of E2c with AR3C (PDB ID: 4MWF (*55*)), providing further structural confirmation of the antigenic integrity sE1E2.Cut_1+2_.SPYΔN-His_6_ (**Fig. S3d**). However, the resolutions obtained from current cryo-EM analysis prevented atomic-level modeling of either complex.

Recently, we applied nsEM to obtain low-resolution (10-12 Å) models of NAb-antigen complexes to validate rationally designed viral antigens such as HIV-1 Env (*82*), RSV F (*88*), and ebolavirus GP (*85*), and to identify NAb epitopes (*88*). Here, we adapted this approach as a low-resolution alternative for antigen model building and NAb epitope mapping, using HCV NAbs with known complex structures as anchors in density fitting to identify both the position of the scaffold (e.g., SPYΔN) and the binding site of an NAb on the scaffolded sE1E2 structure. We first performed nsEM analysis of HCV-1 sE1E2.Cut_1+2_.SPYΔN-His_6_ bound to AR3C and AR4A Fabs individually and in combination. In each case, the complex was imaged on a Talos L120C TEM and processed with cryoSPARC to generate density maps suitable for rigid body fitting (**Fig. S3e**). For AR3C, we generated a model complex by superimposing the crystal structure of the E2mc3-v1/ARC3 complex (PDB ID: 6UYD (*67*)) onto the cryo-EM structure of a scaffolded sE1E2 (PDB ID: 8FSJ (*70*)). Upon fitting this model into the density map, we identified unoccupied density extending from the sE1E2 C-termini (E1-H312 and E2-Y701), into which the SPY domain structure (PDB ID: 4MLI (*92*)) fit well (**Fig. 3a**, left). This “complete” low-resolution complex confirmed that AR3C binds to the FL and CD81bl, consistent with high-resolution cryo-EM data (*70*). Similarly, fitting the AR4A-bound sE1E2.SZ structures (PDB ID: 8FSJ (*70*)) and the SPY domain (PDB ID: 4MLI (*92*)) into the nsEM density produced a complete model for the AR4A-bound HCV-1 sE1E2.Cut_1+2_.SPYΔN-His_6_ complex (**Fig. 3a**, middle), confirming that AR4A binds to the E1-E2 bridging region in E2 (residues 646–701) (*70*). Fitting both AR3C- and AR4A-bound HCV-1 sE1E2.Cut_1+2_.SPYΔN-His_6_ models into the density map of the ternary complex revealed nearly identical epitopes and angles of approach for the two Fabs, consistent with our cryo-EM analysis (**Fig. 3a**, right; **Fig. S3c**). These structural findings highlight the utility of HCV-1 sE1E2.Cut_1+2_.SPYΔN-His_6_ in nsEM-based NAb epitope mapping.

**Fig. 3.**
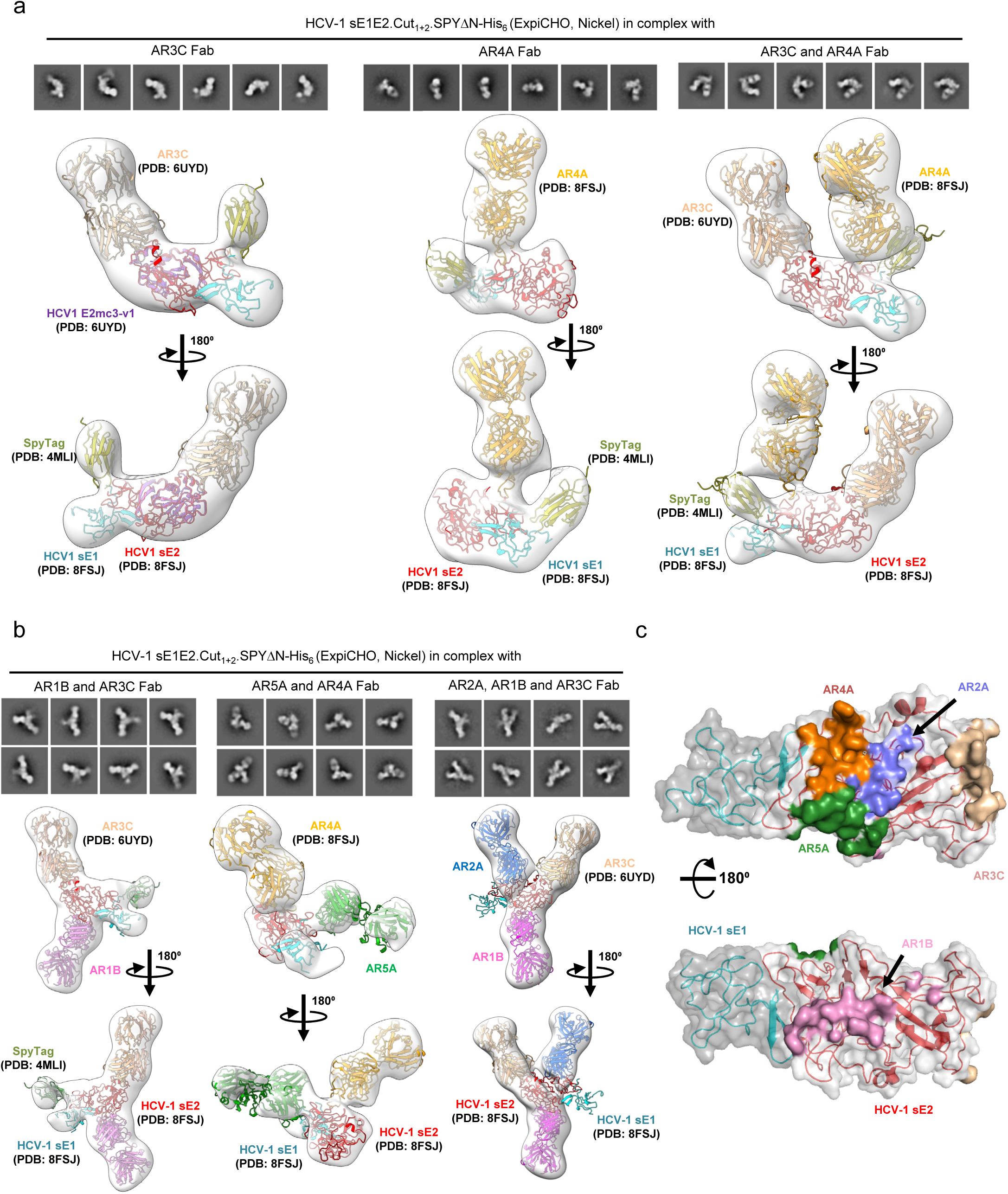
Negative-stain EM analysis of HCV-1 sE1E2.Cut_1+2_.SPYΔN-His_6_ bound to NAbs. (**a**) Selected 2D class averages (top) and EM density maps from 3D reconstructions (bottom) of the sE1E2 antigen in complex with AR3C (left), AR4A (middle), or both AR3C and AR4A (right). Atomic structures were fitted into the corresponding density maps to facilitate interpretation. (**b**) Selected 2D class averages (top) and 3D EM density maps (bottom) of sE1E2 bound to different NAb combinations: AR1B and AR3C (left), AR5A and AR4A (middle), and AR2A, AR1B, and AR3C (right). In (**a**-**b**), the maps and fitted models were rotated 180° to provide alternative viewing angles for the sE1E2/NAb interfaces. (**c**) Epitope mapping of antigenic regions AR1–5 on the surface of HCV-1 sE1E2.Cut1+2.SPYΔN-His6. The locations of AR1, AR2, and AR5 epitopes were inferred from negative-stain EM reconstructions of sE1E2 in complex with AR1B, AR2A, and AR5A, respectively. In (**a**-**c**), structural models were fitted into the EM densities using the cryo-EM structure of engineered sE1E2.SZ bound to HEPC74 and AR4A (PDB ID: 8FSJ), the crystal structure of the E2mc3/AR3C complex (PDB ID: 6UYD), and the crystal structure of the SpyTag/SpyCatcher system (PDB ID: 4MLI).

We then extended this approach to AR1B, AR2A, and AR5A, whose epitopes have not been structurally defined on the native E1E2 heterodimer. Previous studies suggested the AR1B and AR2A epitopes lie on E2, but their precise locations remain unclear (*61*). Using AR3C and/or AR4A Fabs as anchors, we fitted AR1B, AR2A, and AR5A Fab models into unoccupied regions within nsEM density maps generated for HCV-1 sE1E2.Cut_1+2_.SPYΔN-His_6_ bound to various NAb combinations. These model complexes revealed the potential epitope of each NAb on E1E2 (**Figs. 3b, c**, and **Table S3**). AR1B appeared to recognize a loop between β6 and β7, opposite the AR3C site, with 12 of 21 residues identified by mutagenesis studies (*33, 106*) located within the nsEM-defined epitope. AR5A appeared to target a region near the E1-E2 interface, centered around residues R639 and L665, extending toward the back layer (BL) of the E1E2 heterodimer. Notably, R639 and L665 were also identified in previous mutagenesis analyses (*32, 106*). This model complex suggested that the AR5A binding site was spatially distinct from that of AR4A, allowing both NAbs to engage E1E2 simultaneously without competition. However, the presence of one NAb may influence the angle of approach and E2 interactions of the other, especially given the proximity of the AR4A and AR5A binding sites. This finding differed from previous mutagenesis analyses (*32, 33, 106*), suggesting that unrelated mutations causing conformational change may result in the loss of NAb binding. Lastly, a quaternary complex with AR1B, AR3C, and AR2A revealed that AR2A binds the β11 strand bordering the AR4A and AR5A footprints, with 88% of the residues overlapping with the mutagenesis-defined epitope (*33, 106*). The nsEM-based epitope mapping thus provided valuable insights into how NAbs recognize ARs on native E1E2.

Our nsEM analysis, supported by cryo-EM, revealed key structural and antigenic features of a rationally designed sE1E2 antigen. This SPYΔN-scaffolded sE1E2, which presents all five major ARs in a stable, native-like conformation, enables rapid epitope mapping via nsEM for antibodies isolated from infected individuals and immunized animals. The test cases of AR1B, AR2A, and AR5A serve as templates for nsEM-based NAb epitope mapping. However, this sE1E2 scaffold cannot be used to probe stem-directed NAbs, such as IGH526 (*37, 49, 71*).

### Rational design and in vitro characterization of HCV-1 sE1E2-presenting nanoparticles

Several strategies have been explored to develop HCV Env-based NP vaccines (*75*). Previously, we designed a mini-E2 core, termed E2mc3, for strains from genotypes 1a and 6a, and displayed E2mc3 on a ferritin (FR) 24-mer as well as unmodified E2p and I3-01 60-mers (*67*). The E2mc3-E2p 60-mer elicited potent NAb responses with a unique B cell repertoire profile (*67*). Yan et al. reported an sE2-FR NP with improved immunogenicity in mice (*68*). Sliepen et al. designed an sE2E1 construct via permutation and displayed six different sE2E1 antigens on a two-component NP (2c-NP) platform, I53 50, for rabbit immunization (*79*). Although this mosaic sE2E1 2c-NP induced cross-NAbs, the trimeric unit sE2E1-I53-50A failed to bind bNAbs AR4A (*32*) and AT1618 (*108*), suggesting a non-native conformation (*79*). Most recently, an engineered sE1E2.SZ heterodimer (*70*), the HCV core protein, and a Toll-like receptor (TLR) 7/8 agonist were co-assembled with biodegradable polymers into nanocomplexes (*78*). While this vaccine was highly immunogenic in mice, cryo-EM revealed an irregular shape due to random polymer assembly (*78*). Thus, while these nanocomplexes approximate the size of a virion, they do not adhere to the regular ordered assembly that typifies a native virus. To date, a protein NP vaccine presenting “native-like” sE1E2 antigens has not yet been reported.

We aimed to develop such an sE1E2 NP vaccine by integrating sE1E2.Cut_1+2_.SPYΔN into FR and our multilayered 1c-SApNP platform (*82, 86, 87, 90*). Based on the nsEM analysis (**Fig. 3**), sE1E2.Cut_1+2_.SPYΔN adopts an elongated shape, with sE1Δ264-293 and sE2 tightly anchored to SPYΔN, which can be displayed on an SApNP by fusing the C-terminus of SpyCatcherΔN to the N-terminus of an NP subunit (**Fig 4a**, left). We tested the FR 24-mer and I3-01v9a 60-mer (*90*) as candidate SApNP carriers. Notably, I3-01v9a has a multilayered construct design, with an inner layer formed by 60 locking domains (LDs) and a hydrophobic T-help core composed of 60 PADRE peptides (*82, 86, 87*), and has been optimized for displaying monomeric antigens (*90*). E2p, which has tightly clustered N-termini at each threefold axis on the NP surface, is better suited for trimer display and was therefore excluded from this study. To accommodate the non-symmetric sE1E2 shape and enhance epitope exposure, we inserted 10GS and 5GS spacers between SPYΔN and the NP subunit for FR and I3-01v9a-LD7-PADRE (or I3-01v9a-L7P (*90*)), respectively (**Fig. S4a**). Computational modeling estimated particle diameters of 36.9 and 47.0 nm for FR and I3-01v9a respectively, measured at S432 in the E2 NF (**Fig. 4a**, middle and right).

**Fig. 4.**
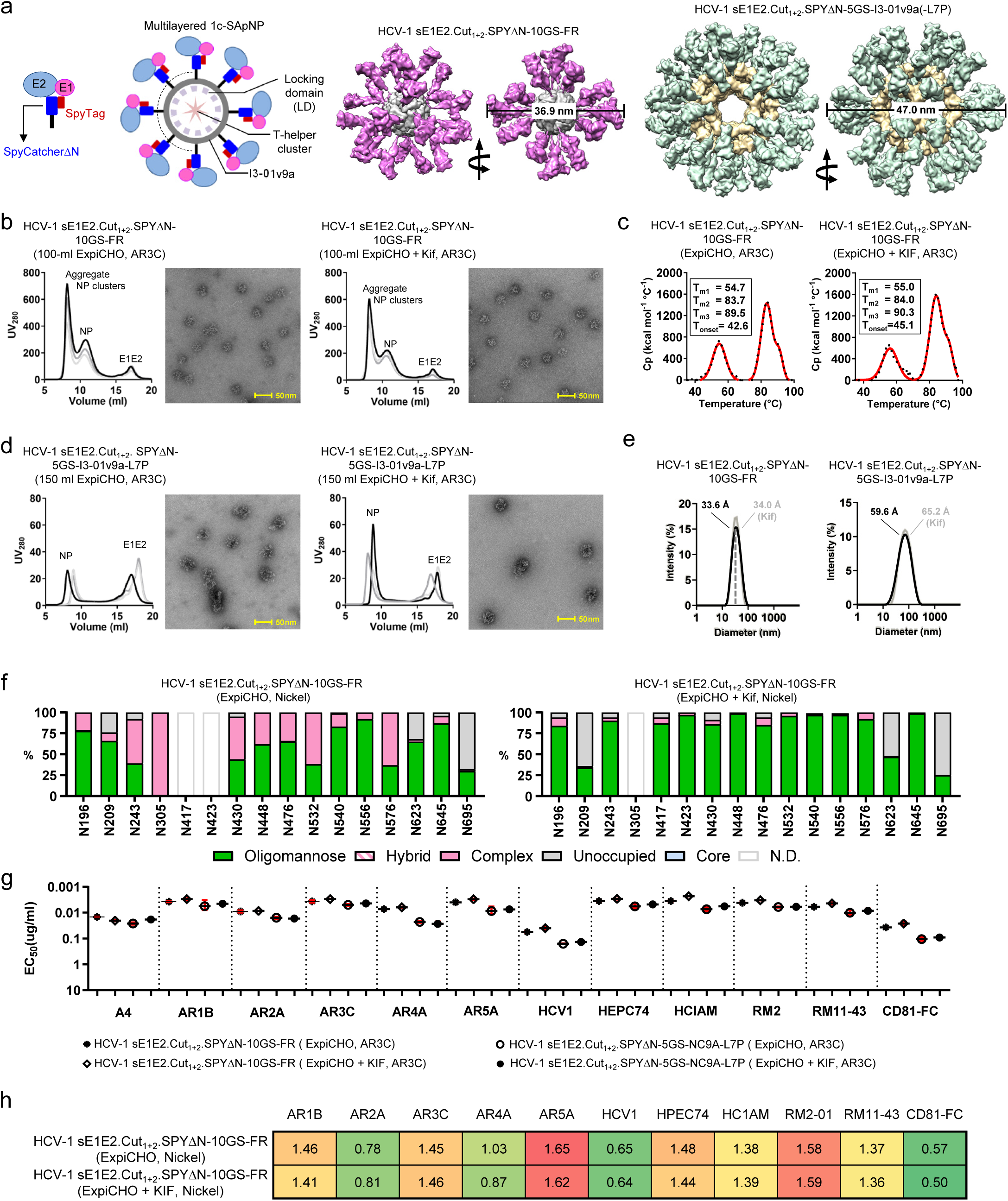
Rational design and in vitro characterization of HCV-1 sE1E2-presenting protein nanoparticles. (**a**) Left: schematic representation of sE1E2.Cut_1+2_.SPYΔN and its multivalent display on a multilayered self-assembling protein nanoparticle (SApNP). Middle: surface model of HCV-1 sE1E2.Cut_1+2_.SPYΔN-10GS-FR, with a diameter of 36.9 nm measured at E2-S432. Right: surface model of HCV-1 sE1E2.Cut_1+2_.SPYΔN-5GS-I3-01v9a-LD7-PADRE (or I3-01v9a-L7P), with a diameter of 47.0 nm. (**b**) SEC profiles of ExpiCHO-produced, AR3C-purified HCV-1 sE1E2.Cut_1+2_.SPYΔN-10GS-FR with unmodified (left) and Kif-treated (right) glycans. An EM micrograph corresponding to the ∼10.6 ml SEC peak is shown beside the chromatogram. Three production runs were used to generate the SEC data. (**c**) DSC thermograms of ExpiCHO-produced, AR3C-purified HCV-1 sE1E2.Cut_1+2_.SPYΔN-10GS-FR with unmodified (left) and Kif-treated (right) glycans. Experimental data and Gaussian fits are shown as black dots and red lines, respectively. Thermal denaturation midpoints 1-3 (T_m1-3_) and onset temperatures (T_onset_) are labeled. (**d**) SEC profiles of ExpiCHO-produced, AR3C-purified HCV-1 sE1E2.Cut_1+2_.SPYΔN-5GS-I3-01v9a-L7P with unmodified (left) and Kif-treated (right) glycans. An EM micrograph corresponding to the ∼8-9 ml SEC peak is shown. Two production runs were used to generate the SEC data. (**e**) Particle size distributions of HCV-1 sE1E2.Cut_1+2_.SPYΔN-presenting FR (left) and I3-01v9a-L7P (right), each with unmodified and Kif-treated glycans. Hydrodynamic diameter (D_h_) was measured by DLS and labeled on the plots. (**f**) Site-specific glycan profiles for unmodified (left) and Kif-treated (right) HCV-1 sE1E2.Cut_1+2_.SPYΔN-10GS-FR. Kifunensine (Kif) was added to ExpiCHO cultures during FR expression to convert N-glycans to oligomannose-type. Glycan compositions are grouped into four types: complex (solid pink), hybrid (pink lines), oligomannose (solid green), and unoccupied (solid gray). Glycan sites that could not be determined are denoted “N.D.” (**g**) ELISA-derived EC_50_ (µg/ml) values of unmodified (left) and Kif-treated (right) HCV-1 sE1E2.Cut_1+2_.SPYΔN-presenting FR and I3-01v9a binding to 11 NAbs and CD81-Fc. All SApNPs were produced in ExpiCHO cells and purified using an AR3C column and SEC. If absorbance at 450 nm is less than 0.5 at the starting and highest concentration (10 μg/ml), the antigen is considered to have negligible binding, and the EC_50_ value is set to 10 μg/ml. (**h**) BLI profiles of unmodified and Kif-treated HCV-1 sE1E2.Cut_1+2_.SPYΔN-10GS-FR SApNPs binding to 10 NAbs (excluding the murine NAb A4) and CD81-Fc. Sensorgrams were obtained on an Octet RED96 using an antigen titration series of six concentrations (starting at 18 nM, followed by 2-fold dilutions) and are shown in **Fig. S4f**. Peak values at the highest concentration are displayed in a matrix, with cells colored from green (weak binding) to red (strong binding).

HCV-1 sE1E2.Cut_1+2_.SPYΔN-10GS-FR was transiently expressed in 100-ml ExpiCHO cells with or without Kif treatment. Using a previously described immunoaffinity chromatography (IAC) method (*85, 87, 90, 109*), we prepared an AR3C antibody column to purify HCV-1 sE1E2-presenting SApNPs. The AR3C-purified protein from three production runs was analyzed by SEC on a Superose 6 10/300 column, with fractions of interest further examined by nsEM (**Fig. 4b**). Regardless of the Kif treatment or production run, all SEC profiles showed three distinct peaks at ∼8.1, ∼10.6, and ∼17.0 ml, likely corresponding to aggregates, NPs, and unassembled subunits, respectively. nsEM analysis confirmed that the middle peak contained well-formed NPs with a recognizable FR core and surface decorations. Additional analyses were conducted to characterize the ∼8.1 ml peak relative to the NP peak at ∼10.6 ml (**Fig. S4b**). Using either AR3C or AR4A (*32*) in IAC purification, large NP aggregates or clusters were observed in the high-MW peak. DSC was used to assess the thermostability of unmodified and Kif-treated FR SApNPs (**Fig. 4c**). Consistent with the DSC data for individual sE1E2 antigens (**Fig. 2b**), the first peak represented sE1E2 unfolding (T_m1_ = ∼55 °C), while the second and third peaks reflected melting of FR (T_m2_ = ∼84 °C) and SPYΔN (T_m3_ = ∼90 °C), respectively. HCV-1 sE1E2.Cut_1+2_.SPYΔN-5GS-I3-01v9a-L7P was similarly analyzed by SEC and nsEM after transient expression in 150-ml ExpiCHO cells, with and without Kif treatment, and AR3C purification (**Fig. 4d**). The yield of I3-01v9a was ∼15-20-fold lower than FR across three production runs, likely due to the complexity of simultaneously assembling 60 native-like sE1E2 heterodimers on the NP surface along with an inner LD layer and a PADRE core. EM confirmed that the SEC peak at ∼8-9 ml contained large NPs with densely arrayed sE1E2 antigens. Due to the low yield, DSC was not performed for I3-01v9a. Reducing SDS-PAGE and blue native (BN) PAGE confirmed the MW of subunits and the purity of intact NPs, respectively (**Fig. S4c**). Both SApNPs showed bands at ∼100-125 kDa on reducing gels, with the Kif-treated material running slightly higher. On the BN gel, FR produced a band above the 669 kDa marker, while I3-01v9a did not migrate from the well due to its large size. Particle size distributions for unmodified and Kif-treated FR and I3-01v9a were determined by dynamic light scattering (DLS) using a Zetasizer (**Fig. 4e**). Based on hydrodynamic diameter (Dh), FR averaged 33-34 nm, whereas I3-01v9a showed broader distributions centered at 59.6 and 65.2 nm for unmodified and Kif-treated SApNPs, respectively. The multilayered I3-01v9a 60-mer appeared more expanded in solution compared to the tightly packed FR 24-mer.

Site-specific glycan glycosylation and occupancy were determined for unmodified and Kif-treated HCV-1 sE1E2.Cut_1+2_.SPYΔN-10GS-FR using the same protocol as for the individual sE1E2 antigen (**Fig. 4f** and **Fig. S4d**). Compared to individual sE1E2 (**Fig. 2c**), the unmodified sE1E2 FR displayed more oligomannose-type glycans on the NP surface, although complex-type glycans still predominated (**Fig. 4f**). The preferential binding of AR3C to Kif-treated sE1E2 (**Fig. 2d**) likely enriched oligomannose-type glycans in sE1E2 FR purified via the AR3C column. Also consistent with individual sE1E2 (**Fig. 2c**), Kif treatment effectively inhibited N-glycan processing on sE1E2 FR, inducing primarily GlcNAc_2_Man_8_ and GlcNAc_2_Man_9_ glycoforms at each site, with minimal impact on site occupancy. The antigenicity of HCV-1 sE1E2.Cut_1+2_.SPYΔN-presenting FR and I3-01v9a SApNPs was assessed by ELISA (**Fig. 4g** and **Fig. S4e**). Compared to the individual sE1E2, particulate display improved NAb binding by up to 3.6- and 3.7-fold and CD81-Fc binding by 12.4- and 9.5-fold for unmodified and Kif-treated sE1E2 FR, respectively. Notably, Kif-treated sE1E2 FR bound the E1E2-specific AR4A and AR5A bNAbs with 3.5- and 3.2-fold higher affinities, respectively, than the Kif-treated sE1E2 alone. However, I3-01v9a appeared less effective in ligand binding than FR and, in ∼40% of cases, even less effective than individual sE1E2. Due to the low yield of I3-01v9a, only FR was assessed by BLI (**Fig. 4H** and **Fig. S4f**). The FR 24-mer showed the greatest improvement in binding to AR5A and RM2-01 compared to individual sE1E2, with 2.0-2.7-fold higher signals. Similarly, I3-01v9a yielded slightly lower peak binding signals than FR for most tested ligands, consistent with the ELISA results.

Our results indicate that sE1E2.Cut_1+2_.SPYΔN is optimal for designing HCV NP vaccines, as the covalent SPYΔN scaffold transforms sE1E2 from a metastable heterodimer into a stable monomer that can be readily displayed on protein NPs. Based on the current construct design, the FR 24-mer with a 10GS spacer exhibited the most desirable in vitro properties, whereas the I3-01v9a 60-mer with a 5GS linker may require further optimization to improve production yield and surface display. Nonetheless, HCV-1 sE1E2.Cut_1+2_.SPYΔN-presenting FR and I3-01v9a SApNPs can provide valuable insight into NP vaccine-induced immune responses.

### HCV-1 sE1E2.Cut_1+2_.SPYΔN heterodimer and NP vaccine-induced NAb responses in mice

We assessed the immunogenicity of ExpiCHO-produced HCV-1 sE1E2.Cut_1+2_.SPYΔN vaccines, including one His_6_-tagged dimer and two SApNPs (FR and I3-01v9a-L7P), in mice (**Fig. 5a**). The antigens —either nickel/SEC-purified dimer or AR3C/SEC-purified SApNPs—were adjuvanted with aluminum hydroxide (AH) and a toll-like receptor 9 (TLR9) agonist (CpG ODN 1826) to enhance vaccine-induced immune responses. Mice received four immunizations at weeks 0, 3, 6, and 9 via the intraperitoneal (i.p.) route. Each dose contained 10 μg of antigen formulated with 100 μl of CpG/AH adjuvant, for a final injection volume of 200 μl. Sera were collected 2 weeks after each immunization for HCV pseudoparticle (HCVpp) neutralization assays (*67*), and 50% inhibitory dilution (ID_50_) values were determined to evaluate serum NAb responses. Week-0 sera from all groups showed no background activity against genotype-1a H77 HCVpp, and no NAb response was detected at week 2. However, ID_50_ titers against H77 HCVpp were observed at weeks 5, 8, and 11 (**Fig. 5b** and **Fig. S5a-b**). At week 5, FR and I3-01v9a SApNPs showed ID_50_ titers of 817 and 670, respectively, which were 45.4- and 37.2-fold higher than the dimer. At week 8, NAb responses in the FR and I3-01v9a groups peaked at ID_50_ titers of 1253 and 1251, respectively, after three doses. In contrast, the dimer group peaked after four doses, reaching an ID_50_ titer of 589 at week 11, which remained lower than both SApNP groups. Week-11 sera were also tested against heterologous 3.1.2, 4.2.2, ED43, and 5.2.1 HCVpps (**Fig. 5c** and **Fig. S5c-d**). While none of the vaccines elicited NAbs against 3.1.2, the I3-01v9a 60-mer induced the highest ID_50_ titers of 70, 145, and 1030 against 4.2.2, ED43, and 5.2.1, which were 1.9-, 2.8-, and 17.4-fold higher than the sE1E2 dimer. These results demonstrate the benefit of NP display of a stable, native-like sE1E2 antigen in enhancing NAb responses. Notably, despite the suboptimal antigenicity of the I3-01v9a 60-mer (**Fig. 4g, h**), its large NP size (∼50 nm) contributed to the elicitation of cross-NAbs.

**Fig. 5.**
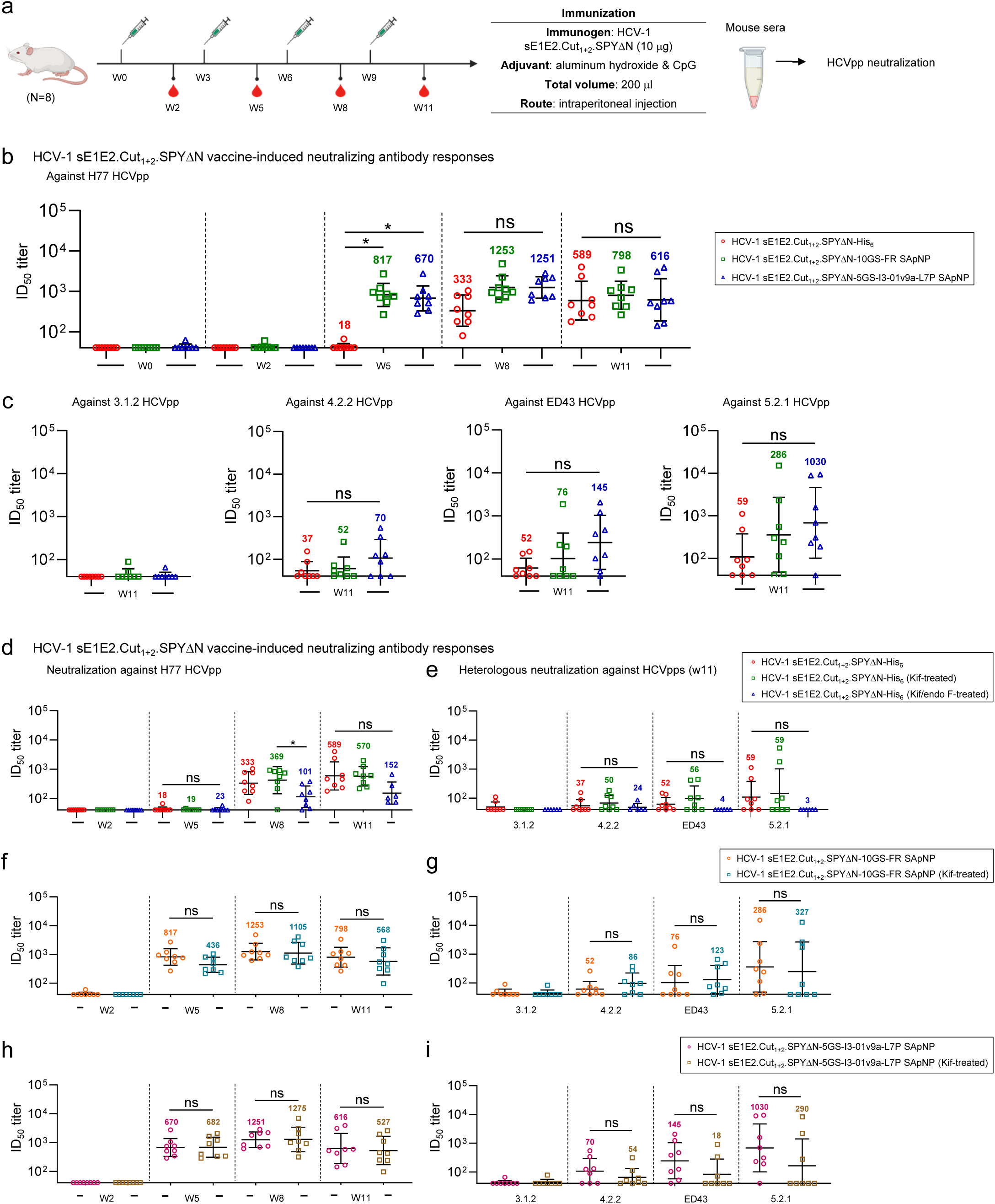
Antibody responses to rationally designed HCV-1 sE1E2.Cut_1+2_.SPYΔN heterodimer and SApNP vaccines in mice. (**a**) Schematic illustration of the mouse immunization regimen for various HCV-1 sE1E2.Cut_1+2_.SPYΔN vaccines (n = 8 mice/group). The administered dose was 200 μl of antigen/AH + CpG adjuvant mix containing 10 μg of immunogen and 100 μl of adjuvant. Mice were immunized via the intraperitoneal (i.p.) route at weeks 0, 3, 6, and 9. (**b**) Longitudinal serum NAb responses induced by HCV-1 sE1E2, FR, and I3-01v9a vaccines against genotype 1a H77 HCVpp. Serum samples from weeks 0, 2, 5, 8, and 11 were tested in neutralization assays to determine the 50% inhibitory dilution (ID_50_) titers. (**c**) Endpoint NAb responses induced by HCV-1 sE1E2, FR, and I3-01v9a vaccines against genotype 3 UKNP3.1.2, genotype 4 UKNP4.2.2 and ED43, and genotype 5 UKNP5.2.1 HCVpps. Serum from week 11 was tested in neutralization assays. (**d**-**e**) NAb responses induced by unmodified, Kif-treated, and Kif/endo F-treated HCV-1 sE1E2.Cut_1+2_.SPYΔN dimer vaccines (produced in ExpiCHO cells) against (**d**) H77 HCVpp at weeks 2, 5, 8, and 11, and (**e**) UKNP3.1.2, UKNP4.2.2, ED43, and UKNP5.2.1 HCVpps at week 11. (**f**-**g**) NAb responses induced by unmodified and Kif-treated HCV-1 sE1E2.Cut_1+2_.SPYΔN-10GS-FR SApNP vaccines against (**f**) H77 HCVpp at weeks 2, 5, 8, and 11, and (**g**) UKNP3.1.2, UKNP4.2.2, ED43, and UKNP5.2.1 HCVpps at week 11. (**h**-**i**) NAb responses induced by unmodified and Kif-treated HCV-1 sE1E2.Cut_1+2_.SPYΔN-5GS-I3-01v9a-L7P SApNP vaccines against (**h**) H77 HCVpp at weeks 2, 5, 8, and 11, and (**i**) UKNP3.1.2, UKNP4.2.2, ED43, and UKNP5.2.1 HCVpps at week 11. In (**b**-**i**), all ID_50_ titers were calculated from HCVpp neutralization assays, with geometric means labeled on the plots. ID_50_ values were derived by setting % neutralization to 0.0-100.0. Data points in the neutralization plots are shown as geometric mean ± geometric SD. Data were analyzed using one-way ANOVA followed by Tukey’s *post hoc* test for each time point. Two-tailed unpaired *t*-tests were used to compare geometric means of two independent groups. Statistical significance is indicated as follows: ns (not significant) and \**p* < 0.05. The mouse immunization schematic was created with BioRender.com.

Next, we assessed the immunogenicity of HCV-1 sE1E2.Cut_1+2_.SPYΔN vaccines carrying oligomannose-type (Kif-treated) and GlcNAc core (Kif/endo F-treated) glycans in mice. We first evaluated NAb responses induced by the sE1E2 dimer vaccines against H77 HCVpp (**Fig. 5d** and **Fig. S5e-f**). None of the dimer groups showed a NAb response at week 2, but all three groups exhibited detectable ID_50_ titers at weeks 5, 8, and 11. Among them, the glycan-trimmed dimer elicited the lowest ID_50_ titers at weeks 8 and 11. The unmodified and Kif-treated dimers produced comparable ID_50_ values of 570-589 at week 11, which were 3.8-3.9-fold higher than those of the glycan-trimmed dimer. Week-11 sera were also assessed against heterologous 3.1.2, 4.2.2, ED43, and 5.2.1 HCVpps (**Fig. 5e** and **Fig. S5g-h**). As with H77 neutralization (**Fig. 5d**), the unmodified and Kif-treated dimers elicited higher ID_50_ titers against 4.2.2, ED43, and 5.2.1 than the glycan-trimmed dimer, although differences were not statistically significant. For both FR and I3-01v9a SApNPs, unmodified and Kif-treated groups yielded comparable ID_50_ titers across all timepoints and all five HCVpps tested in this mouse study (**Fig. 5f-i** and **Fig. S5i-l**). Both unmodified and Kif-treated SApNP groups peaked in NAb responses against H77 HCVpp after three doses, with ID_50_ values ranging from 1105 to 1275 (**Fig. 5f, h**). Neutralization assays revealed measurable NAb titers in these SApNP groups against 4.2.2, ED43, and 5.2.1 HCVpps, consistently higher than those of the corresponding dimer groups (**Fig. 5e**). However, the Kif-treated I3-01v9a SApNP yielded lower ID_50_ titers than expected based on its unmodified counterpart and the trend observed for the FR groups (**Fig. 5i**). Overall, enrichment of oligomannose content did not impair sE1E2-induced NAb responses relative to the unmodified glycans.

### Effect of experimental factors on sE1E2 dimer and SApNP-induced NAb responses

We examined the potential effects of the expression system, sex-specific differences, and injection route on NAb responses induced by HCV-1 sE1E2.Cut_1+2_.SPYΔN vaccines. The protocol used in our earlier immunizations (**Fig. 5a**) was adopted to test these experimental variables (**Fig. 6a**).

**Fig. 6.**
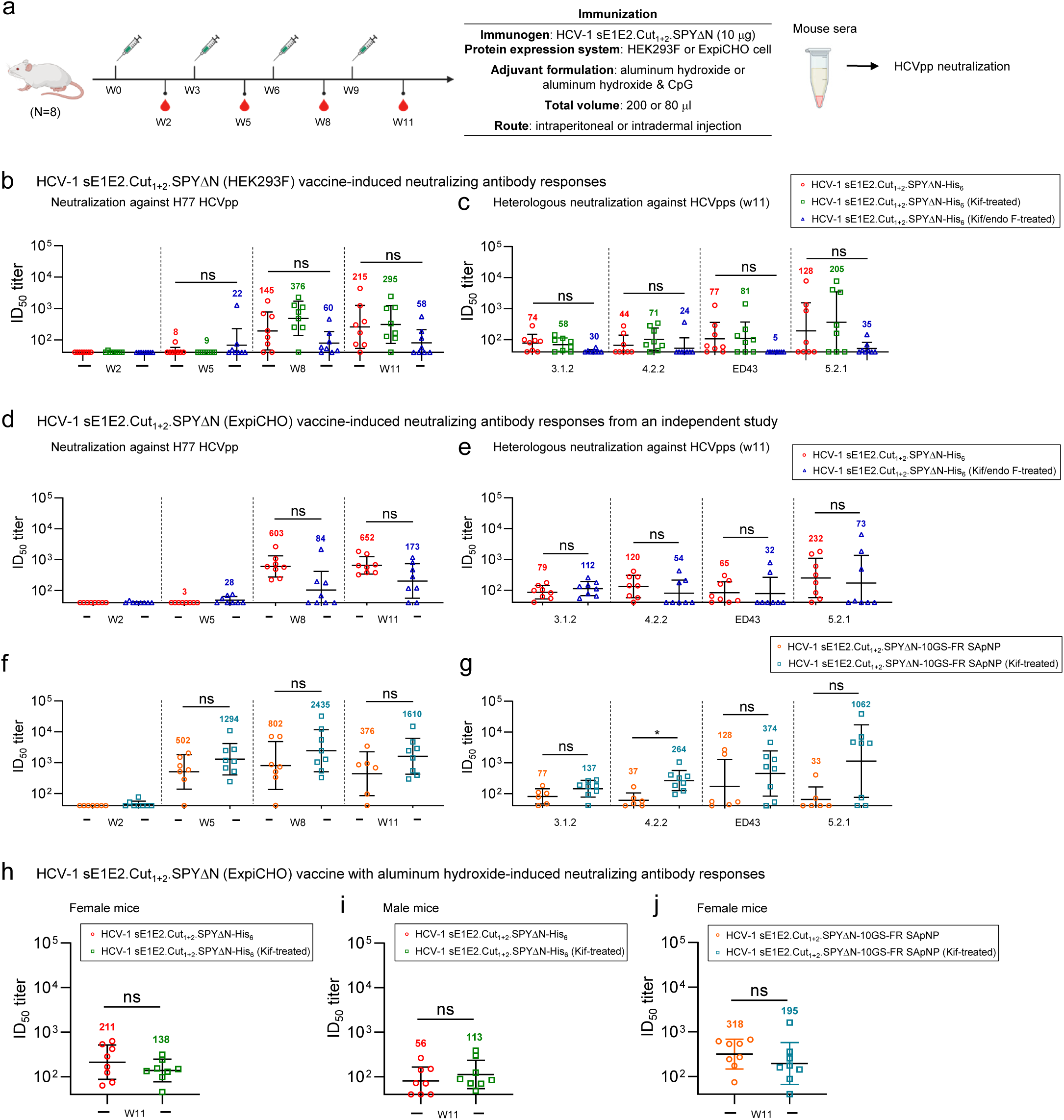
Effect of expression system, sex-specific differences, and injection route on antibody responses elicited by various HCV-1 sE1E2.Cut_1+2_.SPYΔN vaccines. (**a**) Schematic illustration of the mouse immunization regimen for HCV-1 sE1E2.Cut_1+2_.SPYΔN heterodimer and SApNP vaccines (n = 8 mice/group). The administered dose was 200 μl of antigen/AH + CpG adjuvant mix containing 10 μg of antigen and 100 μl of adjuvant via the intraperitoneal (i.p.) route, or 80 μl of antigen/AH adjuvant mix containing 10 μg of antigen and 40 μl of adjuvant via the intradermal (i.d.) route through injection into four footpads. Regardless of antigen dose, adjuvant, and injection route, mice were immunized at weeks 0, 3, 6, and 9. (**b**-**c**) NAb responses induced by unmodified, Kif-treated, and Kif/endo F-treated HCV-1 sE1E2.Cut_1+2_.SPYΔN dimer vaccines (produced in HEK293F cells) against (**b**) H77 HCVpp at weeks 2, 5, 8, and 11, and (**c**) UKNP3.1.2, UKNP4.2.2, ED43, and UKNP5.2.1 HCVpps at week 11. (**d-e**) NAb responses induced by unmodified and Kif-treated HCV-1 sE1E2.Cut_1+2_.SPYΔN dimer vaccines (produced in ExpiCHO cells and tested in an independent mouse study) against (**d**) H77 HCVpp at weeks 2, 5, 8, and 11, and (**e**) UKNP3.1.2, UKNP4.2.2, ED43, and UKNP5.2.1 HCVpps at week 11. (**f**-**g**) NAb responses induced by unmodified and Kif-treated HCV-1 sE1E2.Cut_1+2_.SPYΔN-10GS-FR SApNP vaccines (produced in ExpiCHO cells and tested in an independent mouse study) against (**f**) H77 HCVpp at weeks 2, 5, 8, and 11, and (**g**) UKNP3.1.2, UKNP4.2.2, ED43, and UKNP5.2.1 HCVpps at week 11. (**h**-**i**) NAb responses induced by unmodified and Kif-treated HCV-1 sE1E2.Cut_1+2_.SPYΔN dimer vaccines measured in (**h**) female and (**i**) male mice. (**j**) NAb responses induced by unmodified and Kif-treated HCV-1 sE1E2.Cut_1+2_.SPYΔN-10GS-FR SApNP vaccines in female mice. In this study (**h**-**j**), all vaccine antigens were produced in ExpiCHO cells and adjuvanted with AH prior to administration via the intradermal route (four footpad injections) in mice. Serum from week 11 was tested against H77 HCVpp in neutralization assays. ID_50_ titers were calculated from HCVpp neutralization assays. ID_50_ values were derived by setting % neutralization to 0.0-100.0. Data points in the neutralization plots are shown as geometric mean ± geometric SD. Data were analyzed using one-way ANOVA followed by Tukey’s *post hoc* test for each time point. Two-tailed unpaired *t*-tests were used to compare geometric means of two independent groups. Statistical significance is indicated as follows: ns (not significant) and \**p* < 0.05. The mouse immunization schematic was created with BioRender.com.

We first tested the immunogenicity of HEK293F-produced HCV-1 sE1E2.Cut_1+2_.SPYΔN vaccines with and without glycan modification (**Fig. 6b** and **Fig. S6a-b**). Briefly, each injection dose contained 10 μg of HEK293F-produced antigen formulated with 100 μl of CpG/AH adjuvant. Sera were collected 2 weeks after each dose, and ID_50_ titers were plotted for comparison. None of the dimer groups showed a NAb response against H77 HCVpp at week 2, but ID_50_ titers were detected in all groups at weeks 5, 8, and 11. The glycan-trimmed dimer group elicited the lowest ID_50_ titers at weeks 8 and 11, whereas the Kif-treated dimer group showed the highest ID_50_ titers, 376 and 295, at these time points. Week-11 mouse sera were also assayed against 3.1.2, 4.2.2, ED43, and 5.2.1 HCVpps to evaluate heterologous NAb responses (**Fig. 6c** and **Fig. S6c-d**). The unmodified and Kif-treated dimer groups produced robust responses against all four HCVpps, with ID_50_ titers that were 2.5- to 16.2-fold higher than those of the glycan-trimmed group, although the differences were not statistically significant. NAb responses induced by sE1E2 dimers from both cell lines followed similar patterns, except in the case of 3.1.2 HCVpp neutralization. Overall, the HEK293F-produced dimers appeared to elicit more effective cross-NAb responses than those from ExpiCHO. While promising, this observation warrants further investigation.

Next, we performed additional immunizations as experimental repeats to further examine the effect of glycan modification on vaccine-induced NAb responses. The ExpiCHO-produced HCV-1 sE1E2.Cut_1+2_.SPYΔN antigen (10 μg), either as a dimer (unmodified vs. glycan-trimmed) or FR (unmodified vs. oligomannose-type), was mixed with CpG/AH adjuvant and administered intraperitoneally to female mice. For the dimer, measurable ID_50_ titers against H77 HCVpp were detected in both unmodified and glycan-trimmed groups at weeks 8 and 11 (**Fig. 6d** and **Fig. S6e**). The unmodified dimer elicited ID_50_ titers of 603 and 652 at weeks 8 and 11, respectively, 7.2- and 3.8-fold higher than the glycan-trimmed dimer. It also induced higher NAb titers against 4.2.2, ED43, and 5.2.1 HCVpps than the glycan-trimmed dimer, though these differences were not statistically significant (**Fig. 6e** and **Fig. S6f**). These results aligned with our earlier immunization studies (**Fig. 5d-e**). The Kif-treated FR group showed higher ID_50_ titers against H77 HCVpp than the unmodified FR group at all timepoints, and against 3.1.2, 4.2.2 (statistically significant), ED43, and 5.2.1 HCVpps at week 11 (**Fig. 6f-g** and **Fig. S6g-h**). The beneficial effect of oligomannose glycans was more evident in this analysis.

Lastly, we assessed the influence of sex on vaccine-induced NAb responses (*110, 111*). A modified regimen was used to immunize female and male mice (**Fig. 6a**). The ExpiCHO-produced HCV-1 sE1E2.Cut_1+2_.SPYΔN antigen, adjuvanted with AH, was intradermally injected into four footpads. Each injection consisted of 80 μl of antigen-adjuvant mix containing 10 μg of dimer antigen, evenly distributed across the four footpads. Sera collected at week 11, after four injections, were tested against H77 HCVpp. Within mice of the same sex, unmodified and Kif-treated dimer groups elicited comparable NAb titers, with no statistical significance (**Fig. 6h-i** and **Fig. S6i-j**). Both unmodified and Kif-treated dimer vaccines induced greater NAb responses in female mice than in males. Notably, the unmodified dimer group yielded an ID_50_ titer of 211 in female mice, which was 3.8-fold higher than in males. This result aligns with our previous study of prefusion RSV F trimer vaccines, where females developed stronger NAb responses than males following vaccination (*88*). Additionally, the unmodified and Kif-treated FR groups showed higher ID_50_ titers than their respective dimer groups at week 11 after four intradermal doses in female mice (**Fig. 6j** and **Fig. S6k**), in line with our earlier immunizations (**Fig. 5**).

Our results offer insights into factors that may influence vaccine-induced NAb responses. The ExpiCHO-produced HCV-1 sE1E2 dimer induced a more potent NAb response against H77 HCVpp (of the same genotype), while the HEK293F-produced dimer elicited higher ID_50_ titers against HCVpps from other genotypes. Female mice generated more robust NAb responses than males, as shown in an immunization study using a different adjuvant, dosage, and injection route. Two independent in vivo studies confirmed that the Kif-treated HCV-1 sE1E2.Cut_1+2_.SPYΔN-presenting SApNP is a promising vaccine candidate.

## DISCUSSION

Hepatitis C poses a significant threat to public health worldwide (*1, 112*). If acute HCV infection is not spontaneously cleared, it can progress to a persistent infection that may lead to severe liver diseases (*113*). Because early HCV infection and related liver diseases present few clinical signs or symptoms, serious liver damage often remains undiagnosed until the later stages. Injection drug use and opioid use have fueled the spread of HCV in North America and Europe, causing a “silent epidemic” with substantial economic, social, and healthcare burdens (*114*). Although DAAs are effective in treating chronic HCV infection (*4–6*), they cannot prevent reinfection or reverse liver damage (*9–11*). Therefore, to meet the WHO goal of a 90% reduction in new hepatitis infections by 2030, the development of a safe and effective HCV vaccine remains an urgent priority.

Structure-based design has become the cornerstone of modern vaccine research (*115–118*), accompanied by a growing trend toward using engineered protein NPs for more efficient delivery (*119–121*). In HCV vaccine development, immunogens based on individual epitopes, E2, and E1E2 have been designed to induce NAb responses against diverse HCV strains and quasispecies (*39, 44, 60, 61*). However, the lack of structural information on the E1E2 heterodimer has long impeded the rational design of E1E2-based HCV vaccines. Recently, cryo-EM structures have been reported for full-length E1E2 (*71*), scaffolded sE1E2 (*70*), and a dimer of E1E2 (*72*), paving the way for structure-guided vaccine design. Previously, we established a rational vaccine strategy for class I viral glycoproteins by combining antigen optimization based on metastability analysis, antigen display on multilayered SApNPs, and glycan modification (*85–89*). In this study, following a conceptually similar strategy, we designed stable sE1E2 antigens and sE1E2-presenting SApNPs for in vitro and in vivo characterization to facilitate future HCV vaccine development.

HCV E1E2 poses a unique challenge to antigen optimization (*73*). Unlike class I viral glycoproteins, which form homotrimers of heterodimers, HCV E1E2 is an asymmetric heterodimer known for its conformational disorder. In the case of HIV-1 Env, truncation and redesign of the N-terminus of heptad repeat 1 (HR1) eliminated a major source of metastability and stabilized the Env from its center (*83, 84*). However, the lack of symmetry in HCV E1E2 precludes such a design strategy. Although E2 contains a compact core formed by a disulfide-locked immunoglobulin (Ig) β-sandwich fold (*55*), E1E2 remains intrinsically flexible due to variable regions 1-3 (VR1-3) in E2 and a loosely folded E1 (*70, 71*), with nearly half (∼48%) of the protein unstructured, as indicated by DSSP analysis (*122*). Genetic diversity and a dense glycan shield further complicate E1E2 optimization (*73*). Here, we undertook a two-pronged approach to address these challenges: optimizing the sE1E2 construct for improved stability and identifying a heterodimeric scaffold best suited to accommodate sE1E2. For the former, while stem truncation was beneficial, deletion of the unstructured pFP-containing region in E1 proved most critical for preserving a native-like E1-E2 interface. For the latter, although a heterodimeric coiled coil of canonical length, GCP, outperformed other coiled coils, a covalently linked SPYΔN demonstrated a more balanced in vitro profile (e.g., expression yield, purity, stability, and antigenicity). Structural information on E1E2 (*70, 71*) was essential to our rational design, guiding the selection of optimal cutting sites on E1E2 and anchoring sites on each scaffold. The lead antigen, sE1E2.Cut_1+2_.SPYΔN, was extensively characterized using biochemical, biophysical, antigenic, and structural methods.

Findings from our sE1E2 design and assessment provide critical insights into HCV vaccine development based on these antigens. **First**, the sE1E2.Cut_1+2_ construct design is applicable across diverse HCV strains. Genotype 1a H77 and HCV-1 were more extensively evaluated than strains from genotypes 3 and 5. Among these constructs, HCV-1 sE1E2.Cut_1+2_.SPYΔN-His_6_ showed the best performance when produced in ExpiCHO cells, making it a promising vaccine antigen. The poor CHO expression of five H77 sE1E2.Cut_1+2_ scaffolds suggested a strain-rather than design-specific limitation. Thus, the sE1E2.Cut_1+2_ construct can be used to screen other strains to identify optimal vaccine backbones. **Second**, sE1E2.Cut_1+2_ scaffolds do not require AR4A to improve expression and preserve a native-like E1-E2 interface. In the first sE1E2 scaffolding study that yielded positive AR4A binding, a standard expression and purification protocol was used (*76*). In more recent studies, full-length E1E2 and another sE1E2 scaffold were coexpressed with AR4A to enhance folding and yield (*70, 71*). The intrinsic stability and high yield of sE1E2.Cut_1+2_ will facilitate the development of a robust downstream process for large-scale GMP production. **Third**, Kif treatment offers a close glycan mimic of the native E1E2 heterodimer on virions (*71*). The transmembrane domains (TMDs) of E1 and E2 contain a signal peptide that causes membrane-bound E1E2 to be statically retained in the endoplasmic reticulum (ER), resulting in stunted glycan processing and underprocessed oligomannose-type glycans. Without TMDs, sE1E2 scaffolds expressed in HEK293F and ExpiCHO cells present mostly complex-type glycans; however, Kif treatment effectively inhibits glycan processing and restores the oligomannose-type glycan profile. The beneficial effect of Kif treatment on antigenicity and immunogenicity was evident. **Lastly**, sE1E2.Cut_1+2_.SPYΔN-His_6_ provides an excellent E1E2 probe for nsEM-based epitope mapping. Although we were unable to determine high-resolution cryo-EM structures, we showed that bNAbs with known complex structures can serve as “anchors” in nsEM analysis to map the epitopes of uncharacterized antibodies onto HCV-1 sE1E2.Cut_1+2_.SPYΔN-His_6_, a native-like E1E2 “probe.” In low-resolution nsEM, the globular SPYΔN is more discernible than a coiled coil and provides a distinct structural feature to orient the sE1E2 heterodimer within the EM density.

HCV E1E2 also poses a unique challenge to particulate display. The difficulties stem from HCV E1E2 being a non-covalent heterodimer, which is incompatible with highly symmetric NPs presenting 2-fold, 3-fold, and 5-fold axes on the surface (*123*). Previously, we demonstrated that diverse class I viral glycoproteins can be displayed on 24-meric FR and 60-meric E2p and I3-01v9 by fusing the C-termini of these trimeric antigens with the N-termini of the NP subunits, which are arrayed around threefold axes on the NP surface (*82, 83, 85–87, 91*). We also redesigned the I3-01v9 SApNP (termed I3-01v9a) to optimize surface display of monomeric antigens, such as the extracellular domain of the influenza M2 protein (M2e) (*89, 90*). Recently, Sliepen et al. designed and assessed the immunogenicity of an sE2E1 2c-NP in rabbits (*79*). The vaccine potential of this immunogen design was dampened by the inability of the sE2E1-I53-50A subunit to bind AR4A and by the assumption that HCV E1E2 may form trimers of heterodimers on the virions, which is the basis for the proposed NP display (*79*). In addition, proteolytic cleavage between E2 and E1 may result in misfolded sE1 stumps mixed with sE2E1 antigens on the I53-50 2c-NP surface (*79*). In this study, we overcame these challenges by displaying sE1E2.Cut_1+2_.SPYΔN, a covalently linked heterodimer (equivalent to a monomer) with a native-like E1-E2 interface, onto FR 24-mer and I3-01v9a 60-mer, which are optimal for presenting monomers. The in vitro characterization of HCV-1 sE1E2.Cut_1+2_.SPYΔN SApNPs supported our design but also indicated that a 5GS linker is suboptimal for I3-01v9a, the 60-mer, and likely contributes to difficulties in subunit folding and NP assembly. Notwithstanding, the covalently linked sE1E2.Cut_1+2_.SPYΔN heterodimer can be readily displayed on well-established protein NP platforms for vaccine development.

In this proof-of-concept study, we immunized mice to evaluate rationally designed sE1E2 scaffolds and SApNPs with different glycan modifications. As expected, SApNPs induced a more rapid and potent NAb response than individual dimers, irrespective of the glycan type. Notably, despite its suboptimal linker length, the I3-01v9a 60-mer outperformed the FR 24-mer in inducing cross-bNAbs, underscoring the importance of NP size. Kif-treated immunogens also showed a moderate, although not statistically significant, advantage in NAb induction over unmodified versions, consistent with our previous findings for filovirus vaccines (*85*). These results inform our next steps in HCV vaccine development. First, vaccine constructs can be further refined. Variants of sE1E2.Cut_1+2_.SPYΔN can be generated by altering the spacer (sequence and length) between sE1 and SpyTag or switching SpyTag and SpyCatcherΔN in their fusions with sE1 and sE2. Screening these variants may yield sE1E2.Cut_1+2_.SPYΔN antigens with improved stability and antigenicity. The linker length in the I3-01v9a SApNP construct may be increased to improve NP assembly and NAb accessibility to key epitopes. Second, mechanistic studies are needed to understand how sE1E2 dimers and SApNPs interact with the immune system. We previously investigated vaccine trafficking, retention, presentation, and germinal center reactions in lymph nodes for four rationally designed class I viral vaccines (*82, 85, 90, 109*). Extending this strategy to HCV sE1E2 and SApNPs will offer key insights into how non-class I viral antigens engage the immune system. Lastly, optimized sE1E2 dimer and SApNP vaccines should be evaluated in more relevant animal models (*124, 125*). Previously, we identified and characterized potent NHP NAbs using V_H_1-69-like germline genes, suggesting that NHPs are a promising model for studying NAb responses to HCV E1E2 (*65, 126*). Humanized mice harboring human hepatocyte, or both human immune cells and hepatocytes, could serve as cost-effective, disease-relevant models to assess the protective efficacy of these rationally designed HCV vaccine candidates (*127, 128*).

## METHODS

### Construct design, expression, and purification of HCV sE1E2 immunogens

The E1E2 of genotype 1a H77 (GenBank Accession: AF009606.1) was used in the initial design and evaluation of sE1E2 scaffolds. Four heterodimeric coiled coils—LZ (PDB ID: 1FOS), SZ (PDB ID: 3HE5), GCP (PDB ID: 1U2U), and IAAL (PDB ID: 1U0I)—and a covalently linked SpyTag/SpyCatcher (PDB ID: 4MLI) were used as protein scaffolds to stabilize sE1E2. The E1E2s of genotype 1a HCV-1 (GenBank Accession: M62321.1), genotype 3 UKNP3.1.2 (GenBank Accession: KU285215.1), and genotype 5 UKNP5.2.1 (GenBank Accession: KU285226.1) were used to design sE1E2.Cut_1+2_.SPYΔN-His_6_ heterodimers (**Fig. S1a** and **Fig. S2a**), but only HCV-1 was used to design sE1E2.Cut_1+2_.SPYΔN-presenting FR and I3-01v9a SApNPs (**Fig. S4a**).

His_6_-tagged H77 sE1E2 scaffolds were transiently expressed in HEK293F cells. Briefly, HEK293F cells were thawed and suspended in FreeStyle^TM^ 293 Expression Medium (Life Technologies, CA) and placed in a shaker incubator at 37 °C, at 135 rpm, with 8% CO_2_. When the cells reached a density of 2.0 × 10^6^/ml, they were diluted to a concentration of 1.0 × 10^6^ ml^−1^ in 1 L of FreeStyle^TM^ 293 Expression Medium for transfection using a polyethyleneimine (PEI) (Polysciences, Inc.) transfection protocol. In brief, 800 μg of H77 sE1E2 scaffold plasmid and 300 μg of furin plasmid in 25 ml of Opti-MEM transfection medium (Life Technologies, CA) were mixed with 5 ml of PEI-MAX (1.0 mg/ml) in 25 ml of Opti-MEM and incubated for 30 min at room temperature. The HEK293F cells were then transfected with the DNA-PEI-MAX mixture and incubated in a shaker incubator at 37°C, 135 rpm, and 8% CO_2_. Five days after transfection, supernatants containing the target protein were harvested, clarified by centrifugation at 1126 × g for 22 min, and filtered through a 0.45 μm filter (Thermo Scientific). All H77 sE1E2 scaffolds were purified using Ni Sepharose excel resin (Cytiva), and the bound protein was eluted twice, each time with 15 ml of 0.5 M imidazole, and buffer-exchanged into Tris-buffered saline (TBS; pH 7.2). Nickel-purified H77 sE1E2 scaffolds were further purified by SEC using a Superdex 200 Increase 10/300 GL column (Cytiva).

His_6_-tagged H77, HCV-1, UKNP3.1.2, and UKNP5.2.1 sE1E2.Cut_1+2_.SPYΔN scaffolds, as well as all HCV-1 sE1E2.Cut_1+2_.SPYΔN-presenting SApNPs, were produced in ExpiCHO cells (Thermo Fisher). Briefly, ExpiCHO cells were thawed and incubated in ExpiCHO^TM^ Expression Medium (Thermo Fisher) in a shaker incubator at 37 °C, at 135 rpm, with 8% CO_2_. When the cells reached a density of 10 × 10^6^ ml^−1^, ExpiCHO^TM^ Expression Medium was added to adjust the cell density to 6 × 10^6^ ml^−1^ for transfection. ExpiFectamine^TM^ CHO/plasmid DNA complexes were prepared for 100-ml transfection following the manufacturer’s instructions. Kifunensine (10 mg/l, Tocris Bioscience) was added at the time of ExpiCHO transfection to inhibit α-mannosidase I to generate oligomannose-type glycans. For these constructs, 80 μg of antigen plasmid, 30 μg of furin plasmid, and 320 μl of ExpiFectamine^TM^ CHO reagent were mixed in 7.7 ml of cold OptiPRO™ medium (Thermo Fisher). After the first feed on day 1, ExpiCHO cells were cultured in a shaker incubator at 33 °C, at 115 rpm, with 8% CO_2_ according to the Max Titer protocol, with a second feed on day 5 (Thermo Fisher). Culture supernatants were harvested 13–14 days after transfection, clarified by centrifugation at 3724 × g for 25 min, and filtered using a 0.45 μm filter (Thermo Fisher). All sE1E2 scaffolds were purified using Ni Sepharose excel resin (Cytiva), and the bound protein was eluted twice, each time with 15 ml of 0.5 M imidazole, and then buffer-exchanged into Tris-buffered saline (TBS; pH 7.2). All HCV-1 sE1E2 SApNPs were purified using an AR3C antibody column, eluted three times each with 5 ml of 0.2 M Glycine (pH 2.2), neutralized by 2 M Tris-Base (pH 9.0), and buffer-exchanged into phosphate-buffered saline (PBS; pH 7.2). Nickel-purified sE1E2 scaffolds and AR3C-purified HCV-1 sE1E2 SApNPs were further polished by SEC using a Superdex 200 Increase 10/300 GL column (Cytiva) and a Superose 6 Increase 10/300 GL column (Cytiva), respectively. Selected SEC fractions were pooled, aliquoted, and frozen in liquid nitrogen or at -80 C until use.

### Expression and purification of neutralizing antibodies (NAbs)

All 11 NAbs, except for AR3C, and AR4A and CD81-Fc in the panel were provided by M. Law. AR3C and AR4A in the IgG form were transiently expressed in ExpiCHO cells (Thermo Fisher). At 12-14 days post-transfection, the cells were centrifuged at 3,724 × g for 25 min, with the supernatants filtered using a 0.45-μm filter (Millipore). IgGs were purified using protein A affinity resin (Cytiva) and eluted in 0.3 M citric acid (pH 3.0). The pH of the elution was immediately adjusted to 7.0 by adding 2 M Tris-Base (pH 9.0). The eluate was then concentrated and buffer-exchanged into phosphate buffered saline (PBS) using an Amicon 10 kDa filter (Millipore). IgG concentration was quantified by UV_280_ absorption with theoretical extinction coefficients.

### Glycan trimming by endoglycosidase F (endo F) treatment and enzyme removal

The protocol for endoglycosidase F (endo F) treatment and removal was similar to the endo H protocol used in our previous study (*82*). Endo F1, F2, and F3 were purchased from QA-Bio (catalog nos. E-EF01, E-EF02, and E-EF03, respectively). Briefly, the surface glycans of HCV-1 sE1E2.Cut_1+2_.SPYΔN-His_6_ were trimmed by mixing 1 mg of antigen with 100 μl of 5× Reaction Buffer (QA-Bio), 12.5 μl each of endo F1, F2, and F3, and H_2_O (if necessary) to make a 0.5 ml reaction. The mixture was incubated at room temperature (25 °C) for 5 h to facilitate enzymatic processing of the sE1E2 antigens. After incubation, the mixtures were passed through a Superdex 200 increase 10/300 GL to remove endo F1-3. Notably, to enable glycan trimming by endo F1-3, HCV-1 sE1E2.Cut_1+2_.SPYΔN-His_6_ was expressed in the presence of kifunensine (kif). The SEC-purified fractions were pooled aliquoted, and frozen in liquid nitrogen or at -80 C until use.

### SDS-PAGE and BN-PAGE

HCV sE1E2 scaffolds and SApNPs were analyzed by sodium dodecyl sulfate-polyacrylamide gel electrophoresis (SDS-PAGE) and blue native-polyacrylamide gel electrophoresis (BN-PAGE). The proteins were mixed with loading dye and loaded on a 10% Tris-Glycine Gel (Bio-Rad) or a 4-12% Bis-Tris NativePAGE^TM^ gel (Life Technologies). Two conditions were used in SDS-PAGE. Under non-reducing conditions, the proteins were mixed with 6× Laemmli SDS sample buffer (Thermo Scientific, catalog no. J60660) and heated to 80 °C for 5 min. Under reducing conditions, the proteins were mixed with a different Laemmli buffer (Thermo Scientific, catalog no. J61337.AC) and boiled at 100 °C for 5 min. After loading, SDS-PAGE gels were run for 25 min at 250 V using SDS running buffer (Bio-Rad). SDS-PAGE gels were stained with InstantBlue (Abcam). For BN-PAGE, the proteins were mixed with 4× native dye and loaded on a BN-PAGE gel. The gel was run for 2–2.5 h at 150 V with NativePAGE^TM^ running buffer (Life Technologies) according to the manufacturer’s instructions. BN-PAGE gels were stained with Coomassie Brilliant Blue R-250 (Bio-Rad) and destained using a solution of 6% ethanol and 3% glacial acetic acid. Gel images were acquired using a ChemiDoc^TM^ XRS+ system and processed with Image Lab version 6.1.0 build 7 Standard Edition (Bio-Rad).

### Differential scanning calorimetry (DSC)

Melting temperature (T_m_) and other thermal parameters were obtained for HCV sE1E2 scaffolds, which were expressed in either HEK293F or ExpiCHO cells and purified by a nickel column and SEC, and for HCV-1 sE1E2.Cut_1+2_.SPYΔN-10GS-FR SApNP, which was expressed in ExpiCHO cells and purified by an AR3C column and SEC, using a MicroCal PEAQ-DSC Man instrument (Malvern). Briefly, the purified protein in PBS buffer was diluted to 0.5-5 μM. Melting was probed at a scan rate of 60 °C/h from 20 °C to 100°C. Data processing, including buffer correction, normalization, and baseline subtraction, was conducted using MicroCal PEAQ-DSC software. Gaussian fitting was performed using GraphPad Prism 10.3.1 software.

### Site-specific glycan analysis

To generate site-specific glycan profiles, 60 µg aliquots of each sample were denatured for 1h in 50 mM Tris/HCl, pH 8.0 containing 6 M of urea and 5 mM dithiothreitol (DTT). Next, Env samples were reduced and alkylated by adding 20 mM iodoacetamide (IAA) and incubated for 1h in the dark, followed by a 1h incubation with 20 mM DTT to eliminate residual IAA. The alkylated Env samples were buffer exchanged into 50 mM Tris/HCl, pH 8.0 using Vivaspin columns (10 kDa) and two of the aliquots were digested separately overnight using Chymotrypsin (Mass Spectrometry Grade, Promega) or alpha-lytic protease (Sigma Aldrich) at a ratio of 1:30 (w/w). The next day, the peptides were dried and extracted using an Oasis HLB µElution Plate (Waters).

The peptides were dried again, re-suspended in 0.1% formic acid, and analyzed by nanoLC-ESI MS with a Vanquish Neo (Thermo Fisher Scientific) system coupled to an Orbitrap Eclipse Tribrid mass spectrometer (Thermo Fisher Scientific) using stepped higher energy collision-induced dissociation (HCD) fragmentation. Peptides were separated using a μPAC™ Neo HPLC Column (180 µm × 110 cm). A trapping column (PepMap 100 C18 3μM 75μM × 2cm) was used in line with the LC prior to separation with the analytical column. The LC conditions were as follows: 280-minute linear gradient consisting of 4-32% acetonitrile in 0.1% formic acid over 260 minutes followed by 20 minutes of alternating 76% acetonitrile in 0.1% formic acid and 4% ACN in 0.1% formic acid, used to ensure all the sample had eluted from the column. The flow rate was set to 300 nL/min. The spray voltage was set to 2.5 kV and the temperature of the heated capillary was set to 55 °C. The ion transfer tube temperature was set to 275 °C. The scan range was 350−2000 m/z. The stepped HCD collision energies were set to 15, 25 and 45% and the MS2 for each energy was combined. Precursor and fragment detection was performed using an Orbitrap at a resolution MS1 = 120,000, MS2 = 30,000. A standard AGC target for MS1 (4e5) and MS2 (1e4) and auto injection times (MS1 =50ms MS2 =54ms) were used.

Glycopeptide fragmentation data were extracted from the raw file using Byos (Version 5.5; Protein Metrics Inc.). The glycopeptide fragmentation data were evaluated manually for each glycopeptide; the peptide was scored as true-positive when the correct b and y fragment ions were observed along with oxonium ions corresponding to the glycan identified. The MS data was searched using the Protein Metrics 305 N-glycan library. The relative amounts of each glycan at each site as well as the unoccupied proportion were determined by comparing the extracted chromatographic areas for different glycotypes with an identical peptide sequence. All charge states for a single glycopeptide were summed. The precursor mass tolerance was set at 4 ppm and 10 ppm for fragments. A 1% false discovery rate (FDR) was applied. The relative amounts of each glycan at each site as well as the unoccupied proportion were determined by comparing the extracted ion chromatographic areas for different glycopeptides with an identical peptide sequence. Glycans were categorized according to the composition detected.

HexNAc(2)Hex(9−3) was classified as M9 to M3. Any of these structures containing a fucose were categorised as FM (fucosylated mannose). Complex-type glycans were classified according to the number of HexNAc subunits and the presence or absence of fucose. Core glycans refer to truncated structures smaller than M3. As this fragmentation method does not provide linkage information, compositional isomers are grouped.

### Bio-layer interferometry

Antigenic profiles of HCV-1 sE1E2.Cut_1+2_.SPYΔN-His_6_ heterodimer and sE1E2.Cut_1+2_.SPYΔN-10GS-FR SApNP were measured using Octet RED96 (FortéBio, Pall Life Sciences) against 10 NAbs in the human IgG form and CD81-Fc. All assays were performed with agitation set to 1000 rpm in FortéBio 1× kinetic buffer. The final volume for all solutions was 200 μl per well. Assays were performed at 30 °C in solid black 96-well plates (Geiger Bio-One). For all antigens, 5 μg/ml antibody in 1× kinetic buffer was loaded onto the surface of Anti-Human IgG Quantitation (AHQ) Biosensors for 300 s. Next, a 2-fold concentration gradient of antigen, starting at 1000 nM for HCV-1 sE1E2.Cut_1+2_.SPYΔN-His_6_ dimer and 18 nM for FR SApNP, was used in a dilution series of six. A 60-second biosensor baseline step was applied before analyzing the association of the antibody or CD81-Fc on the biosensor with the antigen in solution for 200 s. Dissociation of the interaction was followed for 300 s. The correction of baseline drift was performed by subtracting the mean shift value recorded for a sensor loaded with antibody or CD81-Fc but not incubated with antigen, and for a sensor without antibody or CD81-Fc but incubated with antigen. Octet data were processed by FortéBio’s data acquisition software v.8.1. Peak signals at the highest antigen concentration were summarized in a matrix and color-coded accordingly to allow comparisons between different constructs. Experimental data for each antigen-antibody or antigen-CD81-Fc pair were fitted using the binding equations describing a 1:1 interaction, and three datasets showing optimal fit were then grouped to determine the K_on_ and K_D_ values. The Octet binding curve was plotted using GraphPad Prism 10.3.1 software.

### Enzyme-linked immunosorbent assay (ELISA)

Costar^TM^ 96-well, high-binding, flat-bottom, half-area plates (Corning) were first coated with 50 µl of PBS containing 0.1 μg of the appropriate HCV sE1E2 dimer or SApNP antigen protein. The plates were incubated overnight at 4 °C and then washed five times with PBST wash buffer containing PBS and 0.05% (v/v) Tween 20. Each well was then blocked for 1 h at room temperature with 150 µl of blocking buffer consisting of 4% (w/v) blotting-grade blocker (Bio-Rad) in PBS. Next, plates were washed five times with PBST wash buffer. To evaluate antibody or CD81-Fc binding to the coating antigens, antibodies or CD81-Fc were diluted in blocking buffer to a maximum concentration of 10 µg/ml followed by a 10-fold dilution series. For each antibody/CD81-Fc dilution, a total volume of 50 µl was added to the appropriate wells. Plates were incubated for 1 h at room temperature and then washed five times with PBST wash buffer. For secondary antibody binding, a 1:5000 dilution of goat anti-human IgG antibody (Jackson ImmunoResearch Laboratories) was used for human and rhesus macaque NAbs and CD81-Fc, whereas a 1:3000 dilution of horseradish peroxidase (HRP)-conjugated goat anti-mouse IgG antibody (Jackson ImmunoResearch Laboratories) was used for the murine NAb A4. The secondary antibody was prepared in PBST wash buffer, and 50 µl of the diluted secondary antibody added to each well. Plates were incubated for 1 h at room temperature and then washed six times with PBST wash buffer. Lastly, the wells were developed with 50 µl of 3,3’,5,5’-tetramethylbenzidine (TMB; Life Sciences) for 3-5 min before the reaction was stopped with 50 µl of 2 N sulfuric acid. The plates were then immediately read on a BioTek Synergy plate reader at a wavelength of 450 nm. EC_50_ values were calculated from full curves using GraphPad Prism 10.3.1 software. When OD_450_ absorbance values were lower than 0.5, the EC_50_ values were set to 10 µg/ml in **Figs. 1f**, **2d**, and **4g** to facilitate EC_50_ plotting and comparison.

### Cryo-EM analysis of HCV-1 sE1E2.Cut_1+2_.SPY**Δ**N-His_6_ in complex with bNAbs

Nickel/SEC-purified HCV-1 sE1E2.Cut_1+2_.SPYΔN-His_6_ heterodimer was used to form complexes with two bNAb pairs under different conditions. AR3C and AR4A Fabs were transiently expressed in ExpiCHO cells (Thermo Fisher) and purified using G affinity resin (Cytiva), and eluted in 0.2 M glycine (pH 2.0). The pH of the elution was immediately adjusted to neutral by adding 2 M Tris-Base (pH 9.0). The purified HCV-1 sE1E2 antigen was incubated with a 1.5 × molar excess of AR3C and AR4A Fabs overnight at 4 C, followed by SEC using a Superose 6 Increase 10/300 GL column (Cytiva). HEPC74 and AR4A Fabs were transiently expressed in Expi293F GnTI-cells (Thermo Fisher) and purified by Capture Select CH1-XL Affinity Matrix (Thermo Fisher), followed by SEC with a Superdex 200 Increase 10/300 column (S200i) (Cytiva). The purified HCV-1 sE1E2 antigen was incubated with a 3× molar excess of AR4A Fab for 1 h at room temperature, followed by SEC and subsequent complexation with HEPC74 Fab.

For HCV-1 sE1E2.Cut_1+2_.SPYΔN-His_6_ bound to AR3C and AR4A, an initial sample was frozen in-house using a Vitrobot Mark IV on Quantifoil R1.2/1.3 300 mesh Cu Grids (Electron Microscopy Sciences; catalog #Q350CR1.3). Grids were screened on a Talos Arctica 200 kV Cryo-TEM equipped with a Gatan K3 camera, and an initial data set of 1,430 movies was collected. Grids from the same plunge freezing session were imaged at the National Cancer Institute’s (NCI) National Cryo-EM Facility on a Titan Krios 300 kV Cryo-TEM using a Gatan K3 camera, yielding 19,080 movies. Additional grids of the complex were also frozen using a VitroJet, aiming to produce thicker ice to improve particle orientation distribution. This yielded an additional 5,248 movies. The initial 19,080 movies were processed using cryoSPARC v4.5.3. Patch motion correction and Patch CTF were performed, after which exposures were filtered to 17,941 micrographs. An initial stack of 2000 micrographs was used to obtain particles for deep picking. Blob picking yielded a stack of 1,218,366, which was refined by 2D and 3D classification to 113,496 particles and was used to generate a model for Deep Picking. 8,699,347 particles were obtained with Deep Picker and were further refined to 85,959. To reduce anisotropy in the map, this dataset was combined with the VitroJet dataset, which included an additional 5,248 movies. Patch motion correction and Patch CTF were performed using cryoSPARC v4.5.3 (*107*). This dataset underwent blob picking, 2D classification, and heterogeneous refinement to yield 89,013 particles. After non-uniform refinement, the datasets were combined to yield the final maps using 174,972 particles. Coordinates from PDB IDs 8FSJ (*70*) and 4MWF (*55*) were fit as rigid bodies into the resulting density maps using UCSF Chimera 1.16 (*129*).

For HCV-1 sE1E2.Cut_1+2_.SPYΔN-His_6_ bound to HEPC74 and AR4A, 5,106 movies were collected on a Titan Krios 300 kV Cryo-TEM using a Gatan K3 camera (NCI National Cryo-EM Facility). Patch motion correction and Patch CTF were performed using cryoSPARC v4.5.3, after which exposures were filtered to 4,141 micrographs (*107, 130*). Blob Picking yielded a stack of 11,577,316 particles, which was filtered to 516,060 particles and subsequently used for template generation. 14,430,734 particles were picked by templates and were filtered to an initial stack of 280,335 particles. 3D variability analysis (3DVA) followed by 3D classification was performed to address moderate flexibility in the AR4A angle of approach and possible anisotropy, yielding a final stack of 36,896 particles. Coordinates of PDB ID 8FSJ (*70*) were fit as a rigid body into the final maps using UCSF Chimera 1.16 (*129*).

### Negative-stain electron microscopy (nsEM)

The nsEM analysis of HCV sE1E2 scaffolds and SApNPs was performed by the Core Microscopy Facility at The Scripps Research Institute. Briefly, samples were prepared at concentrations of 0.008 and 0.01 mg/ml, respectively. Carbon-coated copper grids (400 mesh) were glow-discharged, and 8 µl of each sample was then adsorbed for 2 min. Next, excess sample was removed, and grids were negatively stained with 2% uranyl formate for 2 min. Excess stain was wicked away, and the grids were allowed to dry. Dimer samples were analyzed at 120 kV with a Talos L120C transmission electron microscope (TEM, Thermo Fisher), and images were acquired using a CETA 16 M CMOS camera under 73,000× magnification at a resolution of 1.93 Å/pixel and defocus of 0.5-2 µm. For SApNP samples, images were collected under 52,000× magnification. The resulting pixel size at the specimen plane was 2.05 Å, and the defocus was set to −1.50 µm. Computational analysis of the images was performed using the high-performance computing core facility at The Scripps Research Institute. Briefly, the nsEM images were converted to MRC format by EMAN2 (*131*) for further processing by cryoSPARC v4.3.0 (*107*). Micrographs were contrast transfer function (CTF)-corrected by patch CTF estimation. Particles were selected using a Blob/template picker and later extracted with a box size of 140 pixel for 2D classification, with 100 and 250 Å used as the minimum and maximum particle sizes in blob picking. For all the dimer/NAb complexes, 3D models were generated by *Ab initio* reconstruction and optimized by heterogeneous and homogeneous refinement. All nsEM and fitted structure images were generated using UCSF Chimera (*129*) and Chimera X (*132*).

### Dynamic Light Scattering (DLS)

Particle size distributions of HCV-1 sE1E2.Cut_1+2_.SPYΔN-presenting FR and I3-01v9a SApNPs were obtained using a Zetasizer Ultra instrument (Malvern). Briefly, AR3C/SEC-purified SApNPs produced in ExpiCHO cells were diluted to 0.2 mg/ml in 1× PBS buffer, after which 30 μl of the prepared sample was added to a quartz batch cuvette (Malvern, catalog no. ZEN2112). Particle size was measured at 25 °C using back scattering mode. Data processing was performed on the Zetasizer using the Ultra instrument software. The resulting particle size distribution was plotted using GraphPad Prism 10.3.1 software.

### Mouse immunization and sample collection

We followed mouse immunization protocols similar to those in our previous studies (*82, 85, 88, 90*). The Association for the Assessment and Accreditation of Laboratory Animal Care (AAALAC) guidelines were followed throughout all animal experiments. The animal protocols were approved by the Institutional Animal Care and Use Committee (IACUC) of The Scripps Research Institute (TSRI). Six-to-eight-week-old female BALB/c mice were purchased from The Jackson Laboratory and housed in ventilated cages in environmentally controlled rooms at 20 °C, 50% humidity, and 12-hour light-dark cycles. In all immunization studies except the one testing the effect of sex on vaccine-induced NAb responses, our standard screening regimen was used with mice injected, via the intraperitoneal (i.p.) route, at weeks 0, 3, 6, and 9 with 3-week intervals. The injecting dose was 200 μl of antigen/adjuvant mix containing 10 μg of vaccine antigen and 100 μl of adjuvant, including 40 µg/50 µl of CpG (oligonucleotide 1826, a TLR9 agonist from InvivoGen) adsorbed onto 50 µl of AH (InvivoGen). Blood samples were collected 2 weeks after each injection through the retro-orbital sinus using heparinized capillary tubes. Samples were spun at 12000 rpm for 10 min to separate serum (top layer) and the rest of the whole blood layer. After heat inactivation at 56 °C for 30 min, serum was spun at 12000 rpm for 10 min to remove precipitates. For the last timepoint, the rest of the whole blood layer was diluted with an equal volume of PBS and then overlaid on 4.5 ml of Ficoll in a 15 ml SepMateTM tube (STEMCELL Technologies) and spun at 1200 rpm for 10 min at 20 °C to separate peripheral blood mononuclear cells (PBMCs). Cells were washed once in PBS and then resuspended in 1 ml of ACK Red Blood Cell lysis buffer (Lonza). After washing with PBS, PBMCs were resuspended in 1 ml of Freezing Media (10% DMSO/90% FCS). Two weeks after the last bleed, spleens were harvested and ground against a 70-μm cell strainer (BD Falcon) to release splenocytes into a cell suspension. Splenocytes were centrifuged, washed in PBS, treated with 5 ml of ACK lysing buffer (Lonza), and frozen with 3 ml of freezing media (10% DMSO/90% FCS). For the immunogenicity study examining sex-specific effects, a different regimen was used (*88, 90*). Six-to-eight-week-old female and male BALB/c mice were purchased from The Jackson Laboratory and housed as described above. Mice were immunized at weeks 0, 3, 6, and 9 with 80 μl of antigen/adjuvant mix containing 10 μg of vaccine antigen in 40 μl of PBS and 40 μl of AH adjuvant (InvivoGen). Intradermal (i.d.) immunization was performed via injection into four footpads, each with 20 μl of antigen/adjuvant mix, using a 29-gauge insulin needle under 3% isoflurane anesthesia with oxygen. Blood was collected from the maxillary/facial vein 2 weeks after each immunization. Serum was isolated from blood by centrifugation at 4000 rpm for 10 min. Serum was then heat-inactivated at 56 °C for 30 min, and the supernatant was collected after centrifugation at 8000 rpm for 10 min to remove debris. Heat-inactivated serum was stored at -4°C until use in HCVpp assays to determine vaccine-induce NAb responses.

### Pseudovirus production and neutralization assays

An HCV pseudoparticle (HCVpp) neutralization assay (*67*) was performed to assess the vaccine-induced NAb response in mouse sera. HCVpps were generated by co-transfection of HEK293T cells with the envelope-deficient HIV-1 pNL4-3.lucR-E-plasmid (NIH AIDS reagent program: https://www.aidsreagent.org/) and an expression plasmid encoding the E1E2 gene of genotype 1a H77 (GenBank Accession: AF009606.1), UKNP3.1.2 (GenBank Accession: KU285215.1), UKNP4.2.2 (GenBank Accession: KU285222.1), ED43 (GenBank Accession: GU814265.1), and UKNP5.2.1 (GenBank Accession: KU285226.1) at a 4:1 ratio using a Lipofectamine 3000 transfection protocol (Thermo Fisher). After 72 h, pseudoviruses were collected from the supernatant by centrifugation at 3724 × g for 10 min, aliquoted, and stored at -80 °C until use. For pseudovirus neutralization assays, mouse sera starting at a dilution of 40× were serially diluted 3-fold and incubated with pseudoviruses at 37 °C for 1 h in white solid-bottom 96-half-well plates (Corning). Next, 0.8 × 10^4^ Huh-7 cells were added to each well, and plates were incubated at 37 °C for 60 to 72 h. After incubation, the supernatant was removed, and cells were lysed. Luciferase reporter gene expression was quantified through the addition of Bright-Glo^TM^ Luciferase substrate (Promega), according to the manufacturer’s instructions. Data, expressed in relative light units (RLU), were retrieved from a BioTek microplate reader with Gen 5 software. Values from experimental wells were compared against virus-containing wells, with background luminescence from a series of uninfected wells subtracted from both. Dose-response neutralization curves were fit by nonlinear regression with constraints (0 to 100%) in GraphPad Prism 10.3.1, from which ID_50_ values were calculated and plotted for comparison.

### Statistical analysis

Data were collected from 8 mice per group in immunization studies. HCV_pp_ neutralization assays were performed in duplicate. Various HCV-1 sE1E2.Cut_1+2_.SPYΔN vaccines (e.g., sE1E2 dimer *vs.* SApNP) were compared using one-way analysis of variation (ANOVA), followed by Tukey’s multiple comparison *post hoc* test for each time point. Two-tailed unpaired *t*-tests were used when comparing two independent groups. Statistical significance is indicated in the figures as follows: ns (not significant) and \**p* < 0.05. Graphs were generated, and statistical analyses were performed using GraphPad Prism 10.3.1 software.

## ACKNOWLEDGEMENTS

The cryo-EM analysis was, in part, supported by the National Cancer Institute’s National Cryo-EM Facility at the Frederick National Laboratory for Cancer Research under contract 75N91019D00024. We acknowledge K. Vanderpool, T. Fassel, and S. Henderson of the Core Microscopy Facility at TSRI for their expert assistance with negative-stain EM analysis. **Funding:** This work was supported by NIH awards AI168251 (M.L., J.Z.) and AI168917 (M.L.), and in part by Ufovax/SFP-2018-1013 (J.Z.). **Author contributions:** Project design by L.H., and J.Z.; sE1E2 scaffold and SApNP design by L.H., and J.Z; sE1E2 scaffold and SApNP expression and purification by G.W., C.D., and L.H.; SDS and BN PAGE by G.W., C.D., Y.-Z.L., and L.H.; DSC and DSL by Y.-Z.L., and L.H.; ELISA by Y.-Z.L. and Y.-N.Z.; BLI by Y.-Z.L., and L.H.; sE1E2.LZorg expression and purification by F.G.G., L.K., E.A.T., and T.R.F.; AR3C and AR4A expression and purification by G.W., C.D., and L.H.; other NAb (and CD81-Fc) expression and purification by E.G., and M.L.; glycan analysis by M.N., J.D.A., and M.C.; cryo-EM by B.M.J. and G.O.; negative-stain EM and epitope mapping by Y.- Z.L. S.-H.H., and J.Z.; mouse immunization partly by Y.-N.Z.; HCVpp neutralization assays by G.W., C.D., Y.-N.Z., and L.H.; manuscript written by L.H., Y.-Z.L., Y.-N.Z., M.N., B.M.J., G.O., M.C., and J.Z. All authors were asked to comment on the manuscript. **Competing interests:** Dr. Jiang Zhu serves as the Co-Founder, Consultant, and Scientific Advisory Board member of Uvax Bio, LLC, and holds associated financial interests. Other authors declare that they have no competing interests. **Data and material availability:** All data needed to evaluate the conclusions of this research are available in the main text and Supplementary Information. The cryo-EM data have been deposited in the Electron Microscopy Data Bank (EMDB, https://www.ebi.ac.uk/emdb/) under accession codes EMD-70622 for the complex with AR3C and AR4A and EMD-70623 for the complex with HEPC74 and AR4A. The authors declare that the data supporting the findings of this study are available within the article and its Supplementary Information files. Source data are provided with this paper.

## Notes

### Competing Interest Statement

The authors have declared no competing interest.

