## Supplementary Information for "Native-like soluble E1E2 glycoprotein heterodimers on self-assembling protein nanoparticles for hepatitis C virus vaccine design"

### Figure S1

a

**>H77 sE1E2.LZ-His$_6$**

MGCSFSIFLLALLSCLTVPASAYQVRNSSGLYHVTNDCPNSSIVYEAADAILHTPGCVPCVREGNASRCWVAVTPTVATRDGKLPTTQLRRHIDLLVGSATLCSALYV
GDLCGSVFLVGQLFTFSPRRHWTTQDCNCSIYPGHITGHRMAWDMMMNWSPTAALVVAQLLRIPQAIMDMIAPGGRIARLEEKVKTLKAQNSELASTANMLREQVAQL
KQKVMNY**RRRRRR**ETHVTGGSAGRTTAGLVGLLTPGAKQNIQLINTNGSWHINSTALNCNESLNTGWLAGLFYQHKFNSSGCPERLASCRRLTDFAQGWGPISYANGS
GLDERPYCWHYPPRPCGIVPAKSVCGPVYCFTPSPVVVGTTDRSGAPTYSWGANDTDVFVLNNTRPPLGNWFGCTWMNSTGFTKVCGAPPCVIGGVGNNTLLCPTDCF
RKHPEATYSRCGSGPWITPRCMVDYPYRLWHYPCTINYTIFKVRMYVGGVEHRLEAACNWTRGERCDLEDRDRSELSPLLLSTTQWQVLPCSFTTLPALSTGLIHLHQ
NIVDVQYLYGVGSSIASWAIPGGLTDTLQAETDQLEDKKSALQTEIANLLKEKEKLEFILAAYGSHHHHHH

---

**>H77 sE1E2.Cut$_1$.LZ-His$_6$**

MGCSFSIFLLALLSCLTVPASAYQVRNSSGLYHVTNDCPNSSIVYEAADAILHTPGCVPCVREGNASRCWVAVTPTVATRDGKLPTTQLRRHIDLLVGSATLCSALYV
GDLCGSVFLVGQLFTFSPRRHWTTQSCNCSI**YPGH**PGGRIARLEEKVKTLKAQNSELASTANMLREQVAQLKQKVMNY**RRRRRR**ETHVTGGSAGHTTAGLVGLLTPGA
KQNIQLINTNGSWHINSTALNCNDSLTTGWLAGLFYRHKFNSSGCPERLASCRRLTDFAQGWGPISYANGSGLDERPYCWHYPPRPCGIVPAKSVCGPVYCFTPSPVV
VGTTDRSGAPTYSWGANDTDVFVLNNTRPPLGNWFGCTWMNSTGFTKVCGAPPCVIGGVGNNTLLCPTDCFRKHPEATYSRCGSGPWITPRCMVDYPYRLWHYPCTIN
YTIFKVRMYVGGVEHRLEAACNWTRGERCDLEDRDRSELSPLLLSTTQWQVLPCSFTTLPALSTGLIHLHQNIV**DVQY**PGGLTDTLQAETDQLEDKKSALQTEIANLL
KEKEKLEFILAAYGSHHHHHH

>**H77 sE1E2.Cut$_1$.SZ-His$_6$**

MGCSFSIFLLALLSCLTVPASAYQVRNSSGLYHVTNDCPNSSIVYEAADAILHTPGCVPCVREGNASRCWVAVTPTVATRDGKLPTTQLRRHIDLLVGSATLCSALYV
GDLCGSVFLVGQLFTFSPRRHWTTQSCNCSI**YPGH**PGGRNAYLRKKIARLKKDNLQLERDEQNLEKIIANLRDEIARLENEVA**RRRRRR**ETHVTGGSAGHTTAGLVGL
LTPGAKQNIQLINTNGSWHINSTALNCNDSLTTGWLAGLFYRHKFNSSGCPERLASCRRLTDFAQGWGPISYANGSGLDERPYCWHYPPRPCGIVPAKSVCGPVYCFT
PSPVVVGTTDRSGAPTYSWGANDTDVFVLNNTRPPLGNWFGCTWMNSTGFTKVCGAPPCVIGGVGNNTLLCPTDCFRKHPEATYSRCGSGPWITPRCMVDYPYRLWHY
PCTINYTIFKVRMYVGGVEHRLEAACNWTRGERCDLEDRDRSELSPLLLSTTQWQVLPCSFTTLPALSTGLIHLHQNIV**DVQY**PGGLVAQLENEVASLENENETLKKK
NLHKKDLIAYLEKEIANLRKKIEGSHHHHHH

**>H77 sE1E2.Cut$_1$.GCP-His$_6$**

MGCSFSIFLLALLSCLTVPASAYQVRNSSGLYHVTNDCPNSSIVYEAADAILHTPGCVPCVREGNASRCWVAVTPTVATRDGKLPTTQLRRHIDLLVGSATLCSALYV
GDLCGSVFLVGQLFTFSPRRHWTTQSCNCSI**YPGH**PGGEVAQLEKEVAQLEAENYQLEQEVAQLEHEG**RRRRRR**ETHVTGGSAGHTTAGLVGLLTPGAKQNIQLINTN
GSWHINSTALNCNDSLTTGWLAGLFYRHKFNSSGCPERLASCRRLTDFAQGWGPISYANGSGLDERPYCWHYPPRPCGIVPAKSVCGPVYCFTPSPVVVGTTDRSGAP
TYSWGANDTDVFVLNNTRPPLGNWFGCTWMNSTGFTKVCGAPPCVIGGVGNNTLLCPTDCFRKHPEATYSRCGSGPWITPRCMVDYPYRLWHYPCTINYTIFKVRMYV
GGVEHRLEAACNWTRGERCDLEDRDRSELSPLLLSTTQWQVLPCSFTTLPALSTGLIHLHQNIV**DVQY**PGGEVQALKKRVQALKARNYALKQKVQALRHKGGSHHHHH
H

**>H77 sE1E2.Cut$_1$.IAAL-His$_6$**

MGCSFSIFLLALLSCLTVPASAYQVRNSSGLYHVTNDCPNSSIVYEAADAILHTPGCVPCVREGNASRCWVAVTPTVATRDGKLPTTQLRRHIDLLVGSATLCSALYV
GDLCGSVFLVGQLFTFSPRRHWTTQSCNCSI**YPGH**PGGEIAALEKEIAALEKEIAALEK**RRRRRR**ETHVTGGSAGHTTAGLVGLLTPGAKQNIQLINTNGSWHINSTA
LNCNDSLTTGWLAGLFYRHKFNSSGCPERLASCRRLTDFAQGWGPISYANGSGLDERPYCWHYPPRPCGIVPAKSVCGPVYCFTPSPVVVGTTDRSGAPTYSWGANDT
DVFVLNNTRPPLGNWFGCTWMNSTGFTKVCGAPPCVIGGVGNNTLLCPTDCFRKHPEATYSRCGSGPWITPRCMVDYPYRLWHYPCTINYTIFKVRMYVGGVEHRLEA
ACNWTRGERCDLEDRDRSELSPLLLSTTQWQVLPCSFTTLPALSTGLIHLHQNIV**DVQY**PGGKIAALKEKIAALKEKIAALKEGSHHHHHH

>**H77 sE1E2.Cut$_1$.SPY$\Delta$N-His$_6$**

MGCSFSIFLLALLSCLTVPASAYQVRNSSGLYHVTNDCPNSSIVYEAADAILHTPGCVPCVREGNASRCWVAVTPTVATRDGKLPTTQLRRHIDLLVGSATLCSALYV
GDLCGSVFLVGQLFTFSPRRHWTTQSCNCSI**YPGH**GSGSVPTIVMVDAYKRYK**RRRRRR**ETHVTGGSAGHTTAGLVGLLTPGAKQNIQLINTNGSWHINSTALNCNDS
LTTGWLAGLFYRHKFNSSGCPERLASCRRLTDFAQGWGPISYANGSGLDERPYCWHYPPRPCGIVPAKSVCGPVYCFTPSPVVVGTTDRSGAPTYSWGANDTDVFVLN
NTRPPLGNWFGCTWMNSTGFTKVCGAPPCVIGGVGNNTLLCPTDCFRKHPEATYSRCGSGPWITPRCMVDYPYRLWHYPCTINYTIFKVRMYVGGVEHRLEAACNWTR
GERCDLEDRDRSELSPLLLSTTQWQVLPCSFTTLPALSTGLIHLHQNIV**DVQYAS**DSATHIKFSKRDEDGRELAGATMELRDSSGKTISTWISDGHVKDFYLYPGKYT
FVETAAPDGYEVATAITFTVNEQGQVTVNGEATKGDAHTGSHHHHHH

---

**>H77 sE1E2.Cut$_{1+2}$.LZ-His$_6$**

MGCSFSIFLLALLSCLTVPASAYQVRNSSGLYHVTNDCPNSSIVYEAADAILHTPGCVPCVREGNASRCWVAVTPTVATRDGKLPTTQLRRHIDGSPRRHWTTQSCNC
SI**YPGH**PGGRIARLEEKVKTLKAQNSELASTANMLREQVAQLKQKVMNY**RRRRRR**ETHVTGGSAGHTTAGLVGLLTPGAKQNIQLINTNGSWHINSTALNCNDSLTTG
WLAGLFYRHKFNSSGCPERLASCRRLTDFAQGWGPISYANGSGLDERPYCWHYPPRPCGIVPAKSVCGPVYCFTPSPVVVGTTDRSGAPTYSWGANDTDVFVLNNTRP
PLGNWFGCTWMNSTGFTKVCGAPPCVIGGVGNNTLLCPTDCFRKHPEATYSRCGSGPWITPRCMVDYPYRLWHYPCTINYTIFKVRMYVGGVEHRLEAACNWTRGERC
DLEDRDRSELSPLLLSTTQWQVLPCSFTTLPALSTGLIHLHQNIV**DVQY**PGGLTDTLQAETDQLEDKKSALQTEIANLLKEKEKLEFILAAYGSHHHHHH

**>H77 sE1E2.Cut$_{1+2}$.SZ-His$_6$**

MGCSFSIFLLALLSCLTVPASAYQVRNSSGLYHVTNDCPNSSIVYEAADAILHTPGCVPCVREGNASRCWVAVTPTVATRDGKLPTTQLRRHIDGSPRRHWTTQSCNC
SI**YPGH**PGGRNAYLRKKIARLKKDNLQLERDEQNLEKIIANLRDEIARLENEVA**RRRRRR**ETHVTGGSAGHTTAGLVGLLTPGAKQNIQLINTNGSWHINSTALNCND
SLTTGWLAGLFYRHKFNSSGCPERLASCRRLTDFAQGWGPISYANGSGLDERPYCWHYPPRPCGIVPAKSVCGPVYCFTPSPVVVGTTDRSGAPTYSWGANDTDVFVL
NNTRPPLGNWFGCTWMNSTGFTKVCGAPPCVIGGVGNNTLLCPTDCFRKHPEATYSRCGSGPWITPRCMVDYPYRLWHYPCTINYTIFKVRMYVGGVEHRLEAACNWT
RGERCDLEDRDRSELSPLLLSTTQWQVLPCSFTTLPALSTGLIHLHQNIV**DVQY**PGGLVAQLENEVASLENENETLKKKNLHKKDLIAYLEKEIANLRKKIEGSHHHH
HH

**>H77 sE1E2.Cut$_{1+2}$.GCP-His$_6$**

MGCSFSIFLLALLSCLTVPASAYQVRNSSGLYHVTNDCPNSSIVYEAADAILHTPGCVPCVREGNASRCWVAVTPTVATRDGKLPTTQLRRHIDGSPRRHWTTQSCNC
SI**YPGH**PGGEVAQLEKEVAQLEAENYQLEQEVAQLEHEG**RRRRRR**ETHVTGGSAGHTTAGLVGLLTPGAKQNIQLINTNGSWHINSTALNCNDSLTTGWLAGLFYRHK
FNSSGCPERLASCRRLTDFAQGWGPISYANGSGLDERPYCWHYPPRPCGIVPAKSVCGPVYCFTPSPVVVGTTDRSGAPTYSWGANDTDVFVLNNTRPPLGNWFGCTW
MNSTGFTKVCGAPPCVIGGVGNNTLLCPTDCFRKHPEATYSRCGSGPWITPRCMVDYPYRLWHYPCTINYTIFKVRMYVGGVEHRLEAACNWTRGERCDLEDRDRSEL
SPLLLSTTQWQVLPCSFTTLPALSTGLIHLHQNIV**DVQY**PGGEVQALKKRVQALKARNYALKQKVQALRHKGGSHHHHHH

### Figure S1

#### a (continued)

**>H77 sE1E2.Cut$_{1+2}$.IAAL-His$_6$**

MGCSFSIFLLALLSCLTVPASAYQVRNSSGLYHVTNDCPNSSIVYEAADAILHTPGCVPCVREGNASRCWVAVTPTVATRDGKLPTTQLRRHIDGSPRRHWTTQSCNC
SI**YPGH**PGGEIAALEKEIAALEKEIAALEK**RRRRRR**ETHVTGGSAGHTTAGLVGLLTPGAKQNIQLINTNGSWHINSTALNCNDSLTTGWLAGLFYRHKFNSSGCPER
LASCRRLTDFAQGWGPISYANGSGLDERPYCWHYPPRPCGIVPAKSVCGPVYCFTPSPVVVGTTDRSGAPTYSWGANDTDVFVLNNTRPPLGNWFGCTWMNSTGFTKV
CGAPPCVIGGVGNNTLLCPTDCFRKHPEATYSRCGSGPWITPRCMVDYPYRLWHYPCTINYTIFKVRMYVGGVEHRLEAACNWTRGERCDLEDRDRSELSPLLLSTTQ
WQVLPCSFTTLPALSTGLIHLHQNIV**DVQY**PGGKIAALKEKIAALKEKIAALKEGSHHHHHH

**>H77 sE1E2.Cut$_{1+2}$.SPYΔN-His$_6$**

MGCSFSIFLLALLSCLTVPASAYQVRNSSGLYHVTNDCPNSSIVYEAADAILHTPGCVPCVREGNASRCWVAVTPTVATRDGKLPTTQLRRHIDGSPRRHWTTQSCNC
SI**YPGH**GSGSVPTIVMVDAYKRYK**RRRRRR**ETHVTGGSAGHTTAGLVGLLTPGAKQNIQLINTNGSWHINSTALNCNDSLTTGWLAGLFYRHKFNSSGCPERLASCRR
LTDFAQGWGPISYANGSGLDERPYCWHYPPRPCGIVPAKSVCGPVYCFTPSPVVVGTTDRSGAPTYSWGANDTDVFVLNNTRPPLGNWFGCTWMNSTGFTKVCGAPPC
VIGGVGNNTLLCPTDCFRKHPEATYSRCGSGPWITPRCMVDYPYRLWHYPCTINYTIFKVRMYVGGVEHRLEAACNWTRGERCDLEDRDRSELSPLLLSTTQWQVLPC
SFTTLPALSTGLIHLHQNIV**DVQYAS**DSATHIKFSKRDEDGRELAGATMELRDSSGKTISTWISDGHVKDFYLYPGKYTFVETAAPDGYEVATAITFTVNEQGQVTVN
GEATKGDAHTGSHHHHHH

MGCSFSIFLLALLSCLTVPASA: Leader sequence
[257]QLRRHIDLLVGSATL[272]CSALYVGDLCGSVF[285]LVGQLFTF[293]: Unstructured, putative fusion peptide (pFP, 272-285)-containing region
: Heterodimeric scaffold domain-1
RRRRRR: Furin cleavage site motif
: Heterodimeric scaffold domain-2
PGG: Coiled-coil linker; GS: Linker; G: Linker; GSGS: Linker
AS: Enzymatic site
HHHHHH: His$_6$-tag

#### b ELISA curves of HEK293F-expressed H77 sE1E2 scaffolds binding to bNAbs AR3C and AR4A

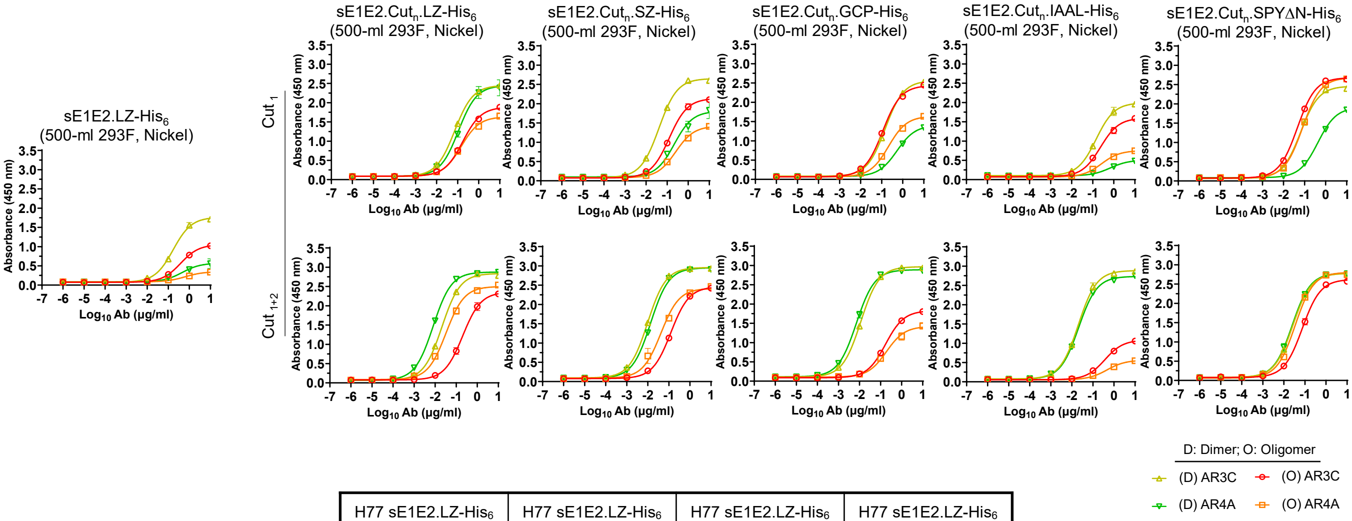


| H77 sE1E2.LZ-His$_6$ (O) AR3C | | H77 sE1E2.LZ-His$_6$ (O) AR4A | | H77 sE1E2.LZ-His$_6$ (D) AR3C | | H77 sE1E2.LZ-His$_6$ (D) AR4A | |
|---|---|---|---|---|---|---|---|
| $EC_{50}$ | STD | $EC_{50}$ | STD | $EC_{50}$ | STD | $EC_{50}$ | STD |
| 0.383 | 0.0335 | 10.0 | — | 0.172 | 0.00870 | 10.0 | — |

| H77 sE1E2.Cut$_1$.LZ-His$_6$ (O) AR3C | | H77 sE1E2.Cut$_1$.LZ-His$_6$ (O) AR4A | | H77 sE1E2.Cut$_1$.LZ-His$_6$ (D) AR3C | | H77 sE1E2.Cut$_1$.LZ-His$_6$ (D) AR4A | |
|---|---|---|---|---|---|---|---|
| $EC_{50}$ | STD | $EC_{50}$ | STD | $EC_{50}$ | STD | $EC_{50}$ | STD |
| 0.175 | 0.00156 | 0.147 | 0.0135 | 0.0722 | 0.00556 | 0.109 | 0.000636 |

| H77 sE1E2.Cut$_{1+2}$.LZ-His$_6$ (O) AR3C | | H77 sE1E2.Cut$_{1+2}$.LZ-His$_6$ (O) AR4A | | H77 sE1E2.Cut$_{1+2}$.LZ-His$_6$ (D) AR3C | | H77 sE1E2.Cut$_{1+2}$.LZ-His$_6$ (D) AR4A | |
|---|---|---|---|---|---|---|---|
| $EC_{50}$ | STD | $EC_{50}$ | STD | $EC_{50}$ | STD | $EC_{50}$ | STD |
| 0.204 | 0.0419 | 0.0329 | 0.00216 | 0.0210 | 0.000467 | 0.00805 | 0.000363 |

| H77 sE1E2.Cut$_1$.SZ-His$_6$ (O) AR3C | | H77 sE1E2.Cut$_1$.SZ-His$_6$ (O) AR4A | | H77 sE1E2.Cut$_1$.SZ-His$_6$ (D) AR3C | | H77 sE1E2.Cut$_1$.SZ-His$_6$ (D) AR4A | |
|---|---|---|---|---|---|---|---|
| $EC_{50}$ | STD | $EC_{50}$ | STD | $EC_{50}$ | STD | $EC_{50}$ | STD |
| 0.126 | 0.0102 | 0.254 | 0.00240 | 0.0410 | 0.000247 | 0.219 | 0.0215 |

| H77 sE1E2.Cut$_{1+2}$.SZ-His$_6$ (O) AR3C | | H77 sE1E2.Cut$_{1+2}$.SZ-His$_6$ (O) AR4A | | H77 sE1E2.Cut$_{1+2}$.SZ-His$_6$ (D) AR3C | | H77 sE1E2.Cut$_{1+2}$.SZ-His$_6$ (D) AR4A | |
|---|---|---|---|---|---|---|---|
| $EC_{50}$ | STD | $EC_{50}$ | STD | $EC_{50}$ | STD | $EC_{50}$ | STD |
| 0.1238 | 0.00 | 0.0406 | 0.00851 | 0.00980 | 0.000185 | 0.0136 | 0.000219 |

| H77 sE1E2.Cut$_1$.GCP-His$_6$ (O) AR3C | | H77 sE1E2.Cut$_1$.GCP-His$_6$ (O) AR4A | | H77 sE1E2.Cut$_1$.GCP-His$_6$ (D) AR3C | | H77 sE1E2.Cut$_1$.GCP-His$_6$ (D) AR4A | |
|---|---|---|---|---|---|---|---|
| $EC_{50}$ | STD | $EC_{50}$ | STD | $EC_{50}$ | STD | $EC_{50}$ | STD |
| 0.112 | 0.00 | 0.210 | 0.0159 | 0.148 | 0.00686 | 0.529 | 0.0153 |

| H77 sE1E2.Cut$_{1+2}$.GCP-His$_6$ (O) AR3C | | H77 sE1E2.Cut$_{1+2}$.GCP-His$_6$ (O) AR4A | | H77 sE1E2.Cut$_{1+2}$.GCP-His$_6$ (D) AR3C | | H77 sE1E2.Cut$_{1+2}$.GCP-His$_6$ (D) AR4A | |
|---|---|---|---|---|---|---|---|
| $EC_{50}$ | STD | $EC_{50}$ | STD | $EC_{50}$ | STD | $EC_{50}$ | STD |
| 0.172 | 0.00163 | 0.188 | 0.00438 | 0.0114 | 0.000474 | 0.00708 | 0.000986 |

| H77 sE1E2.Cut$_1$.IAAL-His$_6$ (O) AR3C | | H77 sE1E2.Cut$_1$.IAAL-His$_6$ (O) AR4A | | H77 sE1E2.Cut$_1$.IAAL-His$_6$ (D) AR3C | | H77 sE1E2.Cut$_1$.IAAL-His$_6$ (D) AR4A | |
|---|---|---|---|---|---|---|---|
| $EC_{50}$ | STD | $EC_{50}$ | STD | $EC_{50}$ | STD | $EC_{50}$ | STD |
| 0.231 | 0.00417 | 0.261 | 0.0155 | 0.145 | 0.00318 | 10.0 | — |

| H77 sE1E2.Cut$_{1+2}$.IAAL-His$_6$ (O) AR3C | | H77 sE1E2.Cut$_{1+2}$.IAAL-His$_6$ (O) AR4A | | H77 sE1E2.Cut$_{1+2}$.IAAL-His$_6$ (D) AR3C | | H77 sE1E2.Cut$_{1+2}$.IAAL-His$_6$ (D) AR4A | |
|---|---|---|---|---|---|---|---|
| $EC_{50}$ | STD | $EC_{50}$ | STD | $EC_{50}$ | STD | $EC_{50}$ | STD |
| 0.365 | 0.00707 | 0.473 | 0.183 | 0.0204 | 0.000643 | 0.0209 | 0.000445 |

| H77 sE1E2.Cut$_1$.SPYΔN-His$_6$ (O) AR3C | | H77 sE1E2.Cut$_1$.SPYΔN-His$_6$ (O) AR4A | | H77 sE1E2.Cut$_1$.SPYΔN-His$_6$ (D) AR3C | | H77 sE1E2.Cut$_1$.SPYΔN-His$_6$ (D) AR4A | |
|---|---|---|---|---|---|---|---|
| $EC_{50}$ | STD | $EC_{50}$ | STD | $EC_{50}$ | STD | $EC_{50}$ | STD |
| 0.0453 | 0.000509 | 0.0800 | 0.00532 | 0.0649 | 0.00216 | 0.431 | 0.0163 |

| H77 sE1E2.Cut$_{1+2}$.SPYΔN-His$_6$ (O) AR3C | | H77 sE1E2.Cut$_{1+2}$.SPYΔN-His$_6$ (O) AR4A | | H77 sE1E2.Cut$_{1+2}$.SPYΔN-His$_6$ (D) AR3C | | H77 sE1E2.Cut$_{1+2}$.SPYΔN-His$_6$ (D) AR4A | |
|---|---|---|---|---|---|---|---|
| $EC_{50}$ | STD | $EC_{50}$ | STD | $EC_{50}$ | STD | $EC_{50}$ | STD |
| 0.0751 | 0.00629 | 0.0346 | 0.000438 | 0.0266 | 0.00252 | 0.0247 | 0.00303 |

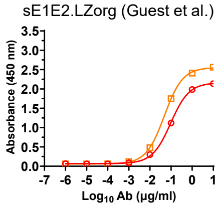


| H77 sE1E2.LZorg (Guest et al.) AR3C | | H77 sE1E2.LZorg (Guest et al.) AR4A | |
|---|---|---|---|
| $EC_{50}$ | STD | $EC_{50}$ | STD |
| 0.0962 | 0.00419 | 0.0484 | 0.00368 |

#### C SEC profile of His$_6$-tagged H77 sE1E2.Cut$_{1+2}$ scaffolds expressed in ExpiCHO cells

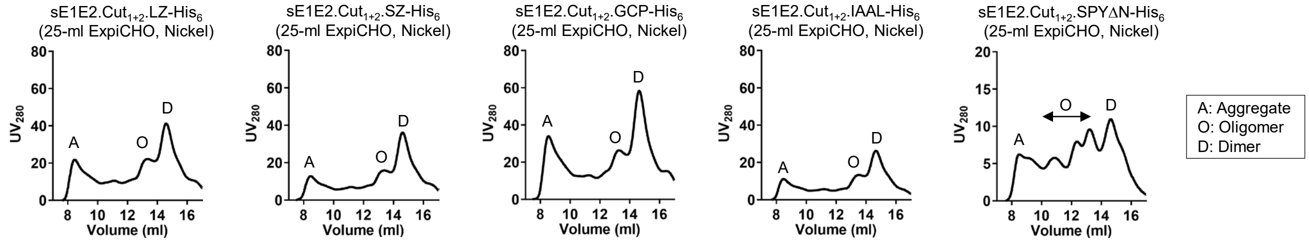


**Fig. S1. Rational design and in vitro characterization of genotype 1a H77 sE1E2 scaffolds.** (**a**) Amino acid sequences of 11 genotype 1a H77 sE1E2 scaffolds, including sE1E2.LZ, sE1E2.Cut$_1$ fused to the LZ, SZ, GCP, IAAL, and SPYΔN scaffolds, and sE1E2.Cut$_{1+2}$ fused to the LZ, SZ, GCP, IAAL, and SPYΔN scaffolds. Of these scaffolds, LZ, SZ, GCP, and IAAL are heterodimeric coiled coils of different sizes, whereas SPYΔN is a modified SpyTag/SpyCatcher with an N-terminal truncation. Two modifications are incorporated into H77 sE1E2: Cut$_1$ indicates the stem truncation of E1 at H312 and E2 at Y703, and Cut$_2$ indicates the deletion of an E1 region (L264-F293) that encompasses the putative fusion peptide (pFP). Signal peptide, E1 region L264-F293 (and pFP), heterodimeric scaffold domain-1, furin cleavage motif, heterodimeric scaffold domain-2, short linkers, enzymatic site, and His$_6$ tag are highlighted in yellow, gray (dark gray), green, red, orange, pink, blue, and cyan, respectively. (**b**) ELISA curves of H77 sE1E2.LZ, five H77 sE1E2.Cut$_1$ scaffolds, five H77 sE1E2.Cut$_{1+2}$ scaffolds, and an sE1E2.LZorg antigen provided by Fuerst and colleagues that was produced using the reported AR4A-coexpression protocol (Guest et al., *PNAS,* 2021), binding to bNAbs AR3C and AR4A. Briefly, each well was coated with 0.1 μg of the appropriate antigen, and antibodies were diluted in a 10-fold dilution series from a starting concentration of 10 μg/ml for the two bNAbs. ELISA-derived half-maximal effective concentration ($EC_{50}$) and standard deviation (SD) values are summarized in a table. (**c**) SEC profiles of five H77 sE1E2.Cut$_{1+2}$ scaffolds expressed in 25-ml ExpiCHO cultures and purified using a nickel column followed by a Superdex 200 Increase 10/300 column.

#### Figure S2

a

**>HCV-1 sE1E2.Cut$_{1+2}$.SPYΔN-His$_6$**
MGCSFSIFLLALLSCLTVPASAYQVRNSTGLYHVTNDCPNSSIVYEAADAILHTPGCVPCVREGNASRCWVAMTPTVATRDGKLPATQLRRHIDGSPRRHWTTQGCNC
SI**YPGH**GSGSVPTIVMVDAYKRYKRRRRRRETHVTGGSAGHTVSGFVSLLAPGAKQNVQLINTNGSWHLNSTALNCNDSLNTGWLAGLFYHHKFNSSGCPERLASCRP
LTDFDQGWGPISYANGSGPDQRPYCWHYPPKPCGIVPAKSVCGPVYCFTPSPVVVGTTDRSGAPTYSWGENDTDVFVLNNTRPPLGNWFGCTWMNSTGFTKVCGAPPC
VIGGAGNNTLHCPTDCFRKHPDATYSRCGSGPWITPRCLVDYPYRLWHYPCTINYTIFKIRMYVGGVEHRLEAACNWTRGERCDLEDRDRSELSPLLLTTTQWQVLPC
SFTTLPALSTGLIHLHQNIV**DVQY**ASDSATHIKFSKRDEDGRELAGATMELRDSSGKTISTWISDGHVKDFYLYPGKYTFVETAAPDGYEVATAITFTVNEQGQVTVN
GEATKGDAHTGSHHHHHH

**>UKNP3.1.2 sE1E2.Cut$_{1+2}$.SPYΔN-His$_6$**
MGCSFSIFLLALLSCLTVPASALEWRNTSGLYILTNDCPNSSIVYEADDVILHTPGCIPCVQDGNTSTCWTSVTPTVAVRYVGATTASIRSHVDGRPRRHQTVQTCNC
SL**YPGH**GSGSVPTIVMVDAYKRYKRRRRRRSTYTTGGQAARGARGIAGLFDLGARQNLQLVNTNGSWHINRTALNCNESINTGFIAGLFYYHKFNSTGCPQRLSSCKP
ITSFRQGWGPLTDANITGSSDDKPYCWHYAPRPCESVPASKVCGPVYCFTPSPVVVGTTDAKGVPTYTWGANETDVFLLNSLRPPKGRWFGCTWMNSTGFTKTCGAPP
CNIYGGGGNSKNESDLFCPTDCFRKHPEATYSRCGAGPWLTPRCMVDYPYRLWHYPCTVNFTLFQVRMFVGGFEHRFTAACNWTRGERCDIEDRDRSEQHPLLHSTTE
LAILPCSFTPMPAWSTGLIHLHQNIV**DVQY**ASDSATHIKFSKRDEDGRELAGATMELRDSSGKTISTWISDGHVKDFYLYPGKYTFVETAAPDGYEVATAITFTVNEQ
GQVTVNGEATKGDAHTGSHHHHHH

**>UKNP5.2.1 sE1E2.Cut$_{1+2}$.SPYΔN-His$_6$**
MGCSFSIFLLALLSCLTVPASAVPYRNASGVYHVTNDCPNSSIVYEANDLILHAPGCVPCVRTGNVSRCWVQITPTLSAPSLGAITAPLRRAVDGRPRQHATVQDCNC
SI**YSGH**GSGSVPTIVMVDAYKRYKRRRRRRETHSVGGVVARDLRSLTSIFTPGPRQNLQLVNTNGSWHINRTALNCQDSLQTGFIAGLLYFNKFNSSGCPERMASCRP
LTAFDQGWGTISYADMSGPSDDKPYCWHYPPRPCGVVPAQSVCGPVYCFTPSPVVVGTTDRRGYPTYNWGSNETDVILLNSTRPPAGSWFGCTWMNATGFVKTCGAPP
CNLGPTGNTSLKCPTDCFRKHPEATYTRCGSGPWLTPRCLVDYPYRLWHYPCTVNYTIFKVRMYIGGLEHRLDAACNWTRGERCDLEDRDRAELSPLLHTTTQWAILP
CSFTPTPALSTGLIHLHQNIV**DTQY**ASDSATHIKFSKRDEDGRELAGATMELRDSSGKTISTWISDGHVKDFYLYPGKYTFVETAAPDGYEVATAITFTVNEQGQVTV
NGEATKGDAHTGSHHHHHH

MGCSFSIFLLALLSCLTVPASA: Leader sequence
$^{257}$QLRRHID$^{263}$: Truncated putative fusion peptide (pFP, 272-285)-containing region
: SpyTag002
RRRRRR: Furin cleavage site motif
: SpyCatcher002ΔN
G: Linker; GSGS: Linker
AS: Enzymatic site
HHHHHH: His$_6$-tag

b **Reducing SDS-PAGE of ExpiCHO-expressed sE1E2.Cut$_{1+2}$.SPYΔN-His$_6$**

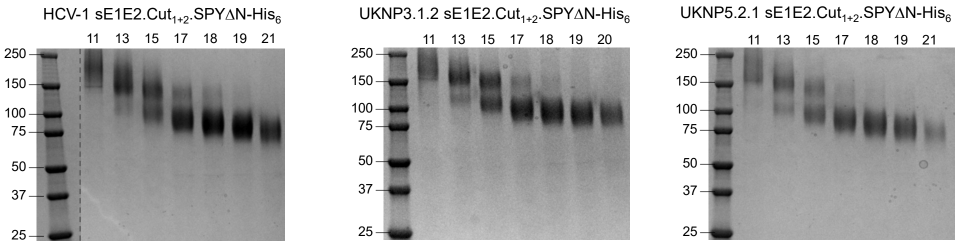


c **SEC profiles of Kif- and Kif/endo F-treated HCV-1 sE1E2.Cut$_{1+2}$.SPYΔN-His$_6$**

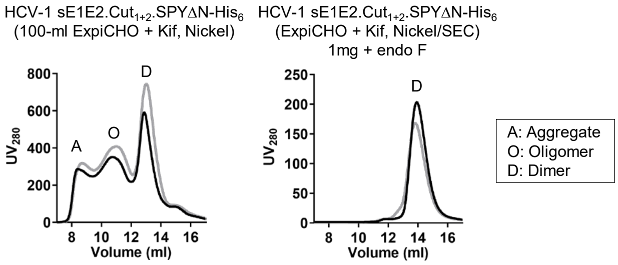


d **DSC profiles of Kif- and Kif/endo F-treated HCV-1 sE1E2.Cut$_{1+2}$.SPYΔN-His$_6$**

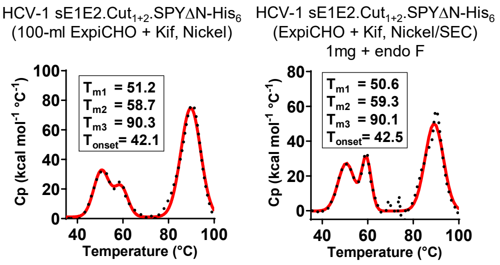

### Figure S2

e

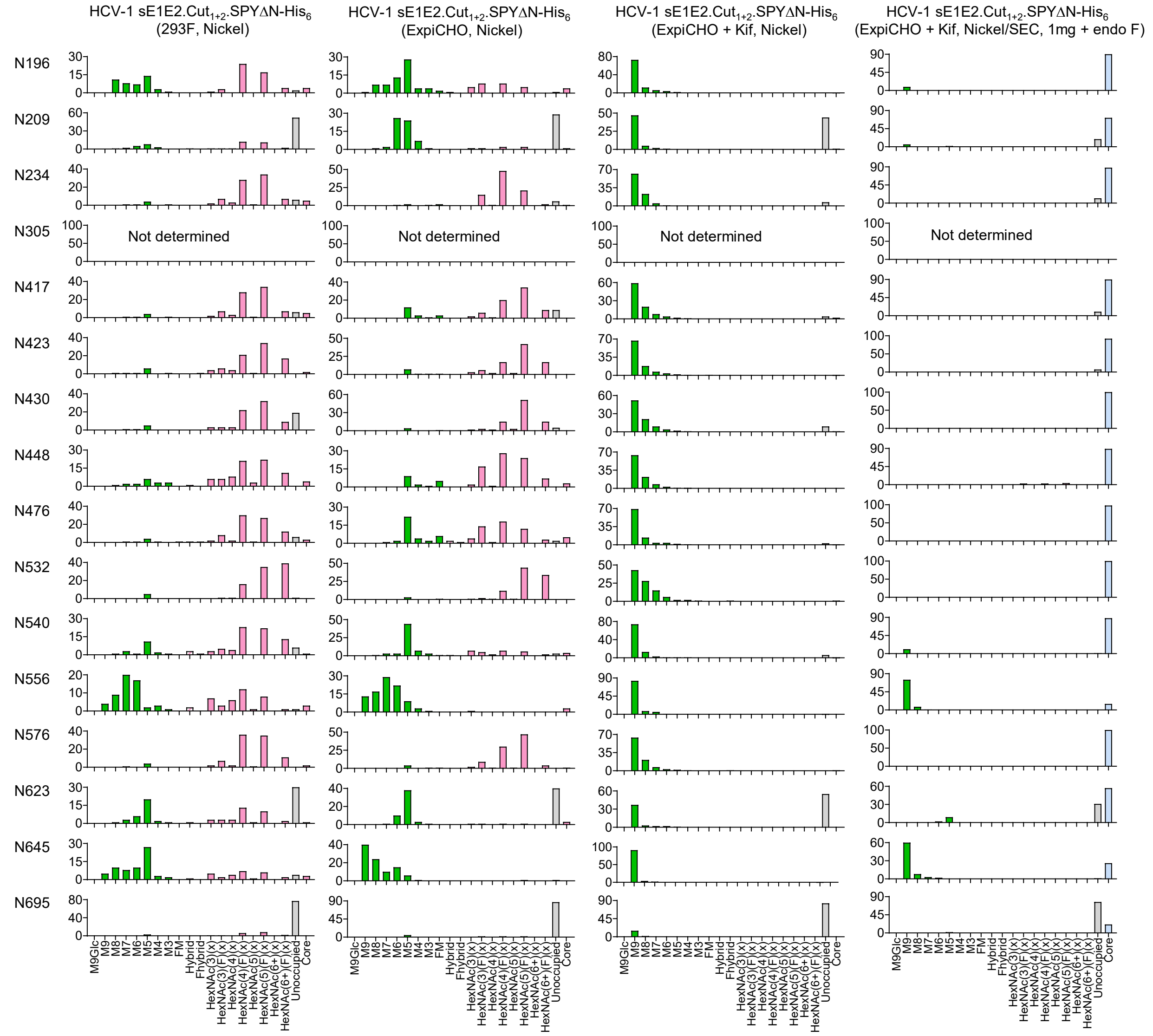


f

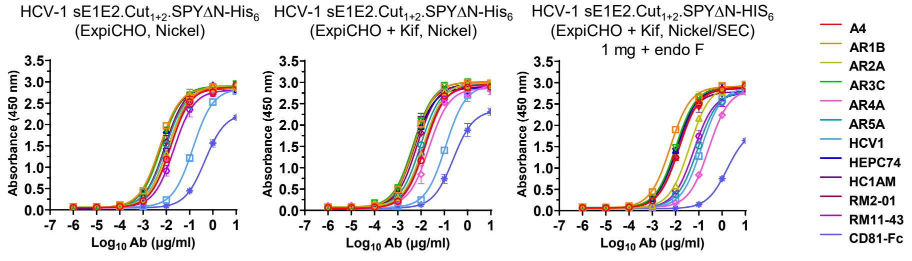


**HCV1 sE1E2.Cut$_{1+2}$.SPYΔN-His$_6$ (ExpiCHO, Nickel)**

| A4 | | AR1B | | AR2A | | AR3C | | AR4A | | AR5A | |
|---|---|---|---|---|---|---|---|---|---|---|---|
| $EC_{50}$ | STD | $EC_{50}$ | STD | $EC_{50}$ | STD | $EC_{50}$ | STD | $EC_{50}$ | STD | $EC_{50}$ | STD |
| 0.0149 | 0.00104 | 0.00512 | 2.55E-05 | 0.0124 | 0.000615 | 0.00607 | 0.000216 | 0.0150 | 0.000255 | 0.00923 | 0.00102 |

| HCV1 | | HEPC74 | | HC1AM | | RM2-01 | | RM11-43 | | CD81-Fc | |
|---|---|---|---|---|---|---|---|---|---|---|---|
| $EC_{50}$ | STD | $EC_{50}$ | STD | $EC_{50}$ | STD | $EC_{50}$ | STD | $EC_{50}$ | STD | $EC_{50}$ | STD |
| 0.140 | 0.00552 | 0.00574 | 0.000144 | 0.0116 | 6.36E-05 | 0.00873 | 0.00214 | 0.0215 | 0.00504 | 0.461 | 0.0463 |

### Figure S2

**HCV-1 sE1E2.Cut$_{1+2}$.SPYΔN-His$_6$ (ExpiCHO + Kif, Nickel)**

| A4 | | AR1B | | AR2A | | AR3C | | AR4A | | AR5A | |
|---|---|---|---|---|---|---|---|---|---|---|---|
| EC$_{50}$ | STD | EC$_{50}$ | STD | EC$_{50}$ | STD | EC$_{50}$ | STD | EC$_{50}$ | STD | EC$_{50}$ | STD |
| 0.0149 | 0.000728 | 0.00520 | 0.000313 | 0.0128 | 0.000629 | 0.00473 | 0.000809 | 0.0218 | 0.00369 | 0.00945 | 0.000252 |

| HCV1 | | HEPC74 | | HCIAM | | RM2-01 | | RM11-43 | | CD81-FC | |
|---|---|---|---|---|---|---|---|---|---|---|---|
| EC$_{50}$ | STD | EC$_{50}$ | STD | EC$_{50}$ | STD | EC$_{50}$ | STD | EC$_{50}$ | STD | EC$_{50}$ | STD |
| 0.114 | 0.00219 | 0.00494 | 5.30E-05 | 0.00434 | 0.000762 | 0.00660 | 0.000438 | 0.0126 | 0.00178 | 0.249 | 0.0953 |

**HCV-1 sE1E2.Cut$_{1+2}$.SPYΔN-His$_6$ (ExpiCHO + Kif, Nickel/SEC, 1 mg + endo F)**

| A4 | | AR1B | | AR2A | | AR3C | | AR4A | | AR5A | |
|---|---|---|---|---|---|---|---|---|---|---|---|
| EC$_{50}$ | STD | EC$_{50}$ | STD | EC$_{50}$ | STD | EC$_{50}$ | STD | EC$_{50}$ | STD | EC$_{50}$ | STD |
| 0.0144 | 0.00173 | 0.00590 | 9.12E-05 | 0.0401 | 0.00207 | 0.0101 | 0.000175 | 0.269 | 0.0159 | 0.0883 | 0.00046 |

| HCV1 | | HEPC74 | | HCIAM | | RM2-01 | | RM11-43 | | CD81-FC | |
|---|---|---|---|---|---|---|---|---|---|---|---|
| EC$_{50}$ | STD | EC$_{50}$ | STD | EC$_{50}$ | STD | EC$_{50}$ | STD | EC$_{50}$ | STD | EC$_{50}$ | STD |
| 0.130 | 0.0131 | 0.0110 | 0.000467 | 0.0110 | 0.000325 | 0.0135 | 0.00146 | 0.061 | 0.0136 | 1.65 | 0.0127 |

g

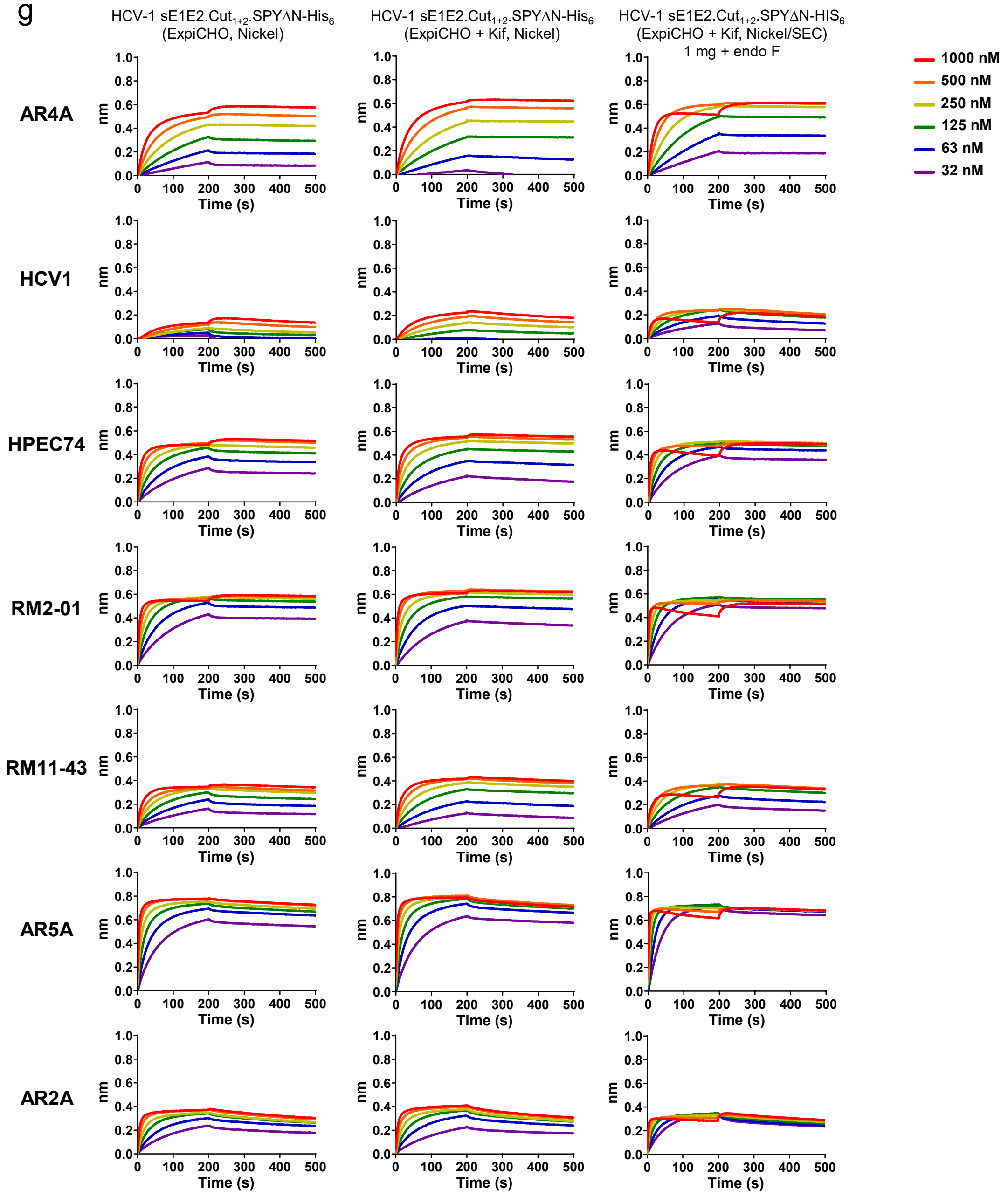

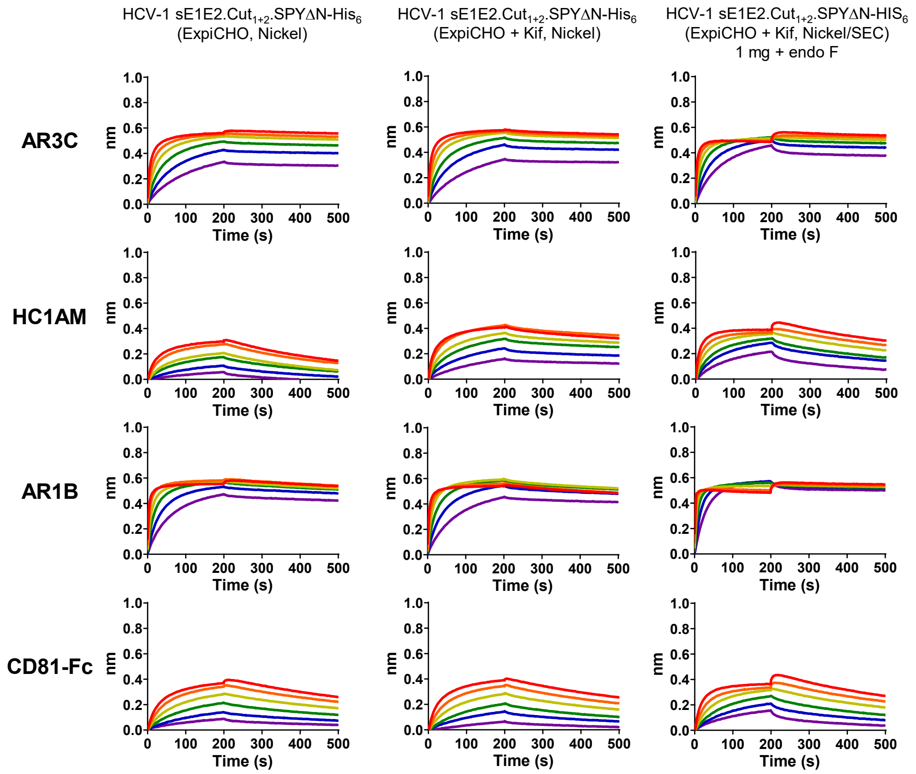


**$K_D$ values for unmodified and glycan-modified HCV-1 sE1E2.Cut$_{1+2}$.SPYΔN-His$_6$ binding to 10 human IgG antibodies and CD81-Fc (M) [a]**

| | AR4A | HCV1 | HPEC74 | RM2-01 | RM11-43 | AR5A | AR2A | AR3C | HC1AM | AR1B | CD81-Fc |
|---|---|---|---|---|---|---|---|---|---|---|---|
| HCV-1 sE1E2.Cut$_{1+2}$.SPYΔN-His$_6$ (ExpiCHO, Nickel) | <1.0E-12 | 1.91E-07 | 1.02E-10 | <1.0E-12 | 2.73E-09 | 4.13E-10 | 2.75E-09 | 2.02E-10 | 6.97E-08 | 3.76E-10 | 4.72E-08 |
| HCV-1 sE1E2.Cut$_{1+2}$.SPYΔN-His$_6$ (ExpiCHO + Kif, Nickel) | 3.20E-10 | 1.15E-09 | 4.13E-10 | 2.88E-11 | 2.99E-09 | 5.45E-10 | 3.69E-09 | 6.36E-10 | 8.10E-09 | 8.43E-10 | 4.86E-08 |
| HCV-1 sE1E2.Cut$_{1+2}$.SPYΔN-His$_6$ (ExpiCHO + Kif, Nickel/SEC) 1 mg + Endo F | <1.0E-12 | 3.93E-09 | <1.0E-12 | <1.0E-12 | 1.33E-09 | 9.49E-11 | 7.29E-10 | 1.33E-11 | 7.87E-09 | 9.39E-12 | 2.92E-08 |

[a] $K_D$ values were derived from bio-layer interferometry (BLI) using the binding equations describing a 1:1 interaction.
[b] "—" indicates cases where the peak signal value at the highest HCV1 sE1E2 dimer concentration is 0.2 or lower.

**Fig. S2. In vitro characterization of HCV sE1E2.Cut$_{1+2}$.SPYΔN-His$_6$ antigens.** (**a**) Amino acid sequences of genotype 1a HCV-1, genotype 3 UKNP3.1.2, and genotype 5 UKNP5.2.1 sE1E2.Cut$_{1+2}$.SPYΔN-His$_6$ antigens. The signal peptide, truncated E1 region, SpyTag002, furin cleavage motif, SpyCatcher002ΔN, short linkers, enzymatic site, and His$_6$ tag are highlighted in yellow, gray, green, red, orange, pink, blue, and cyan, respectively. (**b**) Reducing SDS-PAGE analysis of ExpiCHO-expressed HCV-1, UKNP3.1.2, and UKNP5.2.1 sE1E2.Cut$_{1+2}$.SPYΔN-His$_6$ antigens. SEC fraction numbers are labeled on the gel. (**c**) SEC profiles of Kif-treated and Kif/endo F-treated HCV-1 sE1E2.Cut$_{1+2}$.SPYΔN-His$_6$ antigens. In both cases, the HCV-1 sE1E2 scaffold was expressed in ExpiCHO cells in the presence of Kifunensine (Kif), followed by nickel and SEC purification. One milligram of Kif-treated antigen was processed using endo F1-3 and passed through SEC a second time to obtain the Kif/end F-treated antigen. (**d**) DSC profiles of Kif-treated and Kif/endo F-treated HCV-1 sE1E2.Cut$_{1+2}$.SPYΔN-His$_6$ antigens. Key thermal parameters such as $T_m$, $T_{onset}$, and $T_{\Delta 1/2}$ are labeled on the plot. (**e**) Compositional site-specific glycan analysis of HCV-1 sE1E2.Cut$_{1+2}$.SPYΔN-His$_6$ antigens obtained from two cell lines and two glycan modifications (four total samples). The graphs summarize quantitative mass spectrometric analysis of the glycan population present at individual N-linked glycosylation sites simplified into categories of glycans. The oligomannose-type glycan series (M9 to M5; Man9GlcNAc2 to Man5GlcNAc2) is colored green, afucosylated and fucosylated hybrid-type glycans (hybrid and F hybrid) are dashed pink, and complex glycans are grouped according to the number of antennae and presence of core fucosylation and are colored pink. Unoccupancy of an N-linked glycan site is represented in gray. Glycan sites that could not be determined are denoted as such. (**f**) ELISA analysis of unmodified, Kif-treated, and Kif/endo F-treated HCV-1 sE1E2.Cut$_{1+2}$.SPYΔN-His$_6$ antigens binding to 11 NAbs and CD81-Fc. Briefly, each well was coated with 0.1 μg of the appropriate antigen, and IgG or CD81-Fc was diluted in a 10-fold dilution series from a starting concentration of 10 μg/ml for all tested ligands. Error bars represent the difference between these duplicate values at each concentration tested for each sample. (**g**) BLI analysis of unmodified, Kif-treated, and Kif/endo F-treated HCV-1 sE1E2.Cut$_{1+2}$.SPYΔN-His$_6$ antigens binding to 10 human NAbs in the IgG form and CD81-Fc. Sensorgrams were obtained on an Octet RED96 with AHQ biosensors. A two-fold concentration gradient of antigen, starting at 1000 nM, was used in a dilution series of six. $K_D$ values derived from a 1:1 fitting model are summarized in a table.

**Figure S3**

a

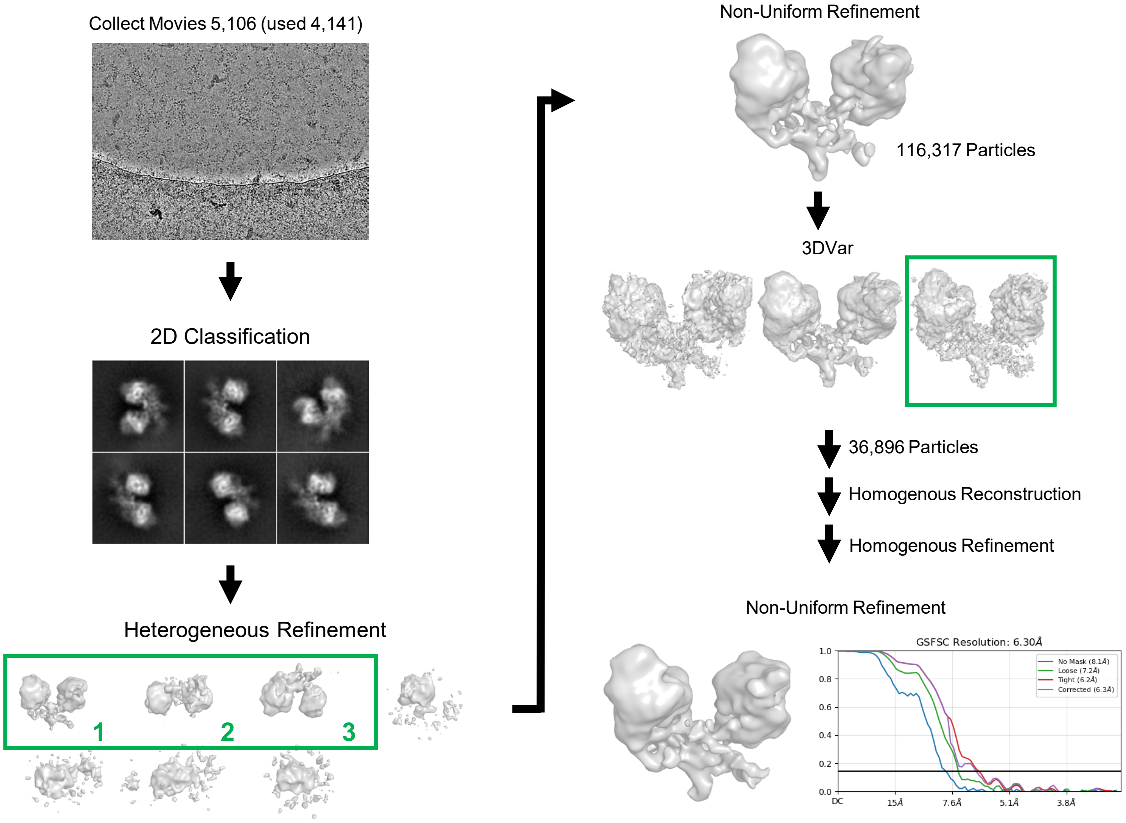

Collect Movies 5,106 (used 4,141)
2D Classification
Heterogeneous Refinement
1
2
3
Non-Uniform Refinement
116,317 Particles
3DVar
36,896 Particles
Homogenous Reconstruction
Homogenous Refinement
Non-Uniform Refinement
GSFSC Resolution: 6.30Å
No Mask (8.1Å)
Loose (7.2Å)
Tight (6.2Å)
Corrected (6.3Å)
1.0
0.8
0.6
0.4
0.2
0.0
DC
15Å
7.6Å
5.1Å
3.8Å


b

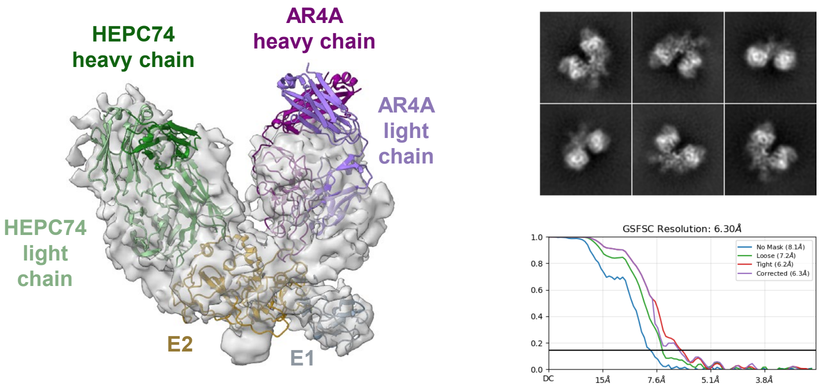

HEPC74 heavy chain
AR4A heavy chain
AR4A light chain
HEPC74 light chain
E2
E1
GSFSC Resolution: 6.30Å
No Mask (8.1Å)
Loose (7.2Å)
Tight (6.2Å)
Corrected (6.3Å)
1.0
0.8
0.6
0.4
0.2
0.0
DC
15Å
7.6Å
5.1Å
3.8Å

### Figure S3

c

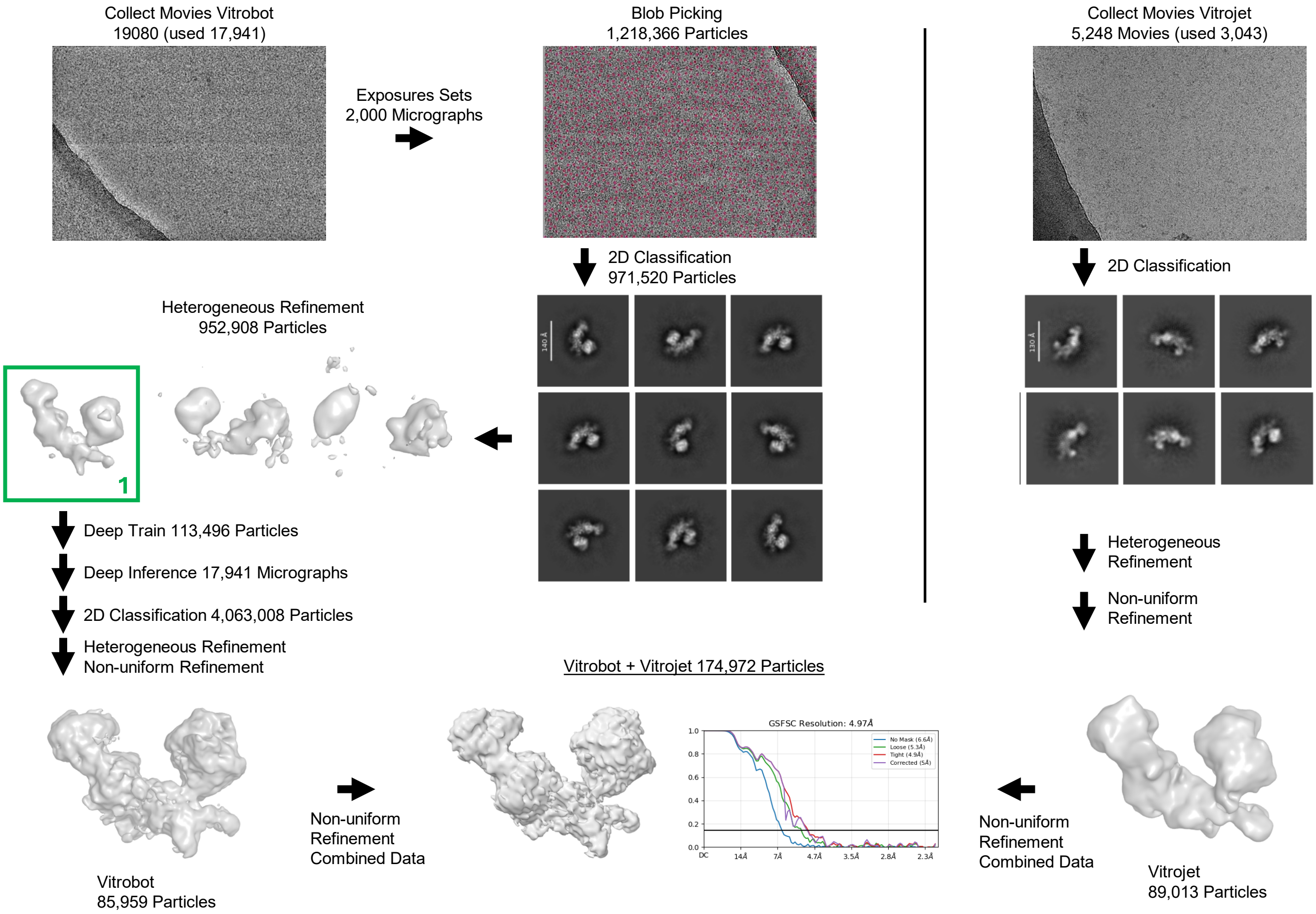
Collect Movies Vitrobot
19080 (used 17,941)
Exposures Sets
2,000 Micrographs
Blob Picking
1,218,366 Particles
Collect Movies Vitrojet
5,248 Movies (used 3,043)
2D Classification
971,520 Particles
2D Classification
Heterogeneous Refinement
952,908 Particles
1
Deep Train 113,496 Particles
Deep Inference 17,941 Micrographs
2D Classification 4,063,008 Particles
Heterogeneous Refinement
Non-uniform Refinement
Heterogeneous
Refinement
Non-uniform
Refinement
Vitrobot + Vitrojet 174,972 Particles
GSFSC Resolution: 4.97Å
No Mask (6.6Å)
Loose (5.3Å)
Tight (4.9Å)
Corrected (5Å)
Vitrobot
85,959 Particles
Non-uniform
Refinement
Combined Data
Non-uniform
Refinement
Combined Data
Vitrojet
89,013 Particles

d

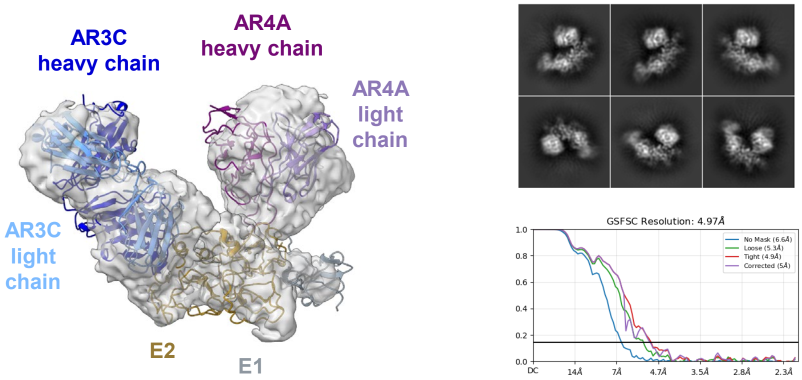
AR3C
heavy chain
AR4A
heavy chain
AR4A
light
chain
AR3C
light
chain
E2
E1
GSFSC Resolution: 4.97Å
No Mask (6.6Å)
Loose (5.3Å)
Tight (4.9Å)
Corrected (5Å)

### Figure S3

e

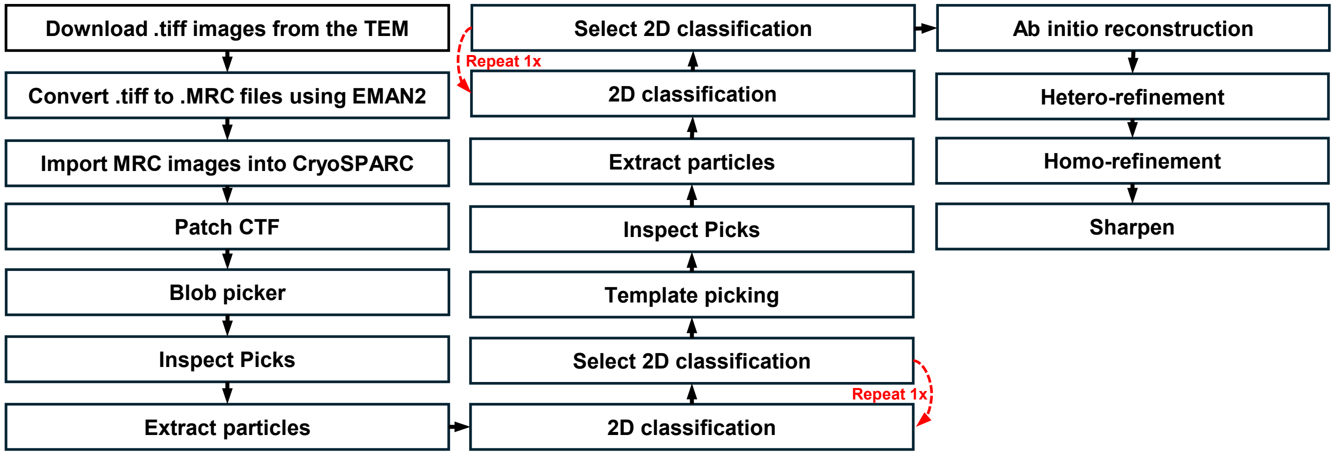


**Fig. S3. Structural characterization of HCV-1 sE1E2.Cut$_{1+2}$.SPYΔN-His$_6$ bound to NAbs by cryo-EM and nsEM.** For the cryo-EM analysis, images were collected on a Titan Krios 300 kV Cryo-transmission electron microscope (cryo-TEM) at the National Cancer Institute's (NCI) National Cryo-EM Facility, whereas for the nsEM analysis, images were collected on a Talos L120C TEM at 120kV at the Scripps Research Institute Core Microscopy Facility. (**a**) Cryo-EM processing of HCV-1 sE1E2.Cut$_{1+2}$.SPYΔN-His$_6$ in complex HEPC74 and AR4A Fabs. Key steps in the pipeline are shown. Classes selected for further processing from heterogeneous refinement jobs are boxed and numbered. (**b**) Cryo-EM reconstruction of HCV-1 sE1E2.Cut$_{1+2}$.SPYΔN-His$_6$ in complex HEPC74 and AR4A Fabs. The obtained cryo-EM map was rigid-body fitted with the antibody-bound structure of a scaffolded sE1E2 (PDB ID: 8FSJ). GSFSC curves and representative 2D classes are shown next to the 3D model. (**c**) Cryo-EM processing of HCV-1 sE1E2.Cut$_{1+2}$.SPYΔN-His$_6$ in complex AR3C and AR4A Fabs. Key steps in the pipeline are shown. Classes selected for further processing from heterogeneous refinement jobs are boxed and numbered. (**d**) Cryo-EM reconstruction of HCV-1 sE1E2.Cut$_{1+2}$.SPYΔN-His$_6$ in complex with AR3C and AR4A Fabs. The obtained cryo-EM map was rigid-body fitted with antibody-bound E1E2 and E2 structures (PDB IDs: 8FSJ and 4MWF, respectively). GSFSC curves and representative 2D classes are shown next to the 3D model. (**e**) nsEM processing pipeline used for epitope mapping of AR-series NAbs All steps in the pipeline are shown.

### Figure S4

a

**>HCV-1 sE1E2.Cut$_{1+2}$.SPYΔN-10GS-FR**

MGCSFSIFLLALLSCLTVPASAYQVRNSTGLYHVTNDCPNSSIVYEAADAILHTPGCVPCVREGNASRCWVAMTPTVATRDGKLPATQLRRHIDGSPRRHWTTQGCNC
SI**YPGH**GSGSVPTIVMVDAYKRYKRRRRRRETHVTGGSAGHTVSGFVSLLAPGAKQNVQLINTNGSWHLNSTALNCNDSLNTGWLAGLFYHHKFNSSGCPERLASCRP
LTDFDQGWGPISYANGSGPDQRPYCWHYPPKPCGIVPAKSVCGPVYCFTPSPVVVGTTDRSGAPTYSWGENDTDVFVLNNTRPPLGNWFGCTWMNSTGFTKVCGAPPC
VIGGAGNNTLHCPTDCFRKHPDATYSRCGSGPWITPRCLVDYPYRLWHYPCTINYTIFKIRMYVGGVEHRLEAACNWTRGERCDLEDRDRSELSPLLLTTTQWQVLPC
SFTTLPALSTGLIHLHQNIV**DVQYAS**DSATHIKFSKRDEDGRELAGATMELRDSSGKTISTWISDGHVKDFYLYPGKYTFVETAAPDGYEVATAITFTVNEQGQVTVN
GEATKGDAHTGGGGSGGGGSDIIKLLNEQVNKEMQSSNLYMSMSSWCYTHSLDGAGLFLFDHAAEEYEHAKKLIIFLNENNVPVQLTSISAPEHKFEGLTQIFQKAYE
HEQHISESINNIVDHAIKSKDHATFNFLQWYVAEQHEEEVLFKDILDKIELIGNENHGLYLADQYVKGIAKSRKS

**>HCV-1 sE1E2.Cut$_{1+2}$.SPYΔN-5GS-I3-01v9a-LD7-PADRE (or I3-01v9a-L7P)**

MGCSFSIFLLALLSCLTVPASAYQVRNSTGLYHVTNDCPNSSIVYEAADAILHTPGCVPCVREGNASRCWVAMTPTVATRDGKLPATQLRRHIDGSPRRHWTTQGCNC
SI**YPGH**GSGSVPTIVMVDAYKRYKRRRRRRETHVTGGSAGHTVSGFVSLLAPGAKQNVQLINTNGSWHLNSTALNCNDSLNTGWLAGLFYHHKFNSSGCPERLASCRP
LTDFDQGWGPISYANGSGPDQRPYCWHYPPKPCGIVPAKSVCGPVYCFTPSPVVVGTTDRSGAPTYSWGENDTDVFVLNNTRPPLGNWFGCTWMNSTGFTKVCGAPPC
VIGGAGNNTLHCPTDCFRKHPDATYSRCGSGPWITPRCLVDYPYRLWHYPCTINYTIFKIRMYVGGVEHRLEAACNWTRGERCDLEDRDRSELSPLLLTTTQWQVLPC
SFTTLPALSTGLIHLHQNIV**DVQYAS**DSATHIKFSKRDEDGRELAGATMELRDSSGKTISTWISDGHVKDFYLYPGKYTFVETAAPDGYEVATAITFTVNEQGQVTVN
GEATKGDAHTGGGGSAKLAEELQKKMEELFKKHHKIVAVLRANSVEEAKMKALAVFVGGVHLIEITFTVPDADTVIKELSFLKELGAIIGAGTVTSVEQCRKAVESGAE
FIVSPHLDEEISQFCKEKGVFYMPGVMTPTELVKAMKLGHTILKLFPGEVVGPQFVKAMKGPFPNVKFVPTGGVNLDNVCEWFKAGVLAVGVGSALVKGTIAEVAAKA
AAFVEKIRGCTEGGGGSSPAVDIGDRLDELEKALEALSAEDGHDDVGQRLESLLRRWNSRRADGSAKFVAAWTLKAAA

MGCSFSIFLLALLSCLTVPASA: Leader sequence
$^{257}$QLRRHID$^{263}$: Truncated putative fusion peptide (pFP, 272-285)-containing region
: SpyTag002
RRRRRR: Furin cleavage site motif
: SpyCatcher002ΔN
G: Linker; GSGS: Linker; GGGGS: Linker
AS: Enzymatic site
: FR or I3-01v9a-L7P

b

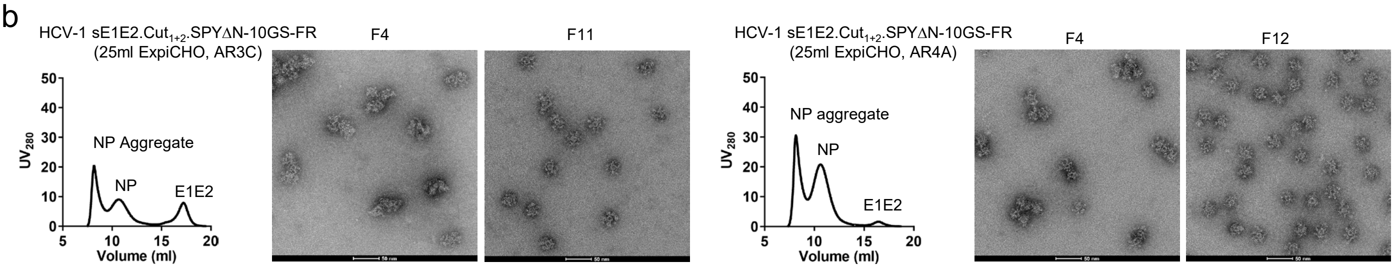


c

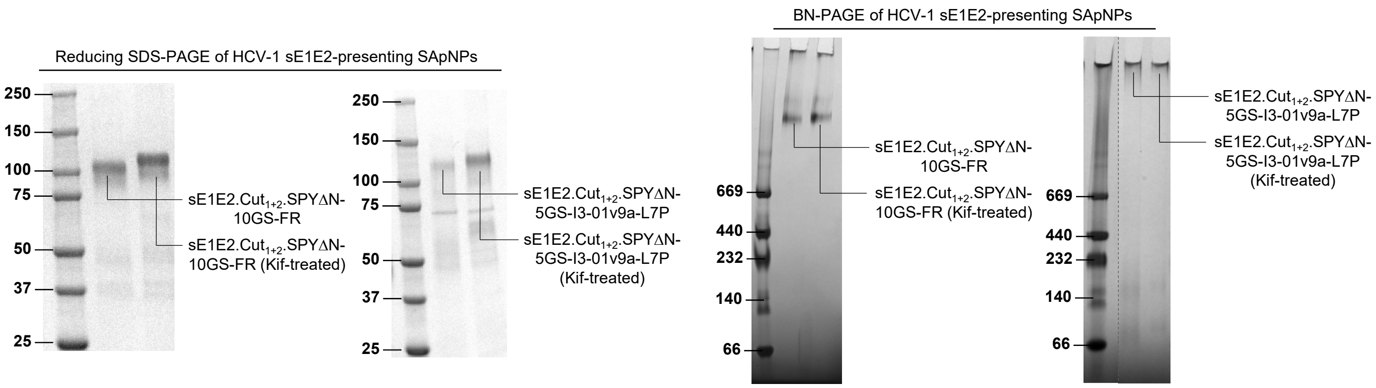

### Figure S4

d

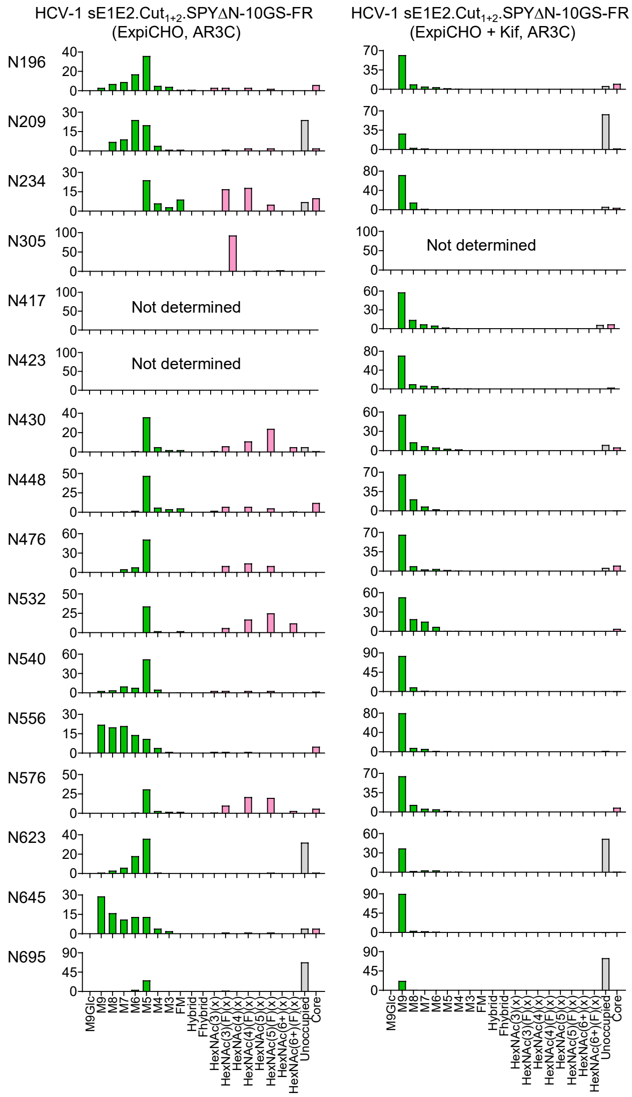


e

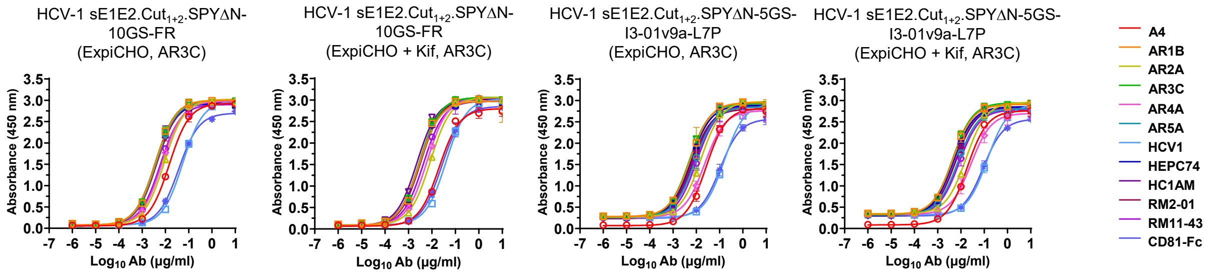


**HCV-1 sE1E2.Cut$_{1+2}$.SPYΔN-10GS-FR (ExpiCHO, AR3C)**

| A4 | | AR1B | | AR2A | | AR3C | | AR4A | | AR5A | |
|---|---|---|---|---|---|---|---|---|---|---|---|
| $EC_{50}$ | STD | $EC_{50}$ | STD | $EC_{50}$ | STD | $EC_{50}$ | STD | $EC_{50}$ | STD | $EC_{50}$ | STD |
| 0.0146 | 9.9E-05 | 0.00377 | 0.00012 | 0.00903 | 0.000924 | 0.00367 | 0.000394 | 0.00722 | 0.000223 | 0.004 | 0.0005 |

| HCV1 | | HEPC74 | | HC1AM | | RM2-01 | | RM11-43 | | CD81-Fc | |
|---|---|---|---|---|---|---|---|---|---|---|---|
| $EC_{50}$ | STD | $EC_{50}$ | STD | $EC_{50}$ | STD | $EC_{50}$ | STD | $EC_{50}$ | STD | $EC_{50}$ | STD |
| 0.056 | 0.00656 | 0.00355 | 0.000134 | 0.00356 | 0.00034 | 0.00417 | 0.000411 | 0.00596 | 0.000228 | 0.0373 | 0.00221 |

**HCV-1 sE1E2.Cut$_{1+2}$.SPYΔN-10GS-FR (ExpiCHO + Kif, AR3C)**

| A4 | | AR1B | | AR2A | | AR3C | | AR4A | | AR5A | |
|---|---|---|---|---|---|---|---|---|---|---|---|
| $EC_{50}$ | STD | $EC_{50}$ | STD | $EC_{50}$ | STD | $EC_{50}$ | STD | $EC_{50}$ | STD | $EC_{50}$ | STD |
| 0.0204 | 0.000849 | 0.00299 | 0.000194 | 0.00857 | 0.000124 | 0.00299 | 2.62E-05 | 0.00618 | 0.00024 | 0.00301 | 0.000156 |

| HCV1 | | HEPC74 | | HC1AM | | RM2-01 | | RM11-43 | | CD81-Fc | |
|---|---|---|---|---|---|---|---|---|---|---|---|
| $EC_{50}$ | STD | $EC_{50}$ | STD | $EC_{50}$ | STD | $EC_{50}$ | STD | $EC_{50}$ | STD | $EC_{50}$ | STD |
| 0.0399 | 0.00470 | 0.00296 | 9.12E-05 | 0.00231 | 1.91E-05 | 0.00331 | 4.81E-05 | 0.00437 | 0.000321 | 0.0263 | 0.00435 |

#### Figure S4

**HCV-1 sE1E2.Cut$_{1+2}$.SPYΔN-5GS-I3-01v9a-L7P (ExpiCHO, AR3C)**

| A4 | | AR1B | | AR2A | | AR3C | | AR4A | | AR5A | |
|---|---|---|---|---|---|---|---|---|---|---|---|
| $EC_{50}$ | STD | $EC_{50}$ | STD | $EC_{50}$ | STD | $EC_{50}$ | STD | $EC_{50}$ | STD | $EC_{50}$ | STD |
| 0.0264 | 0.0026 | 0.00559 | 0.00235 | 0.0157 | 0.00139 | 0.00503 | 0.000396 | 0.0229 | 0.00153 | 0.00858 | 0.00283 |

| HCV1 | | HEPC74 | | HC1AM | | RM2-01 | | RM11-43 | | CD81-Fc | |
|---|---|---|---|---|---|---|---|---|---|---|---|
| $EC_{50}$ | STD | $EC_{50}$ | STD | $EC_{50}$ | STD | $EC_{50}$ | STD | $EC_{50}$ | STD | $EC_{50}$ | STD |
| 0.159 | 0.00509 | 0.00570 | 0.00118 | 0.00738 | 0.00115 | 0.00603 | 0.000304 | 0.0101 | 0.00215 | 0.106 | 0.0164 |

**HCV-1 sE1E2.Cut$_{1+2}$.SPYΔN-5GS-I3-01v9a-L7P (ExpiCHO + Kif, AR3C)**

| A4 | | AR1B | | AR2A | | AR3C | | AR4A | | AR5A | |
|---|---|---|---|---|---|---|---|---|---|---|---|
| $EC_{50}$ | STD | $EC_{50}$ | STD | $EC_{50}$ | STD | $EC_{50}$ | STD | $EC_{50}$ | STD | $EC_{50}$ | STD |
| 0.0184 | 0.000665 | 0.00451 | 0.000643 | 0.0168 | 0.00122 | 0.00441 | 0.000414 | 0.0271 | 0.00521 | 0.00746 | 0.000114 |

| HCV1 | | HEPC74 | | HC1AM | | RM2-01 | | RM11-43 | | CD81-Fc | |
|---|---|---|---|---|---|---|---|---|---|---|---|
| $EC_{50}$ | STD | $EC_{50}$ | STD | $EC_{50}$ | STD | $EC_{50}$ | STD | $EC_{50}$ | STD | $EC_{50}$ | STD |
| 0.137 | 0.0187 | 0.00478 | 0.00019 | 0.00569 | 0.00072 | 0.00592 | 0.000538 | 0.00843 | 0.00151 | 0.0915 | 0.00812 |

f

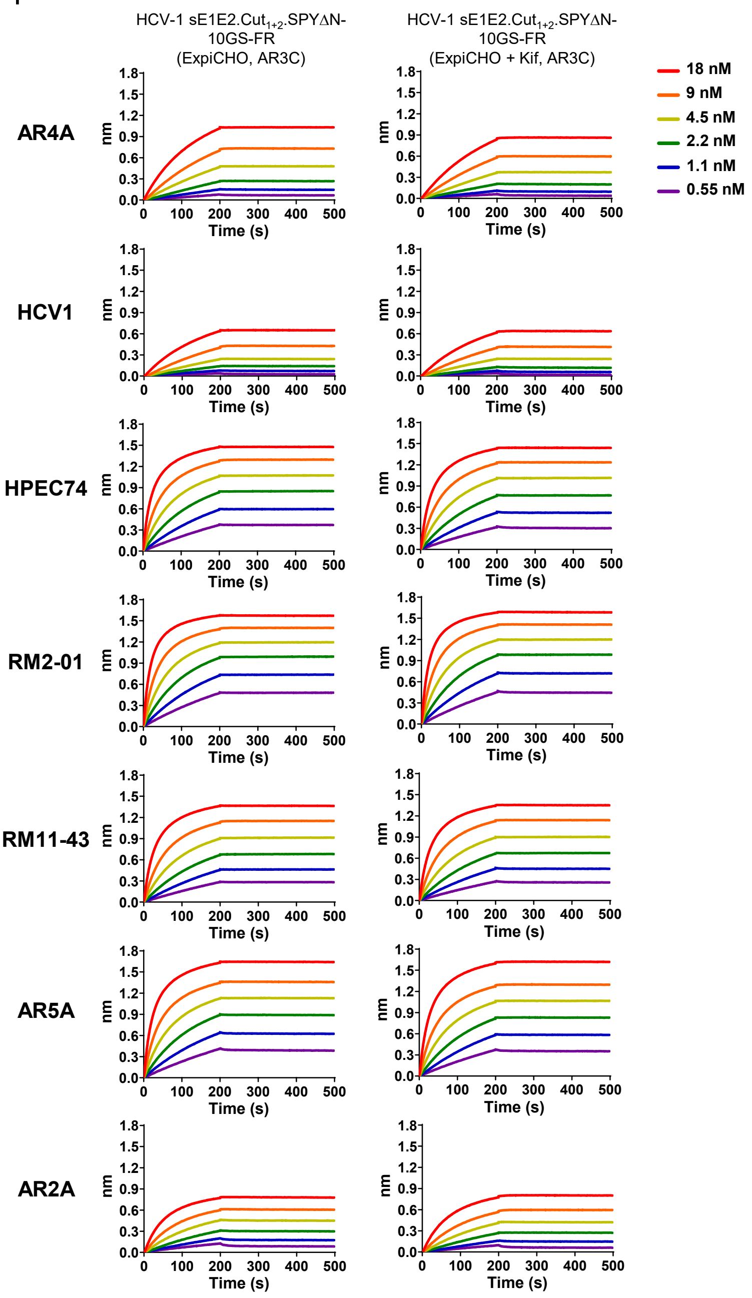

### Figure S4

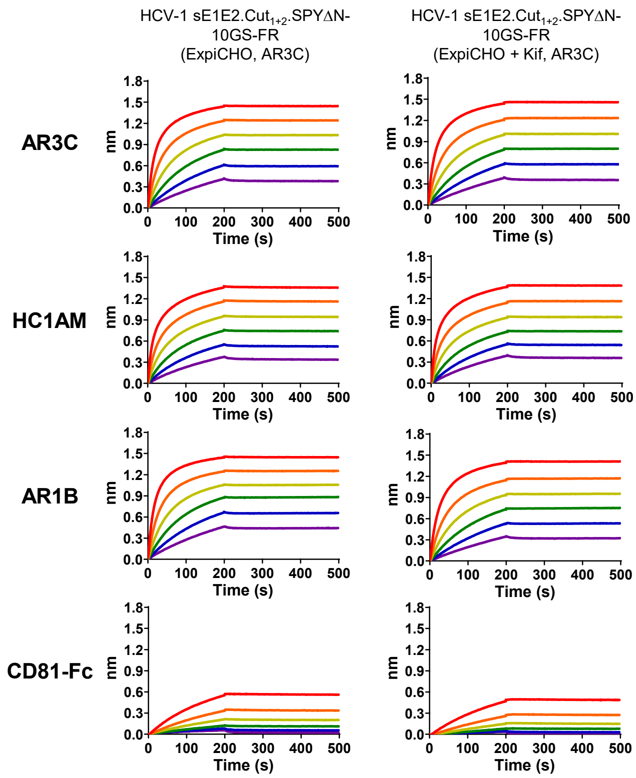


**$K_D$ values for unmodified and glycan-modified HCV-1 sE1E2-presenting FR SApNP binding to 10 human IgG antibodies and CD81-Fc (M) [a]**

| | AR4A | HCV1 | HPEC74 | RM2-01 | RM11-43 | AR5A | AR2A | AR3C | HC1AM | AR1B | CD81-Fc |
|---|---|---|---|---|---|---|---|---|---|---|---|
| HCV-1 sE1E2.Cut$_{1+2}$.SPYΔN-10GS-FR (ExpiCHO, AR3C) | <1.0E-12 | <1.0E-12 | <1.0E-12 | <1.0E-12 | 1.89E-11 | <1.0E-12 | 8.66E-11 | <1.0E-12 | <1.0E-12 | <1.0E-12 | 5.56E-10 |
| HCV1 sE1E2.Cut$_{1+2}$.SPYΔN-10GS-FR (ExpiCHO + Kif, AR3C) | <1.0E-12 | <1.0E-12 | <1.0E-12 | <1.0E-12 | <1.0E-12 | <1.0E-12 | <1.0E-12 | <1.0E-12 | <1.0E-12 | <1.0E-12 | <1.0E-12 |

[a] $K_D$ values were derived from bio-layer interferometry (BLI) using the binding equations describing a 1:1 interaction.
[b] "—" indicates cases where the peak signal value at the highest HCV-1 sE1E2 FR concentration is 0.2 or lower.

**Fig. S4. Rational design and in vitro characterization of HCV-1 sE1E2.Cut$_{1+2}$.SPYΔN-presenting SApNPs.** (**a**) Amino acid sequences of genotype 1a HCV-1 sE1E2.Cut$_{1+2}$.SPYΔN-presenting ferritin (FR) and I3-01v9a-LD7-PADRE (I3-01v9a-L7P) SApNPs. The signal peptide, truncated E1 region, SpyTag002, furin cleavage motif, SpyCatcher002ΔN, short linkers, enzymatic site, and SApNP backbone are highlighted in yellow, gray, green, red, orange, pink, blue, and cyan, respectively. (**b**) SEC and nsEM characterization of HCV-1 sE1E2.Cut$_{1+2}$.SPYΔN-10GS-FR expressed in 25-ml ExpiCHO cultures (without Kif-treatment) and IAC-purified using an AR3C (left) and AR4A (right) columns. SEC peaks corresponding to the NP aggregates, well-separate NPs, and unassembled subunits are labeled on the profile. Fractions 2 (~8.1 ml) and 11-12 (~10.6 ml) were analyzed by nsEM with the micrographs shown to the right of the respective SEC profile. (**c**) Reducing SDS-PAGE (left) and BN-PAGE (right) of HCV-1 sE1E2.Cut$_{1+2}$.SPYΔN-presenting FR and I3-01v9a SApNPs. Both unmodified and Kif-treated materials were analyzed and labeled on the gel. (**d**) Compositional site-specific glycan analysis of unmodified and Kif-treated HCV-1 sE1E2.Cut$_{1+2}$.SPYΔN-10GS-FR. Graphs summarize quantitative mass spectrometric analysis of the glycan populations at individual N-linked glycan sites, simplified into glycan categories. The oligomannose-type glycan series (M9 to M5; Man9GlcNAc2 to Man5GlcNAc2) is colored green; afucosylated and fucosylated hybrid-type glycans (hybrid and F hybrid) are dashed pink; and complex glycans are grouped by antenna number and presence of core fucosylation and are colored pink. Unoccupied N-linked glycan sites are shown in gray. Glycan sites that could not be determined are indicated as such. (**e**) ELISA analysis of unmodified and Kif-treated HCV-1 sE1E2.Cut$_{1+2}$.SPYΔN-presenting FR and I3-01v9a SApNPs binding to 11 NAbs and CD81-Fc. Briefly, each well was coated with 0.1 μg of the appropriate antigen, and IgG or CD81-Fc was diluted in a 10-fold dilution series from a starting concentration of 10 μg/ml for all tested ligands. Error bars represent the difference between duplicate values at each concentration tested for each sample. (**f**) BLI analysis of unmodified and Kif-treated, both AR3C-purified, HCV-1 sE1E2.Cut$_{1+2}$.SPYΔN-10GS-FR binding to 10 human NAbs in the IgG form and CD81-Fc. Sensorgrams were obtained on an Octet RED96 with AHQ biosensors. A two-fold concentration gradient of antigen, starting at 18 nM, was used in a dilution series of six. $K_D$ values derived from a 1:1 fitting model are summarized in a table. In (**c**)-(**f**), all SApNPs were produced in ExpiCHO cells, with or without Kif treatment, followed by AR3C purification.

### Figure. S5

a **Sera from mice immunized with HCV-1 sE1E2.Cut$_{1+2}$.SPYΔN vaccines against H77 HCVpp**

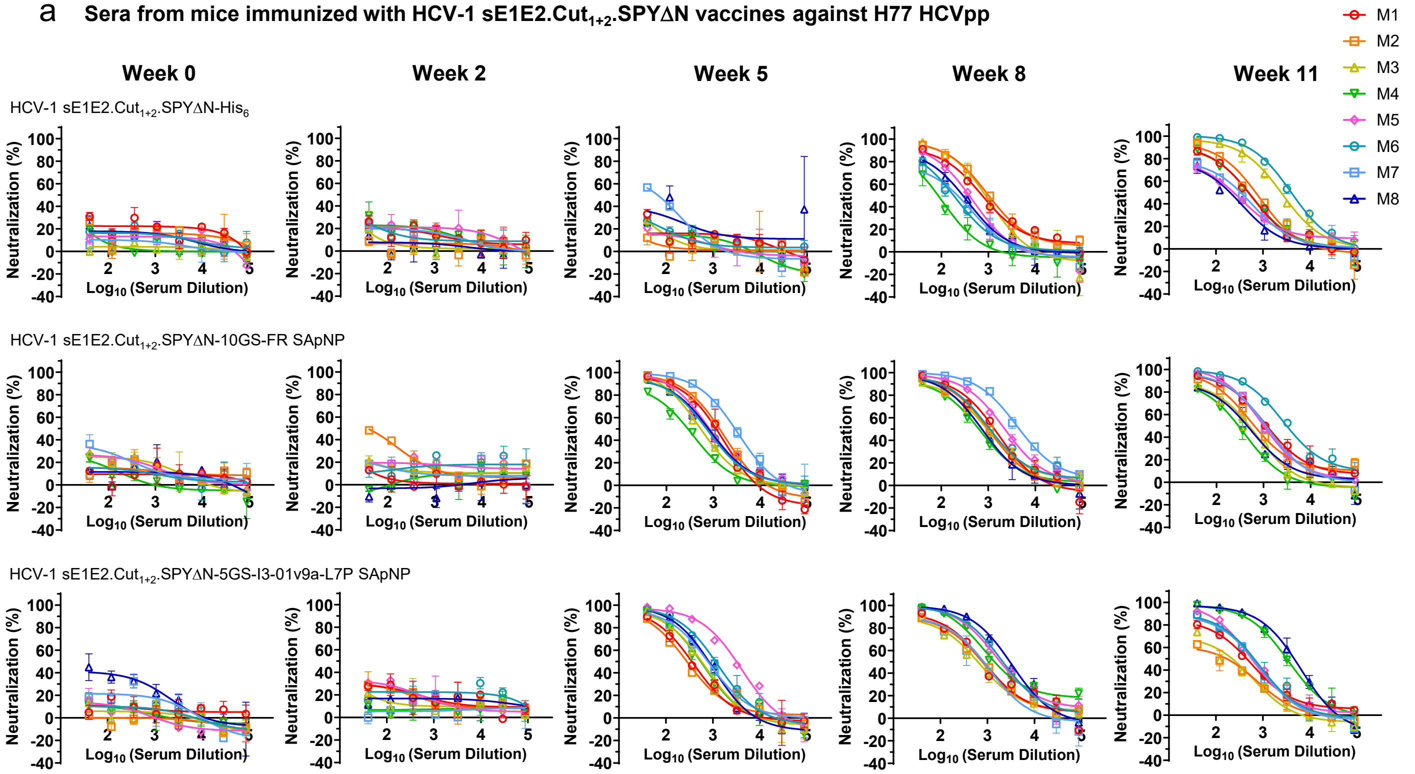


b **Mouse serum neutralizing ID$_{50}$ titers**

**Week 0**

| Antigen | ID50 titers (week 0) M1 | M2 | M3 | M4 | M5 | M6 | M7 | M8 | Geometric Mean |
|---|---|---|---|---|---|---|---|---|---|
| HCV-1 sE1E2.Cut$_{1+2}$.SPYΔN-His$_6$ | <40 | <40 | <40 | <40 | <40 | <40 | <40 | <40 | N/A |
| HCV-1 sE1E2.Cut$_{1+2}$.SPYΔN-10GS-FR SApNP | <40 | <40 | <40 | <40 | <40 | <40 | <40 | <40 | N/A |
| HCV-1 sE1E2.Cut$_{1+2}$.SPYΔN-5GS-I3-01v9a-L7P SApNP | <40 | <40 | <40 | <40 | <40 | <40 | <40 | 62.27 | N/A |

**Week 2**

| Antigen | ID50 titers (week 2) M1 | M2 | M3 | M4 | M5 | M6 | M7 | M8 | Geometric Mean |
|---|---|---|---|---|---|---|---|---|---|
| HCV-1 sE1E2.Cut$_{1+2}$.SPYΔN-His$_6$ | <40 | <40 | <40 | <40 | <40 | <40 | <40 | <40 | N/A |
| HCV-1 sE1E2.Cut$_{1+2}$.SPYΔN-10GS-FR SApNP | <40 | 59.42 | <40 | <40 | <40 | <40 | <40 | <40 | N/A |
| HCV-1 sE1E2.Cut$_{1+2}$.SPYΔN-5GS-I3-01v9a-L7P SApNP | <40 | <40 | <40 | <40 | <40 | <40 | <40 | <40 | N/A |

**Week 5**

| Antigen | ID50 titers (week 5) M1 | M2 | M3 | M4 | M5 | M6 | M7 | M8 | Geometric Mean |
|---|---|---|---|---|---|---|---|---|---|
| HCV-1 sE1E2.Cut$_{1+2}$.SPYΔN-His$_6$ | <40 | <40 | <40 | <40 | <40 | <40 | 66.68 | <40 | 17.8 |
| HCV-1 sE1E2.Cut$_{1+2}$.SPYΔN-10GS-FR SApNP | 942.9 | 1204 | 546.2 | 264.8 | 850.2 | 696.7 | 2697 | 755.6 | 816.8 |
| HCV-1 sE1E2.Cut$_{1+2}$.SPYΔN-5GS-I3-01v9a-L7P SApNP | 339 | 284.5 | 519.5 | 503.5 | 2739 | 1054 | 753.6 | 737.9 | 669.8 |

**Week 8**

| Antigen | ID50 titers (week 8) M1 | M2 | M3 | M4 | M5 | M6 | M7 | M8 | Geometric Mean |
|---|---|---|---|---|---|---|---|---|---|
| HCV-1 sE1E2.Cut$_{1+2}$.SPYΔN-His$_6$ | 749.4 | 1045 | 780.2 | 79.79 | 360.8 | 178.2 | 180.3 | 265 | 332.6 |
| HCV-1 sE1E2.Cut$_{1+2}$.SPYΔN-10GS-FR SApNP | 1225 | 962.5 | 932.7 | 615.9 | 2262 | 1136 | 4831 | 723.5 | 1253.2 |
| HCV-1 sE1E2.Cut$_{1+2}$.SPYΔN-5GS-I3-01v9a-L7P SApNP | 829.9 | 749.2 | 607.8 | 1779 | 2080 | 2187 | 690.5 | 2843 | 1251.1 |

**Week 11**

| Antigen | ID50 titers (week 11) M1 | M2 | M3 | M4 | M5 | M6 | M7 | M8 | Geometric Mean |
|---|---|---|---|---|---|---|---|---|---|
| HCV-1 sE1E2.Cut$_{1+2}$.SPYΔN-His$_6$ | 456.5 | 624.6 | 2376 | 461.2 | 231 | 4006 | 282.2 | 176.8 | 588.7 |
| HCV-1 sE1E2.Cut$_{1+2}$.SPYΔN-10GS-FR SApNP | 1240 | 698.1 | 432.3 | 262.5 | 1058 | 3396 | 1098 | 422.1 | 797.5 |
| HCV-1 sE1E2.Cut$_{1+2}$.SPYΔN-5GS-I3-01v9a-L7P SApNP | 423.6 | 141.9 | 163.5 | 3087 | 586.8 | 575 | 518 | 3912 | 616.0 |

**Statistical analysis**

| One-way ANOVA with Tukey's multiple comparisons test (w5) | Statistics | Adjusted P Value |
|---|---|---|
| HCV-1 sE1E2.Cut$_{1+2}$.SPYΔN-His$_6$ vs. HCV-1 sE1E2.Cut$_{1+2}$.SPYΔN-10GS-FR SApNP | * | 0.0146 |
| HCV-1 sE1E2.Cut$_{1+2}$.SPYΔN-His$_6$ vs. HCV-1 sE1E2.Cut$_{1+2}$.SPYΔN-5GS-I3-01v9a-L7P SApNP | * | 0.0356 |
| HCV-1 sE1E2.Cut$_{1+2}$.SPYΔN-10GS-FR SApNP vs. HCV-1 sE1E2.Cut$_{1+2}$.SPYΔN-5GS-I3-01v9a-L7P SApNP | ns | 0.9125 |

| One-way ANOVA with Tukey's multiple comparisons test (w8) | Statistics | Adjusted P Value |
|---|---|---|
| HCV-1 sE1E2.Cut$_{1+2}$.SPYΔN-His$_6$ vs. HCV-1 sE1E2.Cut$_{1+2}$.SPYΔN-10GS-FR SApNP | ns | 0.0738 |
| HCV-1 sE1E2.Cut$_{1+2}$.SPYΔN-His$_6$ vs. HCV-1 sE1E2.Cut$_{1+2}$.SPYΔN-5GS-I3-01v9a-L7P SApNP | ns | 0.1161 |
| HCV-1 sE1E2.Cut$_{1+2}$.SPYΔN-10GS-FR SApNP vs. HCV-1 sE1E2.Cut$_{1+2}$.SPYΔN-5GS-I3-01v9a-L7P SApNP | ns | 0.9695 |

| One-way ANOVA with Tukey's multiple comparisons test (w11) | Statistics | Adjusted P Value |
|---|---|---|
| HCV-1 sE1E2.Cut$_{1+2}$.SPYΔN-His$_6$ vs. HCV-1 sE1E2.Cut$_{1+2}$.SPYΔN-10GS-FR SApNP | ns | >0.9999 |
| HCV-1 sE1E2.Cut$_{1+2}$.SPYΔN-His$_6$ vs. HCV-1 sE1E2.Cut$_{1+2}$.SPYΔN-5GS-I3-01v9a-L7P SApNP | ns | 0.9872 |
| HCV-1 sE1E2.Cut$_{1+2}$.SPYΔN-10GS-FR SApNP vs. HCV-1 sE1E2.Cut$_{1+2}$.SPYΔN-5GS-I3-01v9a-L7P SApNP | ns | 0.987 |

#### C Week-11 sera from mice immunized with HCV-1 sE1E2.Cut$_{1+2}$.SPYΔN vaccines against heterologous HCVpps

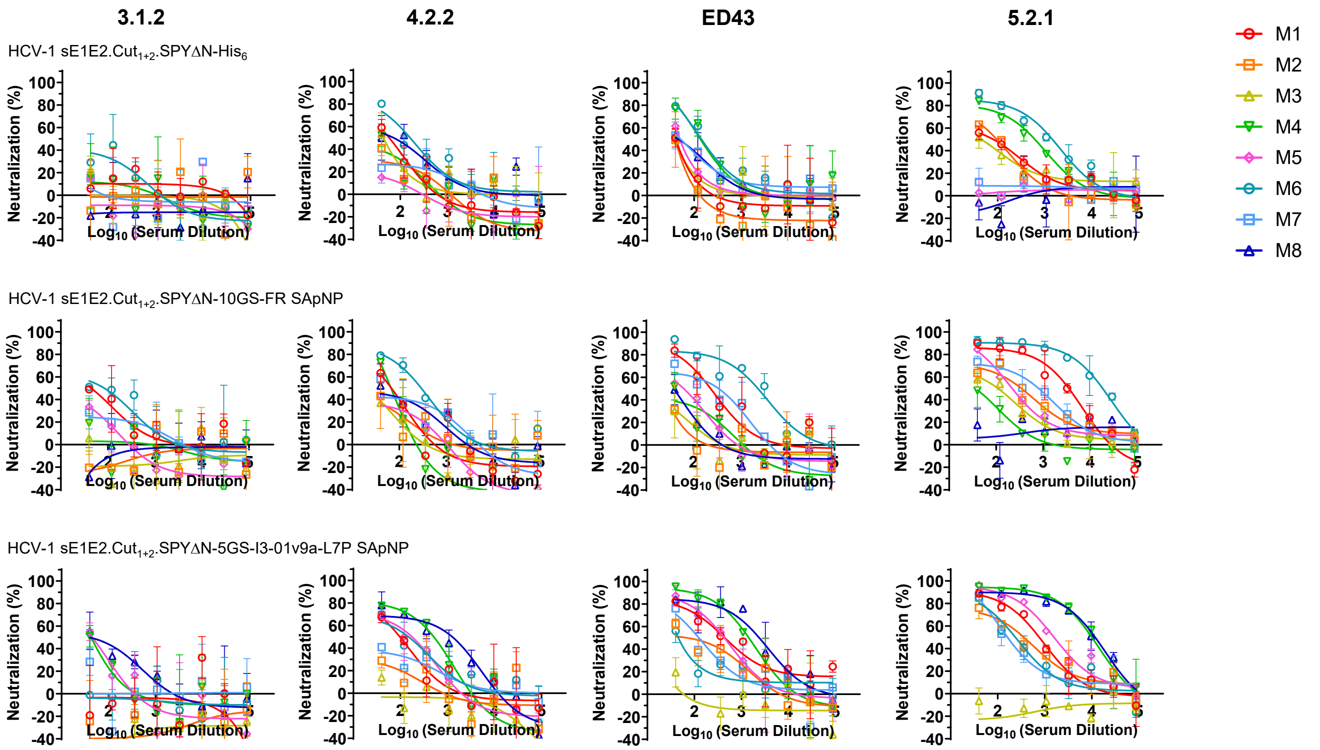


#### d Mouse serum neutralizing ID$_{50}$ titers

Against 3.1.2 HCVpp

| Antigen | ID50 titers (week 11) | | | | | | | | Geometric Mean |
|---|---|---|---|---|---|---|---|---|---|
| | M1 | M2 | M3 | M4 | M5 | M6 | M7 | M8 | |
| HCV-1 sE1E2.Cut$_{1+2}$.SPYΔN-His$_6$ | <40 | <40 | <40 | <40 | <40 | <40 | <40 | <40 | N/A |
| HCV-1 sE1E2.Cut$_{1+2}$.SPYΔN-10GS-FR SApNP | 51.74 | <40 | <40 | <40 | <40 | 89.24 | <40 | <40 | N/A |
| HCV-1 sE1E2.Cut$_{1+2}$.SPYΔN-5GS-I3-01v9a-L7P SApNP | <40 | <40 | <40 | <40 | <40 | <40 | <40 | 65.6 | N/A |

Against 4.2.2 HCVpp

| Antigen | ID50 titers (week 11) | | | | | | | | Geometric Mean |
|---|---|---|---|---|---|---|---|---|---|
| | M1 | M2 | M3 | M4 | M5 | M6 | M7 | M8 | |
| HCV-1 sE1E2.Cut$_{1+2}$.SPYΔN-His$_6$ | 49.96 | <40 | <40 | <40 | <40 | 152.9 | <40 | 87.21 | 37.0 |
| HCV-1 sE1E2.Cut$_{1+2}$.SPYΔN-10GS-FR SApNP | 55.29 | <40 | <40 | 44.08 | <40 | 257.8 | 71.24 | 61.22 | 52.1 |
| HCV-1 sE1E2.Cut$_{1+2}$.SPYΔN-5GS-I3-01v9a-L7P SApNP | 88.57 | <40 | <40 | 339.9 | 128.6 | 124 | <40 | 549.5 | 69.9 |

Against ED43 HCVpp

| Antigen | ID50 titers (week 11) | | | | | | | | Geometric Mean |
|---|---|---|---|---|---|---|---|---|---|
| | M1 | M2 | M3 | M4 | M5 | M6 | M7 | M8 | |
| HCV-1 sE1E2.Cut$_{1+2}$.SPYΔN-His$_6$ | <40 | <40 | <40 | 134 | 46.13 | 148.2 | 60.81 | 55.09 | 52.4 |
| HCV-1 sE1E2.Cut$_{1+2}$.SPYΔN-10GS-FR SApNP | 213.5 | <40 | <40 | <40 | 57.86 | 1876 | 195.9 | <40 | 76.0 |
| HCV-1 sE1E2.Cut$_{1+2}$.SPYΔN-5GS-I3-01v9a-L7P SApNP | 469.6 | 103.4 | <40 | 1225 | 383 | 51.91 | 124 | 2064 | 145.0 |

Against 5.2.1 HCVpp

| Antigen | ID50 titers (week 11) | | | | | | | | Geometric Mean |
|---|---|---|---|---|---|---|---|---|---|
| | M1 | M2 | M3 | M4 | M5 | M6 | M7 | M8 | |
| HCV-1 sE1E2.Cut$_{1+2}$.SPYΔN-His$_6$ | 92.62 | 90.54 | 65.78 | 475 | <40 | 1121 | <40 | <40 | 58.7 |
| HCV-1 sE1E2.Cut$_{1+2}$.SPYΔN-10GS-FR SApNP | 2327 | 303.6 | 112.6 | 43.03 | 239.8 | 15202 | 553.6 | <40 | 285.7 |
| HCV-1 sE1E2.Cut$_{1+2}$.SPYΔN-5GS-I3-01v9a-L7P SApNP | 689.2 | 308.3 | <40 | 8928 | 1576 | 229.9 | 190.4 | 9417 | 1030.3 |

#### Statistical analysis

Against 4.2.2 HCVpp

| One-way ANOVA with Tukey's multiple comparisons test (w11) | Statistics | Adjusted P Value |
|---|---|---|
| HCV-1 sE1E2.Cut$_{1+2}$.SPYΔN-His$_6$ vs. HCV-1 sE1E2.Cut$_{1+2}$.SPYΔN-10GS-FR SApNP | ns | 0.9457 |
| HCV-1 sE1E2.Cut$_{1+2}$.SPYΔN-His$_6$ vs. HCV-1 sE1E2.Cut$_{1+2}$.SPYΔN-5GS-I3-01v9a-L7P SApNP | ns | 0.1944 |
| V-1 sE1E2.Cut$_{1+2}$.SPYΔN-10GS-FR SApNP vs. HCV-1 sE1E2.Cut$_{1+2}$.SPYΔN-5GS-I3-01v9a-L7P SApNP | ns | 0.3205 |

Against ED43 HCVpp

| One-way ANOVA with Tukey's multiple comparisons test (w11) | Statistics | Adjusted P Value |
|---|---|---|
| HCV-1 sE1E2.Cut$_{1+2}$.SPYΔN-His$_6$ vs. HCV-1 sE1E2.Cut$_{1+2}$.SPYΔN-10GS-FR SApNP | ns | 0.6732 |
| HCV-1 sE1E2.Cut$_{1+2}$.SPYΔN-His$_6$ vs. HCV-1 sE1E2.Cut$_{1+2}$.SPYΔN-5GS-I3-01v9a-L7P SApNP | ns | 0.2146 |
| V-1 sE1E2.Cut$_{1+2}$.SPYΔN-10GS-FR SApNP vs. HCV-1 sE1E2.Cut$_{1+2}$.SPYΔN-5GS-I3-01v9a-L7P SApNP | ns | 0.6567 |

Against 5.2.1 HCVpp

| One-way ANOVA with Tukey's multiple comparisons test (w11) | Statistics | Adjusted P Value |
|---|---|---|
| HCV-1 sE1E2.Cut$_{1+2}$.SPYΔN-His$_6$ vs. HCV-1 sE1E2.Cut$_{1+2}$.SPYΔN-10GS-FR SApNP | ns | 0.5781 |
| HCV-1 sE1E2.Cut$_{1+2}$.SPYΔN-His$_6$ vs. HCV-1 sE1E2.Cut$_{1+2}$.SPYΔN-5GS-I3-01v9a-L7P SApNP | ns | 0.4071 |
| V-1 sE1E2.Cut$_{1+2}$.SPYΔN-10GS-FR SApNP vs. HCV-1 sE1E2.Cut$_{1+2}$.SPYΔN-5GS-I3-01v9a-L7P SApNP | ns | 0.9384 |

### Figure. S5

**e** Sera from mice immunized with glycan-modified HCV-1 sE1E2.Cut$_{1+2}$.SPYΔN-His$_6$ dimer vaccines against H77 HCVpp

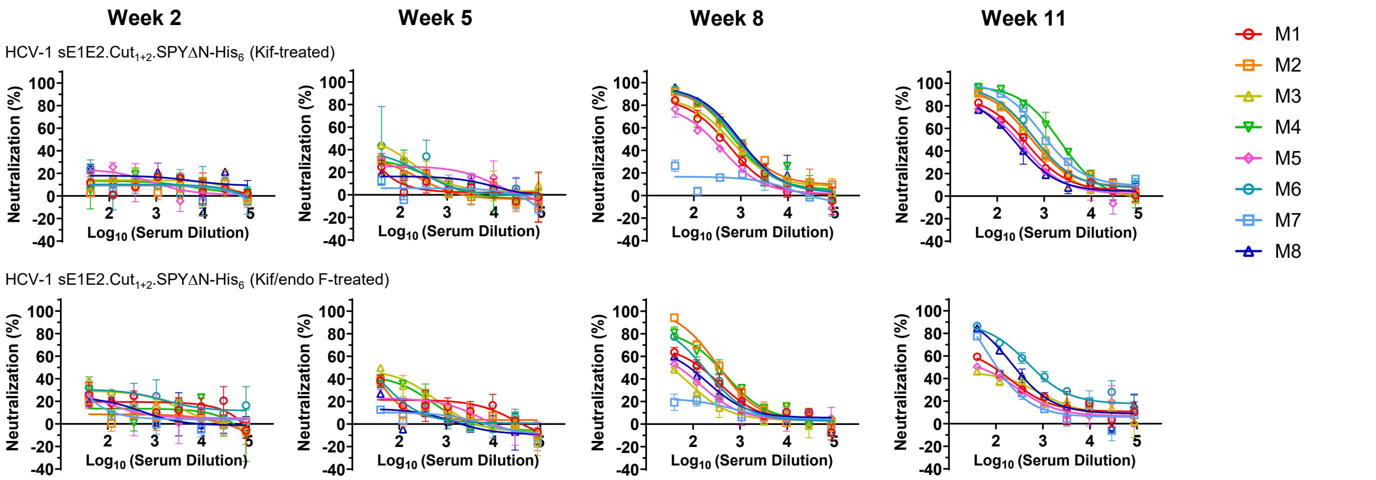


**f** Mouse serum neutralizing ID$_{50}$ titers

**Week 2**

| Antigen | ID50 titers (week 2) | | | | | | | | Geometric Mean |
|---|---|---|---|---|---|---|---|---|---|
| | M1 | M2 | M3 | M4 | M5 | M6 | M7 | M8 | |
| HCV-1 sE1E2.Cut$_{1+2}$.SPYΔN-His$_6$ | <40 | <40 | <40 | <40 | <40 | <40 | <40 | <40 | N/A |
| HCV-1 sE1E2.Cut$_{1+2}$.SPYΔN-His$_6$ (Kif-treated) | <40 | <40 | <40 | <40 | <40 | <40 | <40 | <40 | N/A |
| HCV-1 sE1E2.Cut$_{1+2}$.SPYΔN-His$_6$ (Kif/endo F-treated) | <40 | <40 | <40 | <40 | <40 | <40 | <40 | <40 | N/A |

**Week 5**

| Antigen | ID50 titers (week 5) | | | | | | | | Geometric Mean |
|---|---|---|---|---|---|---|---|---|---|
| | M1 | M2 | M3 | M4 | M5 | M6 | M7 | M8 | |
| HCV-1 sE1E2.Cut$_{1+2}$.SPYΔN-His$_6$ | <40 | <40 | <40 | <40 | <40 | <40 | 66.68 | <40 | 17.8 |
| HCV-1 sE1E2.Cut$_{1+2}$.SPYΔN-His$_6$ (Kif-treated) | <40 | <40 | 45.36 | <40 | <40 | <40 | <40 | <40 | 19.2 |
| HCV-1 sE1E2.Cut$_{1+2}$.SPYΔN-His$_6$ (Kif/endo F-treated) | <40 | <40 | 58.31 | 43.14 | <40 | <40 | <40 | <40 | 23.2 |

**Week 8**

| Antigen | ID50 titers (week 8) | | | | | | | | Geometric Mean |
|---|---|---|---|---|---|---|---|---|---|
| | M1 | M2 | M3 | M4 | M5 | M6 | M7 | M8 | |
| HCV-1 sE1E2.Cut$_{1+2}$.SPYΔN-His$_6$ | 749.4 | 1045 | 780.2 | 79.79 | 360.8 | 178.2 | 180.3 | 265 | 332.6 |
| HCV-1 sE1E2.Cut$_{1+2}$.SPYΔN-His$_6$ (Kif-treated) | 375.3 | 904.9 | 559.6 | 739 | 217.3 | 831.8 | <40 | 922.4 | 368.7 |
| HCV-1 sE1E2.Cut$_{1+2}$.SPYΔN-His$_6$ (Kif/endo F-treated) | 137.4 | 378.8 | 44.62 | 315.3 | 62.33 | 166.2 | <40 | 91.48 | 101.0 |

**Week 11**

| Antigen | ID50 titers (week 11) | | | | | | | | Geometric Mean |
|---|---|---|---|---|---|---|---|---|---|
| | M1 | M2 | M3 | M4 | M5 | M6 | M7 | M8 | |
| HCV-1 sE1E2.Cut$_{1+2}$.SPYΔN-His$_6$ | 456.5 | 624.6 | 2376 | 461.2 | 231 | 4006 | 282.2 | 176.8 | 588.7 |
| HCV-1 sE1E2.Cut$_{1+2}$.SPYΔN-His$_6$ (Kif-treated) | 345.2 | 585.8 | 602.3 | 1948 | 249.6 | 691.7 | 1249 | 216.5 | 569.6 |
| HCV-1 sE1E2.Cut$_{1+2}$.SPYΔN-His$_6$ (Kif/endo F-treated) | 103.2 | N/A | 74.37 | N/A | 73.76 | 669.4 | 119 | 276.3 | 152.3 |

#### Statistical analysis

| One-way ANOVA with Tukey's multiple comparisons test (w5) | Statistics | Adjusted P Value |
|---|---|---|
| HCV-1 sE1E2.Cut$_{1+2}$.SPYΔN-His$_6$ vs. HCV-1 sE1E2.Cut$_{1+2}$.SPYΔN-His$_6$ (Kif-treated) | ns | 0.9981 |
| HCV-1 sE1E2.Cut$_{1+2}$.SPYΔN-His$_6$ vs. HCV-1 sE1E2.Cut$_{1+2}$.SPYΔN-His$_6$ (Kif/endo F-treated) | ns | 0.8533 |
| HCV-1 sE1E2.Cut$_{1+2}$.SPYΔN-His$_6$ (Kif-treated) vs. HCV-1 sE1E2.Cut$_{1+2}$.SPYΔN-His$_6$ (Kif/endo F-treated) | ns | 0.8229 |

| One-way ANOVA with Tukey's multiple comparisons test (w8) | Statistics | Adjusted P Value |
|---|---|---|
| HCV-1 sE1E2.Cut$_{1+2}$.SPYΔN-His$_6$ vs. HCV-1 sE1E2.Cut$_{1+2}$.SPYΔN-His$_6$ (Kif-treated) | ns | 0.713 |
| HCV-1 sE1E2.Cut$_{1+2}$.SPYΔN-His$_6$ vs. HCV-1 sE1E2.Cut$_{1+2}$.SPYΔN-His$_6$ (Kif/endo F-treated) | ns | 0.1204 |
| HCV-1 sE1E2.Cut$_{1+2}$.SPYΔN-His$_6$ (Kif-treated) vs. HCV-1 sE1E2.Cut$_{1+2}$.SPYΔN-His$_6$ (Kif/endo F-treated) | * | 0.0243 |

| One-way ANOVA with Tukey's multiple comparisons test (w11) | Statistics | Adjusted P Value |
|---|---|---|
| HCV-1 sE1E2.Cut$_{1+2}$.SPYΔN-His$_6$ vs. HCV-1 sE1E2.Cut$_{1+2}$.SPYΔN-His$_6$ (Kif-treated) | ns | 0.7429 |
| HCV-1 sE1E2.Cut$_{1+2}$.SPYΔN-His$_6$ vs. HCV-1 sE1E2.Cut$_{1+2}$.SPYΔN-His$_6$ (Kif/endo F-treated) | ns | 0.222 |
| HCV-1 sE1E2.Cut$_{1+2}$.SPYΔN-His$_6$ (Kif-treated) vs. HCV-1 sE1E2.Cut$_{1+2}$.SPYΔN-His$_6$ (Kif/endo F-treated) | ns | 0.562 |

**Figure. S5**

#### g Sera from mice immunized with glycan-modified HCV-1 sE1E2.Cut$_{1+2}$.SPYΔN-His$_6$ dimer vaccines against heterologous HCVpps

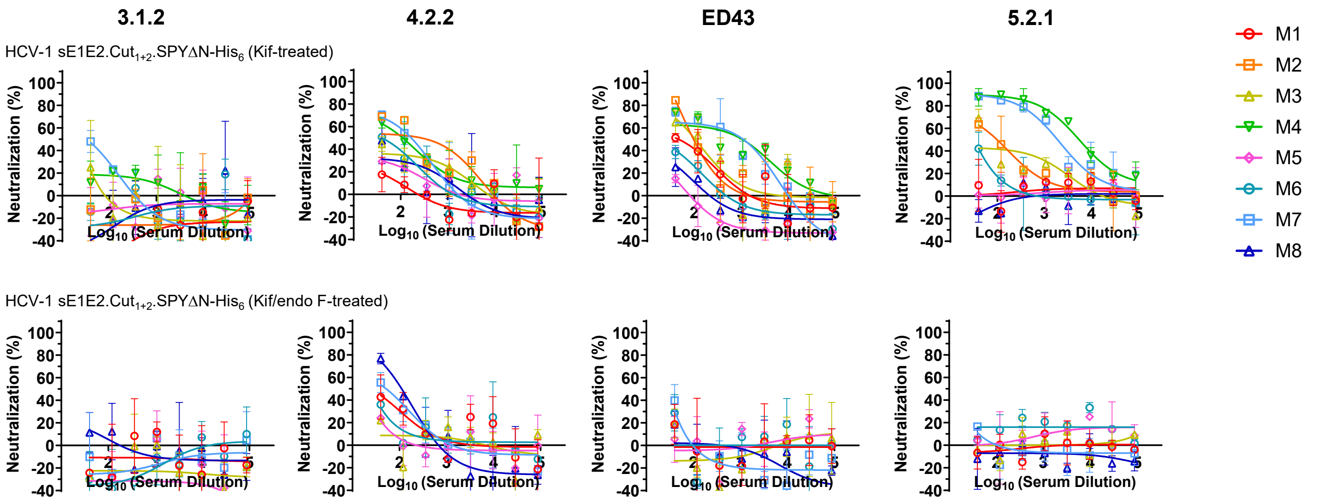


#### h Mouse serum neutralizing ID$_{50}$ titers

Against 3.1.2 HCVpp

| Antigen | ID50 titers (week 11) M1 | M2 | M3 | M4 | M5 | M6 | M7 | M8 | Geometric Mean |
|---|---|---|---|---|---|---|---|---|---|
| HCV-1 sE1E2.Cut$_{1+2}$.SPYΔN-His$_6$ | <40 | <40 | <40 | <40 | <40 | <40 | <40 | <40 | N/A |
| HCV-1 sE1E2.Cut$_{1+2}$.SPYΔN-His$_6$ (Kif-treated) | <40 | <40 | <40 | <40 | <40 | <40 | <40 | <40 | N/A |
| HCV-1 sE1E2.Cut$_{1+2}$.SPYΔN-His$_6$ (Kif/endo F-treated) | <40 | N/A | <40 | N/A | <40 | <40 | <40 | <40 | N/A |

Against 4.2.2 HCVpp

| Antigen | ID50 titers (week 11) M1 | M2 | M3 | M4 | M5 | M6 | M7 | M8 | Geometric Mean |
|---|---|---|---|---|---|---|---|---|---|
| HCV-1 sE1E2.Cut$_{1+2}$.SPYΔN-His$_6$ | 49.96 | <40 | <40 | <40 | <40 | 152.9 | <40 | 87.21 | 37.0 |
| HCV-1 sE1E2.Cut$_{1+2}$.SPYΔN-His$_6$ (Kif-treated) | <40 | 181.1 | 49.35 | 112.5 | <40 | 50.97 | 124.1 | <40 | 50.1 |
| HCV-1 sE1E2.Cut$_{1+2}$.SPYΔN-His$_6$ (Kif/endo F-treated) | <40 | N/A | <40 | N/A | <40 | <40 | 53.57 | 85.8 | 24.4 |

Against ED43 HCVpp

| Antigen | ID50 titers (week 11) M1 | M2 | M3 | M4 | M5 | M6 | M7 | M8 | Geometric Mean |
|---|---|---|---|---|---|---|---|---|---|
| HCV-1 sE1E2.Cut$_{1+2}$.SPYΔN-His$_6$ | <40 | <40 | <40 | 134 | 46.13 | 148.2 | 60.81 | 55.09 | 52.4 |
| HCV-1 sE1E2.Cut$_{1+2}$.SPYΔN-His$_6$ (Kif-treated) | 57.67 | 99.66 | 102.2 | 413.4 | <40 | <40 | 443.3 | 12.23 | 56.0 |
| HCV-1 sE1E2.Cut$_{1+2}$.SPYΔN-His$_6$ (Kif/endo F-treated) | <40 | N/A | <40 | N/A | <40 | <40 | <40 | <40 | 4.2 |

Against 5.2.1 HCVpp

| Antigen | ID50 titers (week 11) M1 | M2 | M3 | M4 | M5 | M6 | M7 | M8 | Geometric Mean |
|---|---|---|---|---|---|---|---|---|---|
| HCV-1 sE1E2.Cut$_{1+2}$.SPYΔN-His$_6$ | 92.62 | 90.54 | 65.78 | 475 | <40 | 1121 | <40 | <40 | 58.7 |
| HCV-1 sE1E2.Cut$_{1+2}$.SPYΔN-His$_6$ (Kif-treated) | <40 | 99.52 | 81.08 | 5314 | <40 | <40 | 1788 | <40 | 58.7 |
| HCV-1 sE1E2.Cut$_{1+2}$.SPYΔN-His$_6$ (Kif/endo F-treated) | <40 | N/A | <40 | N/A | <40 | <40 | <40 | <40 | 3.2 |

##### Statistical analysis

Against 4.2.2 HCVpp

| One-way ANOVA with Tukey's multiple comparisons test (w11) | Statistics | Adjusted P Value |
|---|---|---|
| HCV-1 sE1E2.Cut$_{1+2}$.SPYΔN-His$_6$ vs. HCV-1 sE1E2.Cut$_{1+2}$.SPYΔN-His$_6$ (Kif-treated) | ns | 0.6751 |
| HCV-1 sE1E2.Cut$_{1+2}$.SPYΔN-His$_6$ vs. HCV-1 sE1E2.Cut$_{1+2}$.SPYΔN-His$_6$ (Kif/endo F-treated) | ns | 0.802 |
| HCV-1 sE1E2.Cut$_{1+2}$.SPYΔN-His$_6$ (Kif-treated) vs. HCV-1 sE1E2.Cut$_{1+2}$.SPYΔN-His$_6$ (Kif/endo F-treated) | ns | 0.3475 |

Against ED43 HCVpp

| One-way ANOVA with Tukey's multiple comparisons test (w11) | Statistics | Adjusted P Value |
|---|---|---|
| HCV-1 sE1E2.Cut$_{1+2}$.SPYΔN-His$_6$ vs. HCV-1 sE1E2.Cut$_{1+2}$.SPYΔN-His$_6$ (Kif-treated) | ns | 0.3572 |
| HCV-1 sE1E2.Cut$_{1+2}$.SPYΔN-His$_6$ vs. HCV-1 sE1E2.Cut$_{1+2}$.SPYΔN-His$_6$ (Kif/endo F-treated) | ns | 0.5888 |
| HCV-1 sE1E2.Cut$_{1+2}$.SPYΔN-His$_6$ (Kif-treated) vs. HCV-1 sE1E2.Cut$_{1+2}$.SPYΔN-His$_6$ (Kif/endo F-treated) | ns | 0.0805 |

Against 5.2.1 HCVpp

| One-way ANOVA with Tukey's multiple comparisons test (w11) | Statistics | Adjusted P Value |
|---|---|---|
| HCV-1 sE1E2.Cut$_{1+2}$.SPYΔN-His$_6$ vs. HCV-1 sE1E2.Cut$_{1+2}$.SPYΔN-His$_6$ (Kif-treated) | ns | 0.5323 |
| HCV-1 sE1E2.Cut$_{1+2}$.SPYΔN-His$_6$ vs. HCV-1 sE1E2.Cut$_{1+2}$.SPYΔN-His$_6$ (Kif/endo F-treated) | ns | 0.9567 |
| HCV-1 sE1E2.Cut$_{1+2}$.SPYΔN-His$_6$ (Kif-treated) vs. HCV-1 sE1E2.Cut$_{1+2}$.SPYΔN-His$_6$ (Kif/endo F-treated) | ns | 0.5108 |

### Figure. S5

i Sera from mice immunized with glycan-modified HCV-1 sE1E2.Cut$_{1+2}$.SPYΔN-10GS-FR vaccine against H77 HCVpp

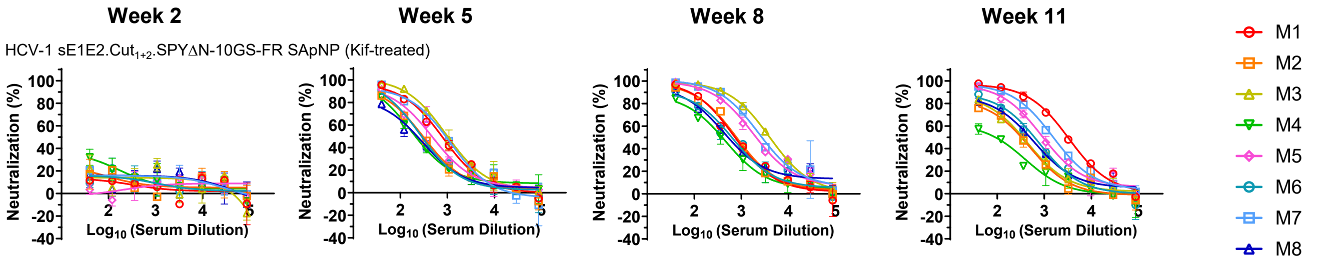


#### Mouse serum neutralizing ID$_{50}$ titers

Week 2

| Antigen | ID50 titers (week 2) | | | | | | | | Geometric Mean |
|---|---|---|---|---|---|---|---|---|---|
| | M1 | M2 | M3 | M4 | M5 | M6 | M7 | M8 | |
| HCV-1 sE1E2.Cut$_{1+2}$.SPYΔN-10GS-FR SApNP | <40 | 59.42 | <40 | <40 | <40 | <40 | <40 | <40 | N/A |
| HCV-1 sE1E2.Cut$_{1+2}$.SPYΔN-10GS-FR SApNP (Kif-treated) | <40 | <40 | <40 | <40 | <40 | <40 | <40 | <40 | N/A |

Week 5

| Antigen | ID50 titers (week 5) | | | | | | | | Geometric Mean |
|---|---|---|---|---|---|---|---|---|---|
| | M1 | M2 | M3 | M4 | M5 | M6 | M7 | M8 | |
| HCV-1 sE1E2.Cut$_{1+2}$.SPYΔN-10GS-FR SApNP | 942.9 | 1204 | 546.2 | 264.8 | 850.2 | 696.7 | 2697 | 755.6 | 816.8 |
| HCV-1 sE1E2.Cut$_{1+2}$.SPYΔN-10GS-FR SApNP (Kif-treated) | 739.1 | 303.7 | 1041 | 232.1 | 470.1 | 293.2 | 813.1 | 215.4 | 436.1 |

Week 8

| Antigen | ID50 titers (week 8) | | | | | | | | Geometric Mean |
|---|---|---|---|---|---|---|---|---|---|
| | M1 | M2 | M3 | M4 | M5 | M6 | M7 | M8 | |
| HCV-1 sE1E2.Cut$_{1+2}$.SPYΔN-10GS-FR SApNP | 1225 | 962.5 | 932.7 | 615.9 | 2262 | 1136 | 4831 | 723.5 | 1253.2 |
| HCV-1 sE1E2.Cut$_{1+2}$.SPYΔN-10GS-FR SApNP (Kif-treated) | 750.7 | 805.2 | 4050 | 373.4 | 2251 | 616.6 | 2943 | 595.5 | 1105.0 |

Week 11

| Antigen | ID50 titers (week 11) | | | | | | | | Geometric Mean |
|---|---|---|---|---|---|---|---|---|---|
| | M1 | M2 | M3 | M4 | M5 | M6 | M7 | M8 | |
| HCV-1 sE1E2.Cut$_{1+2}$.SPYΔN-10GS-FR SApNP | 1240 | 698.1 | 432.3 | 262.5 | 1058 | 3396 | 1098 | 422.1 | 797.5 |
| HCV-1 sE1E2.Cut$_{1+2}$.SPYΔN-10GS-FR SApNP (Kif-treated) | 3099 | 287.3 | 323.6 | 95.5 | 976.2 | 548.6 | 1572 | 469.7 | 568.3 |

#### Statistical analysis

| Unpaired t test (w5) | Statistics | P Value |
|---|---|---|
| HCV-1 sE1E2.Cut$_{1+2}$.SPYΔN-10GS-FR SApNP vs. HCV-1 sE1E2.Cut$_{1+2}$.SPYΔN-10GS-FR SApNP (Kif-treated) | ns | 0.1127 |

| Unpaired t test (w8) | Statistics | P Value |
|---|---|---|
| HCV-1 sE1E2.Cut$_{1+2}$.SPYΔN-10GS-FR SApNP vs. HCV-1 sE1E2.Cut$_{1+2}$.SPYΔN-10GS-FR SApNP (Kif-treated) | ns | 0.9571 |

| Unpaired t test (w11) | Statistics | P Value |
|---|---|---|
| HCV-1 sE1E2.Cut$_{1+2}$.SPYΔN-10GS-FR SApNP vs. HCV-1 sE1E2.Cut$_{1+2}$.SPYΔN-10GS-FR SApNP (Kif-treated) | ns | 0.7621 |

j Sera from mice immunized with glycan-modified HCV-1 sE1E2.Cut$_{1+2}$.SPYΔN-10GS-FR vaccine against heterologous HCVpps

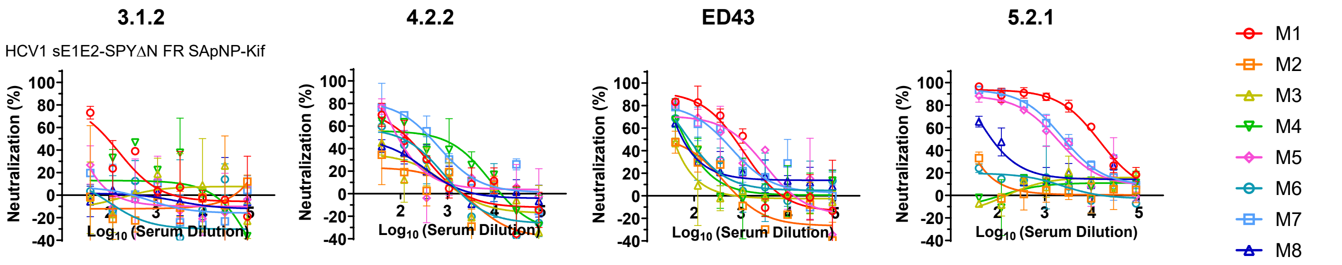


#### Mouse serum neutralizing ID$_{50}$ titers

Against 3.1.2 HCVpp

| Antigen | ID50 titers (week 11) | | | | | | | | Geometric Mean |
|---|---|---|---|---|---|---|---|---|---|
| | M1 | M2 | M3 | M4 | M5 | M6 | M7 | M8 | |
| HCV-1 sE1E2.Cut$_{1+2}$.SPYΔN-10GS-FR SApNP | 51.74 | <40 | <40 | <40 | <40 | 89.24 | <40 | <40 | N/A |
| HCV-1 sE1E2.Cut$_{1+2}$.SPYΔN-10GS-FR SApNP (Kif-treated) | 82.94 | <40 | <40 | <40 | <40 | <40 | <40 | <40 | N/A |

Against 4.2.2 HCVpp

| Antigen | ID50 titers (week 11) | | | | | | | | Geometric Mean |
|---|---|---|---|---|---|---|---|---|---|
| | M1 | M2 | M3 | M4 | M5 | M6 | M7 | M8 | |
| HCV-1 sE1E2.Cut$_{1+2}$.SPYΔN-10GS-FR SApNP | 55.29 | <40 | <40 | 44.08 | <40 | 257.8 | 71.24 | 61.22 | 52.1 |
| HCV-1 sE1E2.Cut$_{1+2}$.SPYΔN-10GS-FR SApNP (Kif-treated) | 111.3 | <40 | <40 | 263.6 | 95.38 | 87.61 | 372.8 | 45.79 | 86.2 |

Against ED43 HCVpp

| Antigen | ID50 titers (week 11) | | | | | | | | Geometric Mean |
|---|---|---|---|---|---|---|---|---|---|
| | M1 | M2 | M3 | M4 | M5 | M6 | M7 | M8 | |
| HCV-1 sE1E2.Cut$_{1+2}$.SPYΔN-10GS-FR SApNP | 213.5 | <40 | <40 | <40 | 57.86 | 1876 | 195.9 | <40 | 76.0 |
| HCV-1 sE1E2.Cut$_{1+2}$.SPYΔN-10GS-FR SApNP (Kif-treated) | 653.1 | 43.81 | <40 | 64.77 | 475.3 | 84.49 | 359.1 | 73.61 | 122.8 |

Against 5.2.1 HCVpp

| Antigen | ID50 titers (week 11) | | | | | | | | Geometric Mean |
|---|---|---|---|---|---|---|---|---|---|
| | M1 | M2 | M3 | M4 | M5 | M6 | M7 | M8 | |
| HCV-1 sE1E2.Cut$_{1+2}$.SPYΔN-10GS-FR SApNP | 2327 | 303.6 | 112.6 | 43.03 | 239.8 | 15202 | 553.6 | <40 | 285.7 |
| HCV-1 sE1E2.Cut$_{1+2}$.SPYΔN-10GS-FR SApNP (Kif-treated) | 12831 | 14.23 | <40 | <40 | 1760 | 16.04 | 2617 | 89.9 | 326.6 |

#### Statistical analysis

Against 4.2.2 HCVpp

| Unpaired t test (w11) | Statistics | P Value |
|---|---|---|
| HCV-1 sE1E2.Cut$_{1+2}$.SPYΔN-10GS-FR SApNP vs. HCV-1 sE1E2.Cut$_{1+2}$.SPYΔN-10GS-FR SApNP (Kif-treated) | ns | 0.2857 |

Against ED43 HCVpp

| Unpaired t test (w11) | Statistics | P Value |
|---|---|---|
| HCV-1 sE1E2.Cut$_{1+2}$.SPYΔN-10GS-FR SApNP vs. HCV-1 sE1E2.Cut$_{1+2}$.SPYΔN-10GS-FR SApNP (Kif-treated) | ns | 0.737 |

Against 5.2.1 HCVpp

| Unpaired t test (w11) | Statistics | P Value |
|---|---|---|
| HCV-1 sE1E2.Cut$_{1+2}$.SPYΔN-10GS-FR SApNP vs. HCV-1 sE1E2.Cut$_{1+2}$.SPYΔN-10GS-FR SApNP (Kif-treated) | ns | 0.8492 |

### Figure. S5

#### k Sera from mice immunized with glycan-modified HCV-1 sE1E2.Cut$_{1+2}$.SPYΔN-5GS-I3-10v9a-L7P vaccine against H77 HCVpp

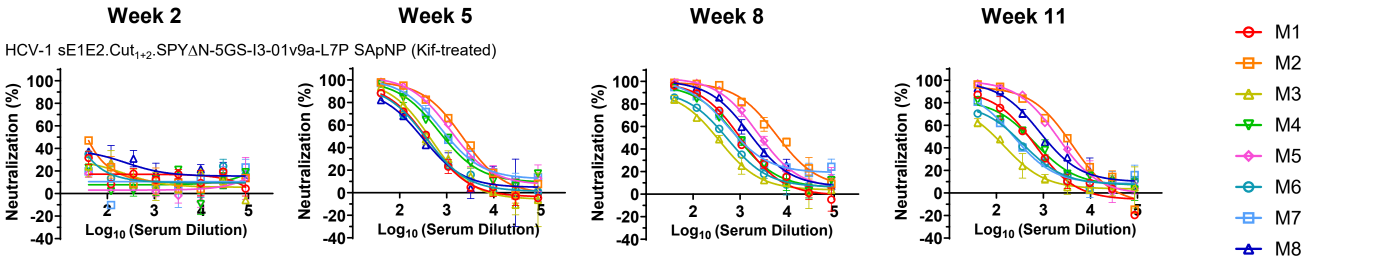


##### Mouse serum neutralizing ID$_{50}$ titers

**Week 2**

| Antigen | ID50 titers (week 2) | | | | | | | | Geometric Mean |
|---|---|---|---|---|---|---|---|---|---|
| | M1 | M2 | M3 | M4 | M5 | M6 | M7 | M8 | |
| HCV-1 sE1E2.Cut$_{1+2}$.SPYΔN-5GS-I3-01v9a-L7P SApNP | <40 | <40 | <40 | <40 | <40 | <40 | <40 | <40 | N/A |
| HCV-1 sE1E2.Cut$_{1+2}$.SPYΔN-5GS-I3-01v9a-L7P SApNP (Kif-treated) | <40 | <40 | <40 | <40 | <40 | <40 | <40 | <40 | N/A |

**Week 5**

| Antigen | ID50 titers (week 5) | | | | | | | | Geometric Mean |
|---|---|---|---|---|---|---|---|---|---|
| | M1 | M2 | M3 | M4 | M5 | M6 | M7 | M8 | |
| HCV-1 sE1E2.Cut$_{1+2}$.SPYΔN-5GS-I3-01v9a-L7P SApNP | 339 | 284.5 | 519.5 | 503.5 | 2739 | 1054 | 753.6 | 737.9 | 669.8 |
| HCV-1 sE1E2.Cut$_{1+2}$.SPYΔN-5GS-I3-01v9a-L7P SApNP (Kif-treated) | 331.5 | 2080 | 393.2 | 916.7 | 1633 | 336.1 | 1203 | 286.3 | 682.3 |

**Week 8**

| Antigen | ID50 titers (week 8) | | | | | | | | Geometric Mean |
|---|---|---|---|---|---|---|---|---|---|
| | M1 | M2 | M3 | M4 | M5 | M6 | M7 | M8 | |
| HCV-1 sE1E2.Cut$_{1+2}$.SPYΔN-5GS-I3-01v9a-L7P SApNP | 829.9 | 749.2 | 607.8 | 1779 | 2080 | 2187 | 690.5 | 2843 | 1251.1 |
| HCV-1 sE1E2.Cut$_{1+2}$.SPYΔN-5GS-I3-01v9a-L7P SApNP (Kif-treated) | 998.8 | 6919 | 317.1 | 921.7 | 3002 | 517.5 | 1108 | 2008 | 1274.9 |

**Week 11**

| Antigen | ID50 titers (week 11) | | | | | | | | Geometric Mean |
|---|---|---|---|---|---|---|---|---|---|
| | M1 | M2 | M3 | M4 | M5 | M6 | M7 | M8 | |
| HCV-1 sE1E2.Cut$_{1+2}$.SPYΔN-5GS-I3-01v9a-L7P SApNP | 423.6 | 141.9 | 163.5 | 3087 | 586.8 | 575 | 518 | 3912 | 616.0 |
| HCV-1 sE1E2.Cut$_{1+2}$.SPYΔN-5GS-I3-01v9a-L7P SApNP (Kif-treated) | 422.9 | 2577 | 96.14 | 445.4 | 1944 | 240.6 | 246 | 1110 | 527.1 |

##### Statistical analysis

| Unpaired t test (w5) | Statistics | P Value |
|---|---|---|
| HCV-1 sE1E2.Cut$_{1+2}$.SPYΔN-5GS-I3-01v9a-L7P SApNP vs. HCV-1 sE1E2.Cut$_{1+2}$.SPYΔN-5GS-I3-01v9a-L7P SApNP (Kif-treated) | ns | 0.9345 |

| Unpaired t test (w8) | Statistics | P Value |
|---|---|---|
| HCV-1 sE1E2.Cut$_{1+2}$.SPYΔN-5GS-I3-01v9a-L7P SApNP vs. HCV-1 sE1E2.Cut$_{1+2}$.SPYΔN-5GS-I3-01v9a-L7P SApNP (Kif-treated) | ns | 0.5528 |

| Unpaired t test (w11) | Statistics | P Value |
|---|---|---|
| HCV-1 sE1E2.Cut$_{1+2}$.SPYΔN-5GS-I3-01v9a-L7P SApNP vs. HCV-1 sE1E2.Cut$_{1+2}$.SPYΔN-5GS-I3-01v9a-L7P SApNP (Kif-treated) | ns | 0.6409 |

#### l Sera from mice immunized with glycan-modified HCV-1 sE1E2.Cut$_{1+2}$.SPYΔN-5GS-I3-10v9a-L7P vaccine against heterologous HCVpps

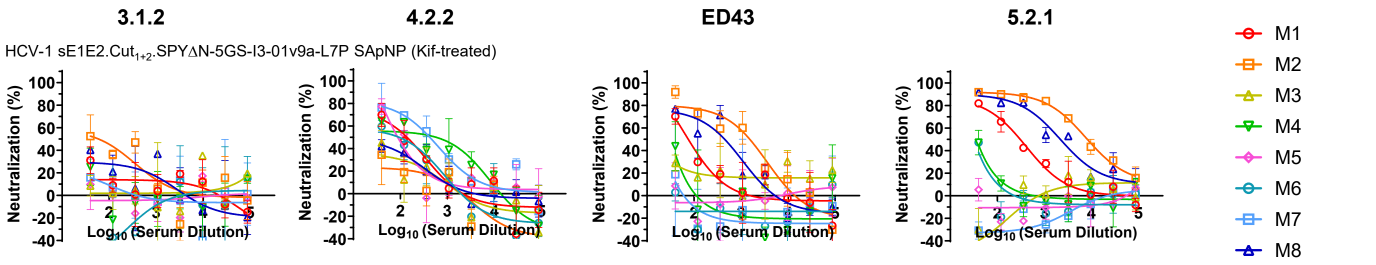


##### Mouse serum neutralizing ID$_{50}$ titers

**Against 3.1.2 HCVpp**

| Antigen | ID50 titers (week 11) | | | | | | | | Geometric Mean |
|---|---|---|---|---|---|---|---|---|---|
| | M1 | M2 | M3 | M4 | M5 | M6 | M7 | M8 | |
| HCV-1 sE1E2.Cut$_{1+2}$.SPYΔN-5GS-I3-01v9a-L7P SApNP | <40 | <40 | <40 | <40 | <40 | <40 | <40 | 65.6 | N/A |
| HCV-1 sE1E2.Cut$_{1+2}$.SPYΔN-5GS-I3-01v9a-L7P SApNP (Kif-treated) | <40 | 77.78 | <40 | <40 | <40 | <40 | <40 | <40 | N/A |

**Against 4.2.2 HCVpp**

| Antigen | ID50 titers (week 11) | | | | | | | | Geometric Mean |
|---|---|---|---|---|---|---|---|---|---|
| | M1 | M2 | M3 | M4 | M5 | M6 | M7 | M8 | |
| HCV-1 sE1E2.Cut$_{1+2}$.SPYΔN-5GS-I3-01v9a-L7P SApNP | 88.57 | <40 | <40 | 339.9 | 128.6 | 124 | <40 | 549.5 | 69.9 |
| HCV-1 sE1E2.Cut$_{1+2}$.SPYΔN-5GS-I3-01v9a-L7P SApNP (Kif-treated) | 51.04 | 113.4 | 61.11 | <40 | <40 | 42.88 | <40 | 310 | 53.6 |

**Against ED43 HCVpp**

| Antigen | ID50 titers (week 11) | | | | | | | | Geometric Mean |
|---|---|---|---|---|---|---|---|---|---|
| | M1 | M2 | M3 | M4 | M5 | M6 | M7 | M8 | |
| HCV-1 sE1E2.Cut$_{1+2}$.SPYΔN-5GS-I3-01v9a-L7P SApNP | 469.6 | 103.4 | <40 | 1225 | 383 | 51.91 | 124 | 2064 | 145.0 |
| HCV-1 sE1E2.Cut$_{1+2}$.SPYΔN-5GS-I3-01v9a-L7P SApNP (Kif-treated) | 71.9 | 908.1 | <40 | <40 | <40 | <40 | <40 | 331.8 | 17.7 |

**Against 5.2.1 HCVpp**

| Antigen | ID50 titers (week 11) | | | | | | | | Geometric Mean |
|---|---|---|---|---|---|---|---|---|---|
| | M1 | M2 | M3 | M4 | M5 | M6 | M7 | M8 | |
| HCV-1 sE1E2.Cut$_{1+2}$.SPYΔN-5GS-I3-01v9a-L7P SApNP | 689.2 | 308.3 | <40 | 8928 | 1576 | 229.9 | 190.4 | 9417 | 1030.3 |
| HCV-1 sE1E2.Cut$_{1+2}$.SPYΔN-5GS-I3-01v9a-L7P SApNP (Kif-treated) | 275.7 | 8223 | <40 | <40 | <40 | <40 | <40 | 2196 | 289.9 |

##### Statistical analysis

**Against 4.2.2 HCVpp**

| Unpaired t test (w11) | Statistics | P Value |
|---|---|---|
| HCV-1 sE1E2.Cut$_{1+2}$.SPYΔN-5GS-I3-01v9a-L7P SApNP vs. HCV-1 sE1E2.Cut$_{1+2}$.SPYΔN-5GS-I3-01v9a-L7P SApNP (Kif-treated) | ns | 0.3082 |

**Against ED43 HCVpp**

| Unpaired t test (w11) | Statistics | P Value |
|---|---|---|
| HCV-1 sE1E2.Cut$_{1+2}$.SPYΔN-5GS-I3-01v9a-L7P SApNP vs. HCV-1 sE1E2.Cut$_{1+2}$.SPYΔN-5GS-I3-01v9a-L7P SApNP (Kif-treated) | ns | 0.2525 |

**Against 5.2.1 HCVpp**

| Unpaired t test (w11) | Statistics | P Value |
|---|---|---|
| HCV-1 sE1E2.Cut$_{1+2}$.SPYΔN-5GS-I3-01v9a-L7P SApNP vs. HCV-1 sE1E2.Cut$_{1+2}$.SPYΔN-5GS-I3-01v9a-L7P SApNP (Kif-treated) | ns | 0.7049 |

**Figure. S5**

**Fig. S5. Immunogenicity of rationally designed HCV-1 sE1E2.Cut$_{1+2}$.SPYΔN vaccines in mice.** (**a**) Neutralization curves of mouse sera from HCV-1 sE1E2.Cut$_{1+2}$.SPYΔN dimer and SApNP vaccine groups (n = 8 mice/group) against H77 HCVpp. The administered dose was 200 µl of antigen/AH + CpG adjuvant mix containing 10 µg of immunogen and 100 µl of adjuvant. Mice were immunized at weeks 0, 3, 6 and 9 with 3-week intervals via the intraperitoneal (i.p.) route. (**b**) (Top) Summary of geometric mean $ID_{50}$ titers measured for HCV-1 sE1E2.Cut$_{1+2}$.SPYΔN vaccine groups against H77 HCVpp. Color coding indicates the level of $ID_{50}$ titers (white: no neutralization; green to red: low to high neutralization). Of note, the $ID_{50}$ values were derived by setting the minimum/maximum % neutralization to 0.0/100.0 as constraints. (Bottom) Summary of statistical analysis performed for each timepoint. (**c**) Neutralization curves of mouse sera from HCV-1 sE1E2.Cut$_{1+2}$.SPYΔN vaccine groups at week 11 against heterologous 3.1.2, 4.2.2, ED43, and 5.2.1 HCVpps. (**d**) Summary of geometric mean $ID_{50}$ values and statistical analysis. (**e**) Neutralization curves of mouse sera from glycan-modified HCV-1 sE1E2.Cut$_{1+2}$.SPYΔN dimer vaccine groups against H77 HCVpp. (**f**) Summary of geometric mean $ID_{50}$ values and statistical analysis. (**g**) Neutralization curves of mouse sera from glycan-modified HCV-1 sE1E2.Cut$_{1+2}$.SPYΔN dimer vaccine groups at week 11 against heterologous 3.1.2, 4.2.2, ED43, and 5.2.1 HCVpps. (**h**) Summary of geometric mean $ID_{50}$ values and statistical analysis. (**i**) Neutralization curves of mouse sera from glycan-modified HCV-1 sE1E2.Cut$_{1+2}$.SPYΔN-10GS-FR SApNP vaccine groups against H77 HCVpp. Summary of geometric mean $ID_{50}$ values and statistical analysis. (**j**) Neutralization curves of mouse sera from glycan-modified HCV1 sE1E2.Cut$_{1+2}$.SPYΔN-10GS-FR SApNP vaccine groups at week 11 against heterologous 3.1.2, 4.2.2, ED43, and 5.2.1 HCVpps. Summary of geometric mean $ID_{50}$ values and statistical analysis. (**k**) Neutralization curves of mouse sera from glycan-modified HCV-1 sE1E2.Cut$_{1+2}$.SPYΔN-5GS-I3-01v9a-L7P SApNP vaccine groups against H77 HCVpp. Summary of geometric mean $ID_{50}$ values and statistical analysis. (**l**) Neutralization curves of mouse sera from glycan-modified HCV-1 sE1E2.Cut$_{1+2}$.SPYΔN-5GS-I3-01v9a-L7P SApNP vaccine groups at week 11 against heterologous 3.1.2, 4.2.2, ED43, and 5.2.1 HCVpps. Summary of geometric mean $ID_{50}$ values and statistical analysis. Error bars represent the difference between duplicate values at each concentration tested for each sample. $ID_{50}$ values were calculated in GraphPad Prism 10.3.1. Data were analyzed using one-way ANOVA, followed by Tukey's multiple comparison post hoc test for each timepoint. Two-tailed unpaired t-tests were used to compare the geometric means of two groups. Statistical significance is indicated as follows: ns (not significant) and *$p < 0.05$.

### Figure. S6

**a** **Sera from mice immunized with glycan-modified HCV-1 sE1E2.Cut$_{1+2}$.SPYΔN-His$_6$ (HEK293F) vaccines against H77 HCVpp**

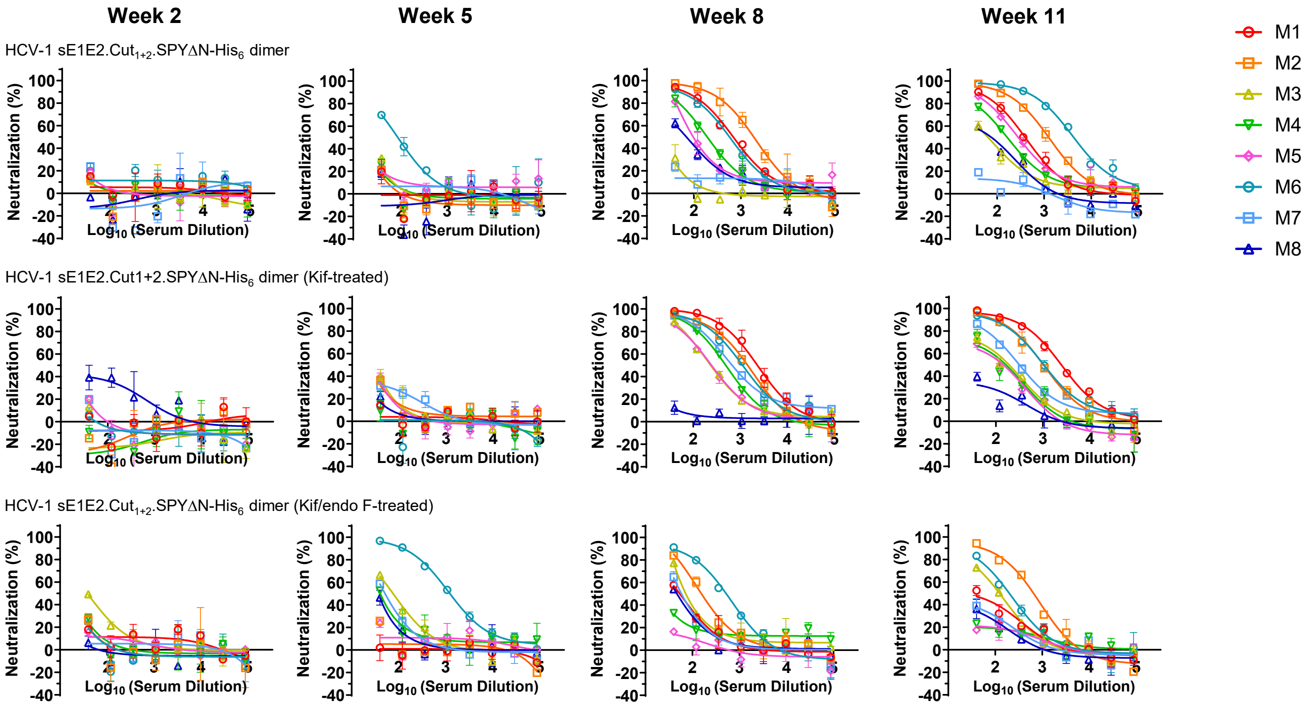


**b** **Mouse serum neutralizing ID$_{50}$ titers**

**Week 2**

| Antigen | ID50 titers (week 2) M1 | M2 | M3 | M4 | M5 | M6 | M7 | M8 | Geometric Mean |
|---|---|---|---|---|---|---|---|---|---|
| HCV-1 sE1E2.Cut$_{1+2}$.SPYΔN-His$_6$ | <40 | <40 | <40 | <40 | <40 | <40 | <40 | <40 | N/A |
| HCV-1 sE1E2.Cut$_{1+2}$.SPYΔN-His$_6$ (Kif-treated) | <40 | <40 | <40 | <40 | <40 | <40 | <40 | 46.28 | N/A |
| HCV-1 sE1E2.Cut$_{1+2}$.SPYΔN-His$_6$ (Kif/endo F-treated) | <40 | <40 | <40 | <40 | <40 | <40 | <40 | <40 | N/A |

**Week 5**

| Antigen | ID50 titers (week 5) M1 | M2 | M3 | M4 | M5 | M6 | M7 | M8 | Geometric Mean |
|---|---|---|---|---|---|---|---|---|---|
| HCV-1 sE1E2.Cut$_{1+2}$.SPYΔN-His$_6$ | <40 | <40 | <40 | <40 | <40 | 86.06 | <40 | <40 | 7.5 |
| HCV-1 sE1E2.Cut$_{1+2}$.SPYΔN-His$_6$ (Kif-treated) | <40 | <40 | <40 | <40 | <40 | <40 | <40 | <40 | 8.5 |
| HCV-1 sE1E2.Cut$_{1+2}$.SPYΔN-His$_6$ (Kif/endo F-treated) | <40 | <40 | 74.51 | <40 | <40 | 1280 | 48.09 | <40 | 22.4 |

**Week 8**

| Antigen | ID50 titers (week 8) M1 | M2 | M3 | M4 | M5 | M6 | M7 | M8 | Geometric Mean |
|---|---|---|---|---|---|---|---|---|---|
| HCV-1 sE1E2.Cut$_{1+2}$.SPYΔN-His$_6$ | 687.5 | 1746 | <40 | 195 | 120.3 | 555.7 | <40 | 74.86 | 145.0 |
| HCV-1 sE1E2.Cut$_{1+2}$.SPYΔN-His$_6$ (Kif-treated) | 2276 | 1262 | 252.3 | 478.1 | 249.9 | 1099 | 768.1 | <40 | 376.0 |
| HCV-1 sE1E2.Cut$_{1+2}$.SPYΔN-His$_6$ (Kif/endo F-treated) | 52.99 | 163.1 | 87.56 | <40 | <40 | 459.6 | 58.6 | 45.87 | 59.7 |

**Week 11**

| Antigen | ID50 titers (week 11) M1 | M2 | M3 | M4 | M5 | M6 | M7 | M8 | Geometric Mean |
|---|---|---|---|---|---|---|---|---|---|
| HCV-1 sE1E2.Cut$_{1+2}$.SPYΔN-His$_6$ | 385.2 | 1219 | 69.76 | 169 | 297.4 | 4365 | <40 | 72.91 | 215.4 |
| HCV-1 sE1E2.Cut$_{1+2}$.SPYΔN-His$_6$ (Kif-treated) | 2438 | 999.3 | 163.3 | 129.5 | 113.5 | 1075 | 331.3 | <40 | 295.2 |
| HCV-1 sE1E2.Cut$_{1+2}$.SPYΔN-His$_6$ (Kif/endo F-treated) | 56.68 | 556.6 | 115.1 | <40 | <40 | 187.2 | <40 | <40 | 57.5 |

**Statistical analysis**

| One-way ANOVA with Tukey's multiple comparisons test (w5) | Statistics | Adjusted P Value |
|---|---|---|
| HCV-1 sE1E2.Cut$_{1+2}$.SPYΔN-His$_6$ vs. HCV-1 sE1E2.Cut$_{1+2}$.SPYΔN-His$_6$ (Kif-treated) | ns | 0.9995 |
| HCV-1 sE1E2.Cut$_{1+2}$.SPYΔN-His$_6$ vs. HCV-1 sE1E2.Cut$_{1+2}$.SPYΔN-His$_6$ (Kif/endo F-treated) | ns | 0.3964 |
| HCV-1 sE1E2.Cut$_{1+2}$.SPYΔN-His$_6$ (Kif-treated) vs. HCV-1 sE1E2.Cut$_{1+2}$.SPYΔN-His$_6$ (Kif/endo F-treated) | ns | 0.3805 |

| One-way ANOVA with Tukey's multiple comparisons test (w8) | Statistics | Adjusted P Value |
|---|---|---|
| HCV-1 sE1E2.Cut$_{1+2}$.SPYΔN-His$_6$ vs. HCV-1 sE1E2.Cut$_{1+2}$.SPYΔN-His$_6$ (Kif-treated) | ns | 0.384 |
| HCV-1 sE1E2.Cut$_{1+2}$.SPYΔN-His$_6$ vs. HCV-1 sE1E2.Cut$_{1+2}$.SPYΔN-His$_6$ (Kif/endo F-treated) | ns | 0.5044 |
| HCV-1 sE1E2.Cut$_{1+2}$.SPYΔN-His$_6$ (Kif-treated) vs. HCV-1 sE1E2.Cut$_{1+2}$.SPYΔN-His$_6$ (Kif/endo F-treated) | ns | 0.0538 |

| One-way ANOVA with Tukey's multiple comparisons test (w11) | Statistics | Adjusted P Value |
|---|---|---|
| HCV-1 sE1E2.Cut$_{1+2}$.SPYΔN-His$_6$ vs. HCV-1 sE1E2.Cut$_{1+2}$.SPYΔN-His$_6$ (Kif-treated) | ns | 0.9412 |
| HCV-1 sE1E2.Cut$_{1+2}$.SPYΔN-His$_6$ vs. HCV-1 sE1E2.Cut$_{1+2}$.SPYΔN-His$_6$ (Kif/endo F-treated) | ns | 0.3509 |
| HCV-1 sE1E2.Cut$_{1+2}$.SPYΔN-His$_6$ (Kif-treated) vs. HCV-1 sE1E2.Cut$_{1+2}$.SPYΔN-His$_6$ (Kif/endo F-treated) | ns | 0.534 |

### Figure. S6

#### C Week-11 sera from mice immunized with HCV-1 sE1E2.Cut$_{1+2}$.SPYΔN dimer (HEK293F) vaccines against heterologous HCVpps

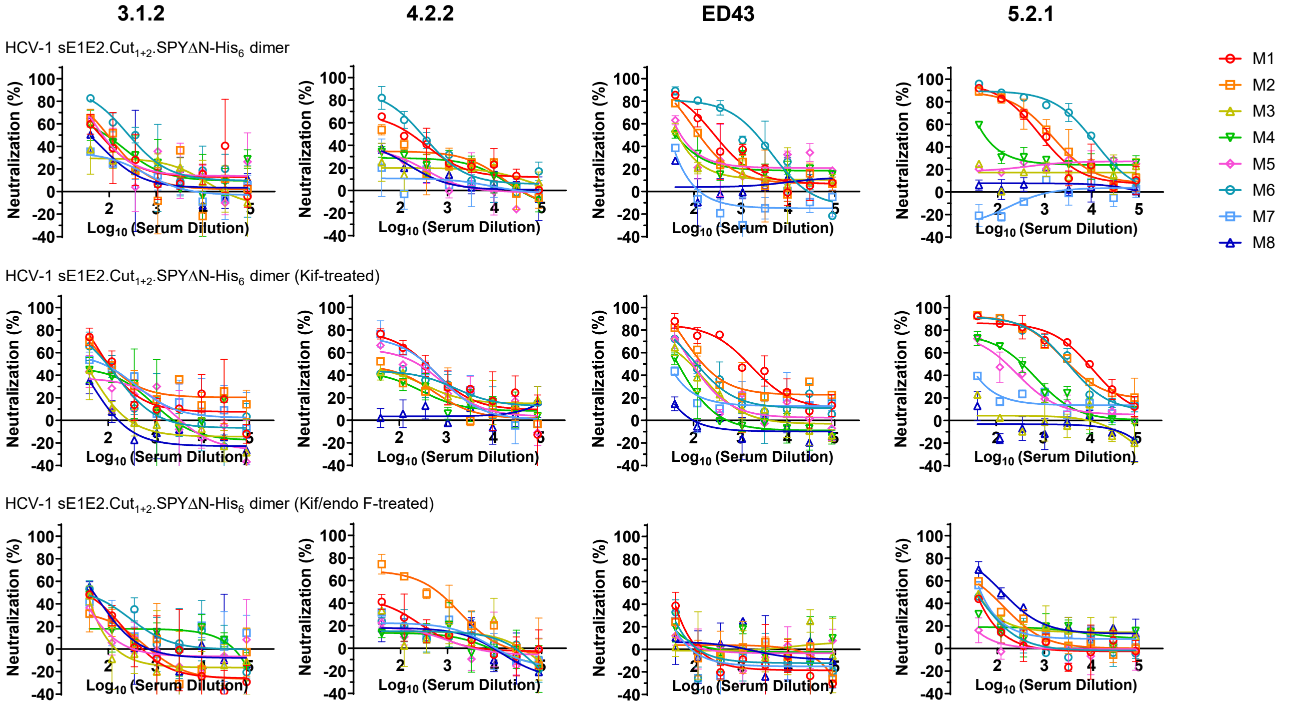


#### d Mouse serum neutralizing ID$_{50}$ titers

Against 3.1.2 HCVpp

| Antigen | ID50 titers (week 11) | | | | | | | | Geometric Mean |
|---|---|---|---|---|---|---|---|---|---|
| | M1 | M2 | M3 | M4 | M5 | M6 | M7 | M8 | |
| HCV-1 sE1E2.Cut$_{1+2}$.SPYΔN-His$_6$ | 79.25 | 94.84 | <40 | 99.23 | 88.83 | 279.2 | <40 | 45.45 | 74.4 |
| HCV-1 sE1E2.Cut$_{1+2}$.SPYΔN-His$_6$ (Kif-treated) | 110.4 | 141.9 | <40 | 57.88 | 44.06 | 85.92 | 94.01 | <40 | 57.7 |
| HCV-1 sE1E2.Cut$_{1+2}$.SPYΔN-His$_6$ (Kif/endo F-treated) | <40 | <40 | <40 | <40 | <40 | 56.43 | <40 | 43.95 | 30.3 |

Against 4.2.2 HCVpp

| Antigen | ID50 titers (week 11) | | | | | | | | Geometric Mean |
|---|---|---|---|---|---|---|---|---|---|
| | M1 | M2 | M3 | M4 | M5 | M6 | M7 | M8 | |
| HCV-1 sE1E2.Cut$_{1+2}$.SPYΔN-His$_6$ | 161.8 | 69.29 | <40 | <40 | <40 | 278.1 | <40 | <40 | 44.3 |
| HCV-1 sE1E2.Cut$_{1+2}$.SPYΔN-His$_6$ (Kif-treated) | 339.9 | 64.19 | 67.7 | 44.19 | 191.6 | 85.77 | 278.9 | <40 | 71.0 |
| HCV-1 sE1E2.Cut$_{1+2}$.SPYΔN-His$_6$ (Kif/endo F-treated) | <40 | 366.6 | <40 | <40 | <40 | <40 | <40 | <40 | 24.3 |

Against ED43 HCVpp

| Antigen | ID50 titers (week 11) | | | | | | | | Geometric Mean |
|---|---|---|---|---|---|---|---|---|---|
| | M1 | M2 | M3 | M4 | M5 | M6 | M7 | M8 | |
| HCV-1 sE1E2.Cut$_{1+2}$.SPYΔN-His$_6$ | 287 | 143.3 | 51.67 | 58 | 63.12 | 1277 | <40 | <40 | 76.9 |
| HCV-1 sE1E2.Cut$_{1+2}$.SPYΔN-His$_6$ (Kif-treated) | 1418 | 216.1 | 81.62 | <40 | 99.3 | 132.1 | <40 | <40 | 81.1 |
| HCV-1 sE1E2.Cut$_{1+2}$.SPYΔN-His$_6$ (Kif/endo F-treated) | <40 | <40 | <40 | <40 | <40 | <40 | <40 | <40 | 5.2 |

Against 5.2.1 HCVpp

| Antigen | ID50 titers (week 11) | | | | | | | | Geometric Mean |
|---|---|---|---|---|---|---|---|---|---|
| | M1 | M2 | M3 | M4 | M5 | M6 | M7 | M8 | |
| HCV-1 sE1E2.Cut$_{1+2}$.SPYΔN-His$_6$ | 854.5 | 1338 | <40 | 81.83 | <40 | 7783 | <40 | <40 | 127.5 |
| HCV-1 sE1E2.Cut$_{1+2}$.SPYΔN-His$_6$ (Kif-treated) | 7752 | 4044 | <40 | 296.3 | 158.9 | 3366 | <40 | <40 | 204.7 |
| HCV-1 sE1E2.Cut$_{1+2}$.SPYΔN-His$_6$ (Kif/endo F-treated) | <40 | 68.19 | 43.7 | <40 | <40 | <40 | 45.54 | 145.8 | 35.0 |

#### Statistical analysis

Against 3.1.2 HCVpp

| One-way ANOVA with Tukey's multiple comparisons test (w11) | Statistics | Adjusted P Value |
|---|---|---|
| HCV-1 sE1E2.Cut$_{1+2}$.SPYΔN-His$_6$ vs. HCV-1 sE1E2.Cut$_{1+2}$.SPYΔN-His$_6$ (Kif-treated) | ns | 0.6781 |
| HCV-1 sE1E2.Cut$_{1+2}$.SPYΔN-His$_6$ vs. HCV-1 sE1E2.Cut$_{1+2}$.SPYΔN-His$_6$ (Kif/endo F-treated) | ns | 0.0735 |
| HCV-1 sE1E2.Cut$_{1+2}$.SPYΔN-His$_6$ (Kif-treated) vs. HCV-1 sE1E2.Cut$_{1+2}$.SPYΔN-His$_6$ (Kif/endo F-treated) | ns | 0.3196 |

Against 4.2.2 HCVpp

| One-way ANOVA with Tukey's multiple comparisons test (w11) | Statistics | Adjusted P Value |
|---|---|---|
| HCV-1 sE1E2.Cut$_{1+2}$.SPYΔN-His$_6$ vs. HCV-1 sE1E2.Cut$_{1+2}$.SPYΔN-His$_6$ (Kif-treated) | ns | 0.5995 |
| HCV-1 sE1E2.Cut$_{1+2}$.SPYΔN-His$_6$ vs. HCV-1 sE1E2.Cut$_{1+2}$.SPYΔN-His$_6$ (Kif/endo F-treated) | ns | 0.9521 |
| HCV-1 sE1E2.Cut$_{1+2}$.SPYΔN-His$_6$ (Kif-treated) vs. HCV-1 sE1E2.Cut$_{1+2}$.SPYΔN-His$_6$ (Kif/endo F-treated) | ns | 0.4244 |

Against ED43 HCVpp

| One-way ANOVA with Tukey's multiple comparisons test (w11) | Statistics | Adjusted P Value |
|---|---|---|
| HCV-1 sE1E2.Cut$_{1+2}$.SPYΔN-His$_6$ vs. HCV-1 sE1E2.Cut$_{1+2}$.SPYΔN-His$_6$ (Kif-treated) | ns | 0.9964 |
| HCV-1 sE1E2.Cut$_{1+2}$.SPYΔN-His$_6$ vs. HCV-1 sE1E2.Cut$_{1+2}$.SPYΔN-His$_6$ (Kif/endo F-treated) | ns | 0.438 |
| HCV-1 sE1E2.Cut$_{1+2}$.SPYΔN-His$_6$ (Kif-treated) vs. HCV-1 sE1E2.Cut$_{1+2}$.SPYΔN-His$_6$ (Kif/endo F-treated) | ns | 0.3942 |

Against 5.2.1 HCVpp

| One-way ANOVA with Tukey's multiple comparisons test (w11) | Statistics | Adjusted P Value |
|---|---|---|
| HCV-1 sE1E2.Cut$_{1+2}$.SPYΔN-His$_6$ vs. HCV-1 sE1E2.Cut$_{1+2}$.SPYΔN-His$_6$ (Kif-treated) | ns | 0.9021 |
| HCV-1 sE1E2.Cut$_{1+2}$.SPYΔN-His$_6$ vs. HCV-1 sE1E2.Cut$_{1+2}$.SPYΔN-His$_6$ (Kif/endo F-treated) | ns | 0.484 |
| HCV-1 sE1E2.Cut$_{1+2}$.SPYΔN-His$_6$ (Kif-treated) vs. HCV-1 sE1E2.Cut$_{1+2}$.SPYΔN-His$_6$ (Kif/endo F-treated) | ns | 0.2449 |

**e** **Sera from mice immunized with glycan-modified HCV-1 sE1E2.Cut$_{1+2}$.SPYΔN-His$_6$ dimer vaccines against H77 HCVpp**

**Figure. S6**

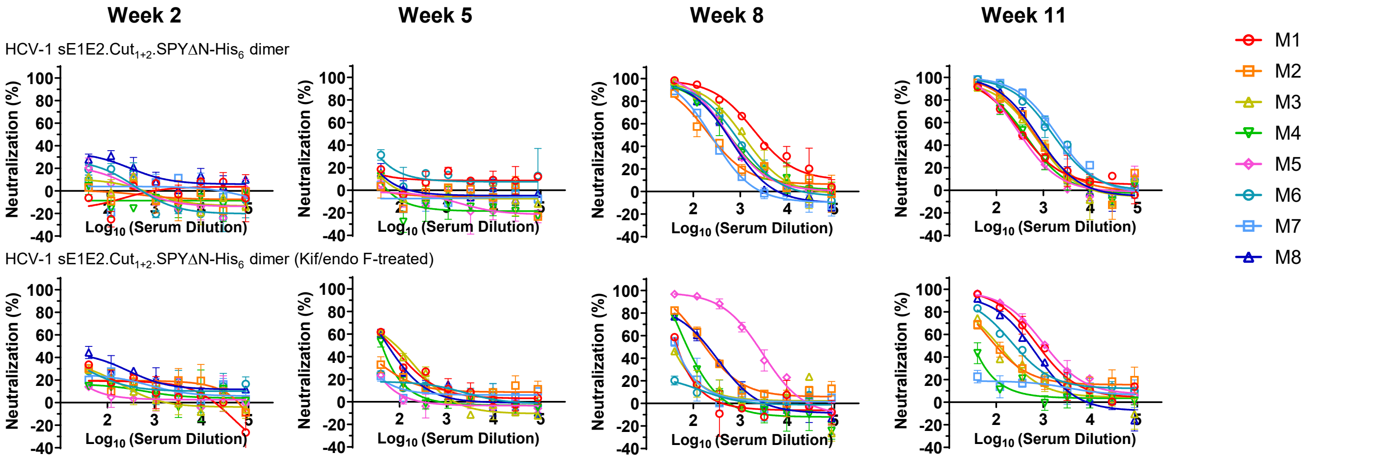


#### Mouse serum neutralizing ID$_{50}$ titers

**Week 2**

| Antigen | ID50 titers (week 2) M1 | M2 | M3 | M4 | M5 | M6 | M7 | M8 | Geometric Mean |
|---|---|---|---|---|---|---|---|---|---|
| HCV-1 sE1E2.Cut$_{1+2}$.SPYΔN-His$_6$ | <40 | <40 | <40 | <40 | <40 | <40 | <40 | <40 | N/A |
| HCV-1 sE1E2.Cut$_{1+2}$.SPYΔN-His$_6$ (Kif/endo F-treated) | <40 | <40 | <40 | <40 | <40 | <40 | <40 | 145.8 | N/A |

**Week 5**

| Antigen | ID50 titers (week 5) M1 | M2 | M3 | M4 | M5 | M6 | M7 | M8 | Geometric Mean |
|---|---|---|---|---|---|---|---|---|---|
| HCV-1 sE1E2.Cut$_{1+2}$.SPYΔN-His$_6$ | <40 | <40 | <40 | <40 | <40 | <40 | <40 | <40 | 2.8 |
| HCV-1 sE1E2.Cut$_{1+2}$.SPYΔN-His$_6$ (Kif/endo F-treated) | 69.62 | <40 | 75.43 | <40 | <40 | <40 | <40 | 54.73 | 28.4 |

**Week 8**

| Antigen | ID50 titers (week 8) M1 | M2 | M3 | M4 | M5 | M6 | M7 | M8 | Geometric Mean |
|---|---|---|---|---|---|---|---|---|---|
| HCV-1 sE1E2.Cut$_{1+2}$.SPYΔN-His$_6$ | 2623 | 241 | 1060 | 570.2 | 589.3 | 719.7 | 221.6 | 483.6 | 602.5 |
| HCV-1 sE1E2.Cut$_{1+2}$.SPYΔN-His$_6$ (Kif/endo F-treated) | <40 | 194.2 | <40 | 68.05 | 2151 | <40 | <40 | 176.4 | 84.1 |

**Week 11**

| Antigen | ID50 titers (week 11) M1 | M2 | M3 | M4 | M5 | M6 | M7 | M8 | Geometric Mean |
|---|---|---|---|---|---|---|---|---|---|
| HCV-1 sE1E2.Cut$_{1+2}$.SPYΔN-His$_6$ | 385.8 | 636.9 | 558.7 | 377.3 | 321.9 | 1548 | 1849 | 685.6 | 652.2 |
| HCV-1 sE1E2.Cut$_{1+2}$.SPYΔN-His$_6$ (Kif/endo F-treated) | 808.9 | 121.7 | 118 | <40 | 1176 | 278.8 | <40 | 473 | 172.9 |

#### Statistical analysis

| Unpaired t test (w8) | Statistics | P Value |
|---|---|---|
| HCV-1 sE1E2.Cut$_{1+2}$.SPYΔN-His$_6$ vs. HCV-1 sE1E2.Cut$_{1+2}$.SPYΔN-His$_6$ (Kif/endo F-treated) | ns | 0.2301 |

| Unpaired t test (w11) | Statistics | P Value |
|---|---|---|
| HCV-1 sE1E2.Cut$_{1+2}$.SPYΔN-His$_6$ vs. HCV-1 sE1E2.Cut$_{1+2}$.SPYΔN-His$_6$ (Kif/endo F-treated) | ns | 0.1199 |

**f** **Sera from mice immunized with glycan-modified HCV-1 sE1E2.Cut$_{1+2}$.SPYΔN-His$_6$ dimer vaccines against heterologous HCVpps**

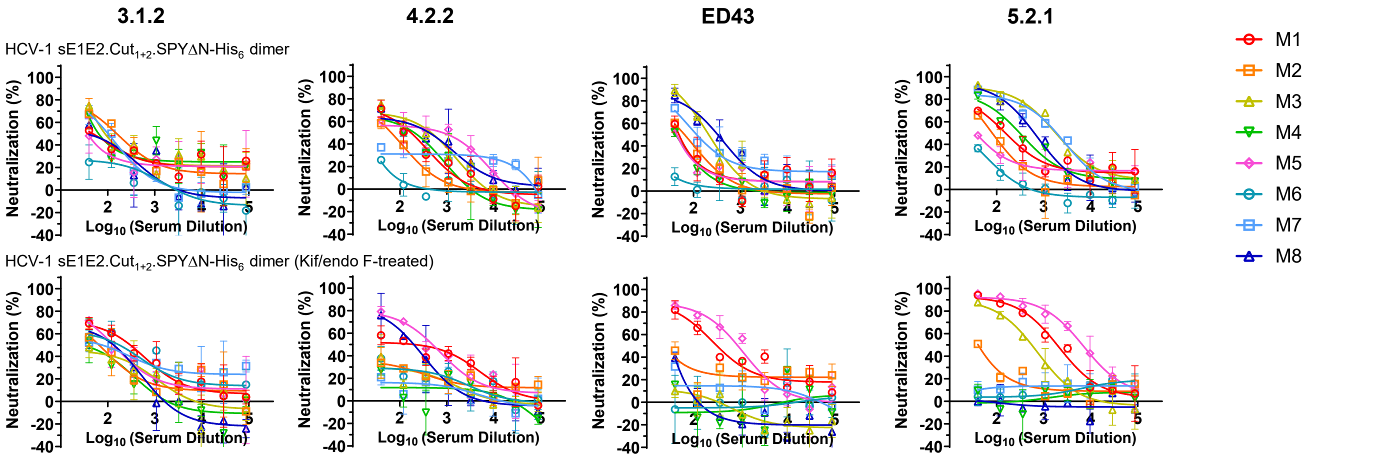


#### Mouse serum neutralizing ID$_{50}$ titers

**Against 3.1.2 HCVpp**

| Antigen | ID50 titers (week 11) M1 | M2 | M3 | M4 | M5 | M6 | M7 | M8 | Geometric Mean |
|---|---|---|---|---|---|---|---|---|---|
| HCV-1 sE1E2.Cut$_{1+2}$.SPYΔN-His$_6$ | 81.3 | 174.2 | 140.9 | 103.3 | 53.59 | <40 | 97.35 | 62.33 | 78.5 |
| HCV-1 sE1E2.Cut$_{1+2}$.SPYΔN-His$_6$ (Kif/endo F-treated) | 254.3 | 76.63 | 64.29 | 54.18 | 145 | 176.3 | 132.1 | 109.8 | 112.2 |

**Against 4.2.2 HCVpp**

| Antigen | ID50 titers (week 11) M1 | M2 | M3 | M4 | M5 | M6 | M7 | M8 | Geometric Mean |
|---|---|---|---|---|---|---|---|---|---|
| HCV-1 sE1E2.Cut$_{1+2}$.SPYΔN-His$_6$ | 139.5 | 65.31 | 270.1 | 138.3 | 409.8 | <40 | 56.57 | 272 | 111.9 |
| HCV-1 sE1E2.Cut$_{1+2}$.SPYΔN-His$_6$ (Kif/endo F-treated) | 209.7 | <40 | <40 | <40 | 425.5 | <40 | <40 | 174.2 | 54.2 |

**Against ED43 HCVpp**

| Antigen | ID50 titers (week 11) M1 | M2 | M3 | M4 | M5 | M6 | M7 | M8 | Geometric Mean |
|---|---|---|---|---|---|---|---|---|---|
| HCV-1 sE1E2.Cut$_{1+2}$.SPYΔN-His$_6$ | 51.23 | 63.52 | 203.6 | <40 | 45.69 | <40 | 156.4 | 297.6 | 65.3 |
| HCV-1 sE1E2.Cut$_{1+2}$.SPYΔN-His$_6$ (Kif/endo F-treated) | 381.5 | <40 | <40 | <40 | 793.6 | <40 | <40 | <40 | 31.6 |

**Against 5.2.1 HCVpp**

| Antigen | ID50 titers (week 11) M1 | M2 | M3 | M4 | M5 | M6 | M7 | M8 | Geometric Mean |
|---|---|---|---|---|---|---|---|---|---|
| HCV-1 sE1E2.Cut$_{1+2}$.SPYΔN-His$_6$ | 175.1 | 83.23 | 1809 | 310.9 | 51.5 | <40 | 1506 | 595.6 | 231.7 |
| HCV-1 sE1E2.Cut$_{1+2}$.SPYΔN-His$_6$ (Kif/endo F-treated) | 1857 | 44.87 | 487.6 | <40 | 6429 | <40 | <40 | <40 | 72.6 |

#### Statistical analysis

**Against 3.1.2 HCVpp**

| Unpaired t test (w11) | Statistics | P Value |
|---|---|---|
| HCV-1 sE1E2.Cut$_{1+2}$.SPYΔN-His$_6$ vs. HCV-1 sE1E2.Cut$_{1+2}$.SPYΔN-His$_6$ (Kif/endo F-treated) | ns | 0.2539 |

**Against 4.2.2 HCVpp**

| Unpaired t test (w11) | Statistics | P Value |
|---|---|---|
| HCV-1 sE1E2.Cut$_{1+2}$.SPYΔN-His$_6$ vs. HCV-1 sE1E2.Cut$_{1+2}$.SPYΔN-His$_6$ (Kif/endo F-treated) | ns | 0.4558 |

**Against ED43 pseudovirus**

| Unpaired t test (w11) | Statistics | P Value |
|---|---|---|
| HCV-1 sE1E2.Cut$_{1+2}$.SPYΔN-His$_6$ vs. HCV-1 sE1E2.Cut$_{1+2}$.SPYΔN-His$_6$ (Kif/endo F-treated) | ns | 0.5459 |

**Against 5.2.1 pseudovirus**

| Unpaired t test (w11) | Statistics | P Value |
|---|---|---|
| HCV-1 sE1E2.Cut$_{1+2}$.SPYΔN-His$_6$ vs. HCV-1 sE1E2.Cut$_{1+2}$.SPYΔN-His$_6$ (Kif/endo F-treated) | ns | 0.444 |

**g** Sera from mice immunized with glycan-modified HCV-1 sE1E2.Cut$_{1+2}$.SPYΔN-10GS-FR SApNP vaccines against H77 HCVpp

### Figure. S6

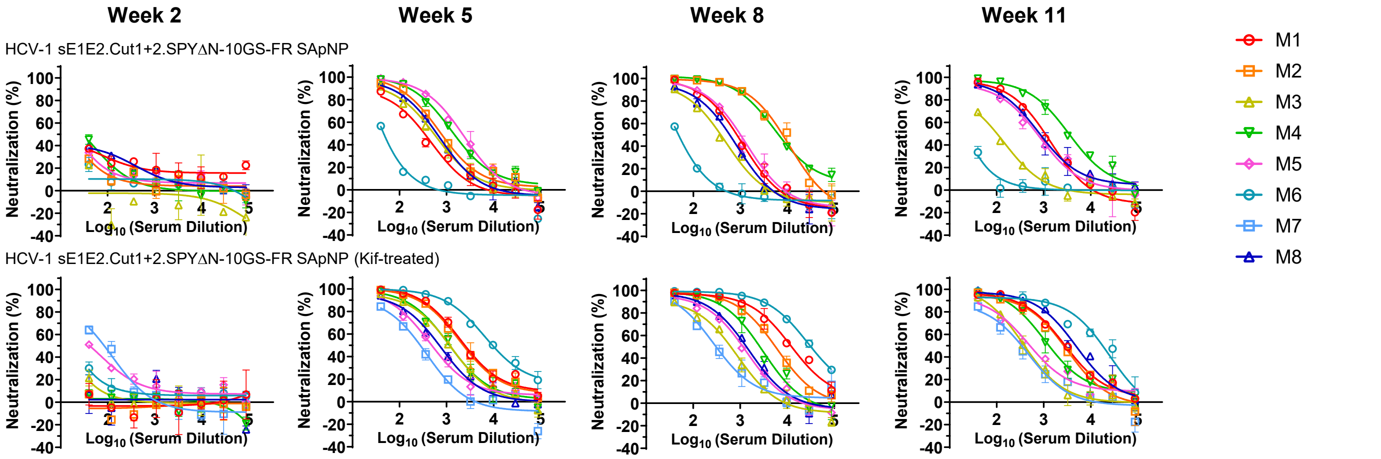


#### Mouse serum neutralizing $ID_{50}$ titers

**Week 2**

| Antigen | ID50 titers (week 2) M1 | M2 | M3 | M4 | M5 | M6 | M7 | M8 | Geometric Mean |
|---|---|---|---|---|---|---|---|---|---|
| HCV-1 sE1E2.Cut$_{1+2}$.SPYΔN-10GS-FR SApNP | <40 | <40 | <40 | <40 | <40 | <40 | N/A | <40 | N/A |
| HCV-1 sE1E2.Cut$_{1+2}$.SPYΔN-10GS-FR SApNP (Kif-treated) | <40 | <40 | <40 | <40 | 52.39 | <40 | 76.51 | <40 | N/A |

**Week 5**

| Antigen | ID50 titers (week 5) M1 | M2 | M3 | M4 | M5 | M6 | M7 | M8 | Geometric Mean |
|---|---|---|---|---|---|---|---|---|---|
| HCV-1 sE1E2.Cut$_{1+2}$.SPYΔN-10GS-FR SApNP | 289.4 | 789.9 | 509.3 | 1491 | 2014 | <40 | N/A | 580.7 | 501.8 |
| HCV-1 sE1E2.Cut$_{1+2}$.SPYΔN-10GS-FR SApNP (Kif-treated) | 2714 | 2467 | 1101 | 1178 | 516.5 | 10841 | 241.6 | 670.6 | 1294.3 |

**Week 8**

| Antigen | ID50 titers (week 8) M1 | M2 | M3 | M4 | M5 | M6 | M7 | M8 | Geometric Mean |
|---|---|---|---|---|---|---|---|---|---|
| HCV-1 sE1E2.Cut$_{1+2}$.SPYΔN-10GS-FR SApNP | 729 | 6993 | 337.5 | 7003 | 866.5 | 40.53 | N/A | 505.7 | 802.3 |
| HCV-1 sE1E2.Cut$_{1+2}$.SPYΔN-10GS-FR SApNP (Kif-treated) | 12393 | 5474 | 539.6 | 2433 | 1032 | 32190 | 327.5 | 1277 | 2435.4 |

**Week 11**

| Antigen | ID50 titers (week 11) M1 | M2 | M3 | M4 | M5 | M6 | M7 | M8 | Geometric Mean |
|---|---|---|---|---|---|---|---|---|---|
| HCV-1 sE1E2.Cut$_{1+2}$.SPYΔN-10GS-FR SApNP | 931.1 | N/A | 95.36 | 3431 | 679.5 | <40 | N/A | 848.9 | 376.1 |
| HCV-1 sE1E2.Cut$_{1+2}$.SPYΔN-10GS-FR SApNP (Kif-treated) | 2903 | 2432 | 395.5 | 1424 | 546.4 | 15035 | 293.8 | 4700 | 1609.8 |

#### Statistical analysis

| Unpaired t test (w5) | Statistics | P Value |
|---|---|---|
| HCV-1 sE1E2.Cut$_{1+2}$.SPYΔN-10GS-FR SApNP vs. HCV-1 sE1E2.Cut$_{1+2}$.SPYΔN-10GS-FR SApNP (Kif-treated) | ns | 0.2437 |

| Unpaired t test (w8) | Statistics | P Value |
|---|---|---|
| HCV-1 sE1E2.Cut$_{1+2}$.SPYΔN-10GS-FR SApNP vs. HCV-1 sE1E2.Cut$_{1+2}$.SPYΔN-10GS-FR SApNP (Kif-treated) | ns | 0.305 |

| Unpaired t test (w11) | Statistics | P Value |
|---|---|---|
| HCV-1 sE1E2.Cut$_{1+2}$.SPYΔN-10GS-FR SApNP vs. HCV-1 sE1E2.Cut$_{1+2}$.SPYΔN-10GS-FR SApNP (Kif-treated) | ns | 0.2572 |

**h** Sera from mice immunized with glycan-modified HCV-1 sE1E2.Cut$_{1+2}$.SPYΔN-10GS-FR SApNP vaccines against heterologous HCVpps

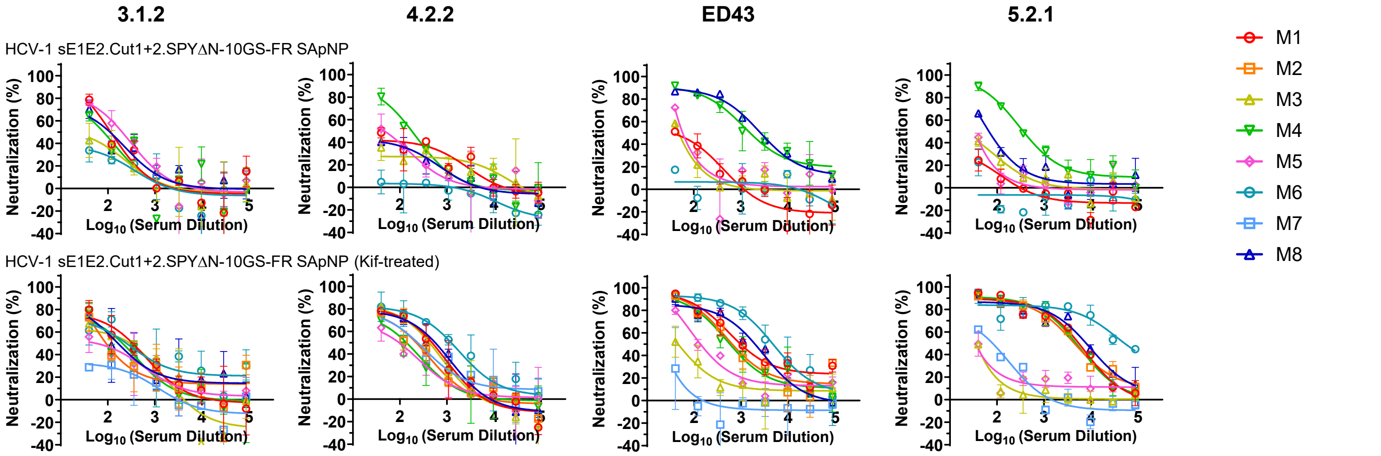


#### Mouse serum neutralizing $ID_{50}$ titers

**Against 3.1.2 HCVpp**

| Antigen | ID50 titers (week 11) M1 | M2 | M3 | M4 | M5 | M6 | M7 | M8 | Geometric Mean |
|---|---|---|---|---|---|---|---|---|---|
| HCV-1 sE1E2.Cut$_{1+2}$.SPYΔN-10GS-FR SApNP | 109.6 | N/A | 46.54 | 76.85 | 181.5 | <40 | N/A | 100.4 | 76.7 |
| HCV-1 sE1E2.Cut$_{1+2}$.SPYΔN-10GS-FR SApNP (Kif-treated) | 258.1 | 119.5 | 178 | 168.7 | 91.83 | 278.1 | <40 | 189.7 | 137.3 |

**Against 4.2.2 HCVpp**

| Antigen | ID50 titers (week 11) M1 | M2 | M3 | M4 | M5 | M6 | M7 | M8 | Geometric Mean |
|---|---|---|---|---|---|---|---|---|---|
| HCV-1 sE1E2.Cut$_{1+2}$.SPYΔN-10GS-FR SApNP | 71.04 | N/A | <40 | 166.3 | 56.53 | <40 | N/A | 47.4 | 36.9 |
| HCV-1 sE1E2.Cut$_{1+2}$.SPYΔN-10GS-FR SApNP (Kif-treated) | 362.7 | 184.8 | 317.5 | 123.3 | 94.91 | 1057 | 220.7 | 403.3 | 263.8 |

**Against ED43 HCVpp**

| Antigen | ID50 titers (week 11) M1 | M2 | M3 | M4 | M5 | M6 | M7 | M8 | Geometric Mean |
|---|---|---|---|---|---|---|---|---|---|
| HCV-1 sE1E2.Cut$_{1+2}$.SPYΔN-10GS-FR SApNP | 53.91 | N/A | 43.82 | 1913 | 55.39 | <40 | N/A | 2677 | 127.5 |
| HCV-1 sE1E2.Cut$_{1+2}$.SPYΔN-10GS-FR SApNP (Kif-treated) | 1303 | 748.7 | 53.08 | 642.1 | 167.9 | 4630 | <40 | 1583 | 374.1 |

**Against 5.2.1 HCVpp**

| Antigen | ID50 titers (week 11) M1 | M2 | M3 | M4 | M5 | M6 | M7 | M8 | Geometric Mean |
|---|---|---|---|---|---|---|---|---|---|
| HCV-1 sE1E2.Cut$_{1+2}$.SPYΔN-10GS-FR SApNP | <40 | N/A | <40 | 402.8 | <40 | <40 | N/A | 70.75 | 32.5 |
| HCV-1 sE1E2.Cut$_{1+2}$.SPYΔN-10GS-FR SApNP (Kif-treated) | 4716 | 4345 | <40 | 4361 | <40 | 38730 | 73.25 | 6964 | 1062.2 |

#### Statistical analysis

**Against 3.1.2 HCVpp**

| Unpaired t test (w11) | Statistics | P Value |
|---|---|---|
| HCV-1 sE1E2.Cut$_{1+2}$.SPYΔN-10GS-FR SApNP vs. HCV-1 sE1E2.Cut$_{1+2}$.SPYΔN-10GS-FR SApNP (Kif-treated) | ns | 0.0853 |

**Against 4.2.2 HCVpp**

| Unpaired t test (w11) | Statistics | P Value |
|---|---|---|
| HCV-1 sE1E2.Cut$_{1+2}$.SPYΔN-10GS-FR SApNP vs. HCV-1 sE1E2.Cut$_{1+2}$.SPYΔN-10GS-FR SApNP (Kif-treated) | * | 0.0483 |

**Against ED43 HCVpp**

| Unpaired t test (w11) | Statistics | P Value |
|---|---|---|
| HCV-1 sE1E2.Cut$_{1+2}$.SPYΔN-10GS-FR SApNP vs. HCV-1 sE1E2.Cut$_{1+2}$.SPYΔN-10GS-FR SApNP (Kif-treated) | ns | 0.6496 |

**Against 5.2.1 HCVpp**

| Unpaired t test (w11) | Statistics | P Value |
|---|---|---|
| HCV-1 sE1E2.Cut$_{1+2}$.SPYΔN-10GS-FR SApNP vs. HCV-1 sE1E2.Cut$_{1+2}$.SPYΔN-10GS-FR SApNP (Kif-treated) | ns | 0.1952 |

**Figure. S6**

#### i Week-11 sera from female mice immunized with HCV-1 sE1E2.Cut$_{1+2}$.SPYΔN-His$_6$ dimer/AH against H77 HCVpp

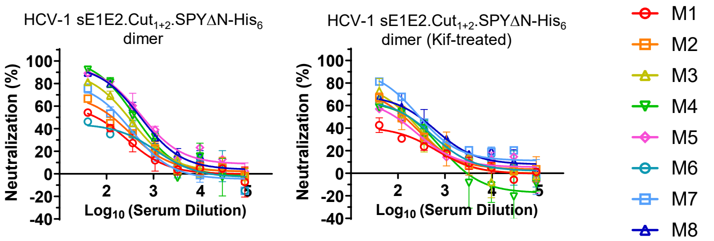


##### Mouse serum neutralizing ID$_{50}$ titers

**Week 11**

| Antigen | ID50 titers (week 11) | | | | | | | | Geometric Mean |
|---|---|---|---|---|---|---|---|---|---|
| | M1 | M2 | M3 | M4 | M5 | M6 | M7 | M8 | |
| HCV-1 sE1E2.Cut$_{1+2}$.SPYΔN-His$_6$ | 74.22 | 124.1 | 284 | 418.1 | 629.6 | 63.18 | 166.9 | 535.8 | 210.7 |
| HCV-1 sE1E2.Cut$_{1+2}$.SPYΔN-His$_6$ (Kif-treated) | 45.21 | 129.3 | 148.2 | 133.2 | 97.89 | 152.5 | 312.3 | 242.8 | 137.9 |

##### Statistical analysis

| Unpaired t test (w11) | Statistics | P Value |
|---|---|---|
| HCV-1 sE1E2.Cut$_{1+2}$.SPYΔN-His$_6$ vs. HCV-1 sE1E2.Cut$_{1+2}$.SPYΔN-His$_6$ (Kif-treated) | ns | 0.1396 |

#### j Week-11 sera from male mice immunized with HCV-1 sE1E2.Cut$_{1+2}$.SPYΔN-His$_6$ dimer/AH against H77 HCVpp

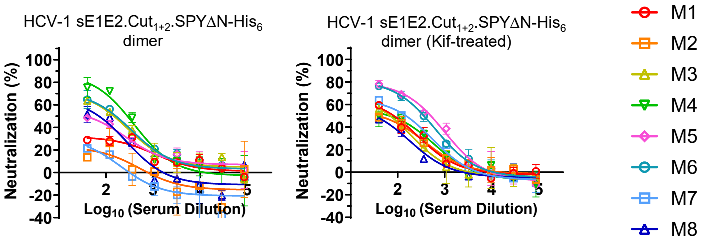


##### Mouse serum neutralizing ID$_{50}$ titers

**Week 11**

| Antigen | ID50 titers (week 11) | | | | | | | | Geometric Mean |
|---|---|---|---|---|---|---|---|---|---|
| | M1 | M2 | M3 | M4 | M5 | M6 | M7 | M8 | |
| HCV-1 sE1E2.Cut$_{1+2}$.SPYΔN-His$_6$ | <40 | <40 | 142.8 | 262.9 | 70.67 | 149.5 | <40 | 72.39 | 56.4 |
| HCV-1 sE1E2.Cut$_{1+2}$.SPYΔN-His$_6$ (Kif-treated) | 84.46 | 69.56 | 71.21 | 89.11 | 386.1 | 298.4 | 123.8 | 48.47 | 112.6 |

##### Statistical analysis

| Unpaired t test (w11) | Statistics | P Value |
|---|---|---|
| HCV-1 sE1E2.Cut$_{1+2}$.SPYΔN-His$_6$ vs. HCV-1 sE1E2.Cut$_{1+2}$.SPYΔN-His$_6$ (Kif-treated) | ns | 0.3444 |

#### k Week-11 sera from female mice immunized with HCV-1 sE1E2.Cut$_{1+2}$.SPYΔN-10GS-FR SApNP/AH against H77 HCVpp

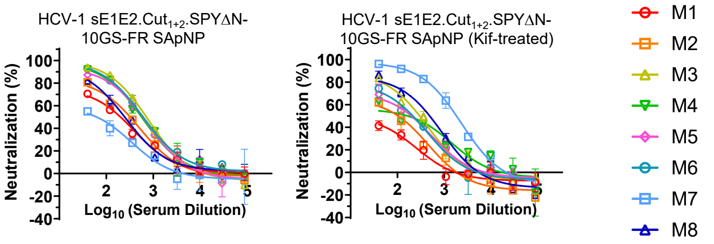


##### Mouse serum neutralizing ID$_{50}$ titers

**Week 11**

| Antigen | ID50 titers (week 11) | | | | | | | | Geometric Mean |
|---|---|---|---|---|---|---|---|---|---|
| | M1 | M2 | M3 | M4 | M5 | M6 | M7 | M8 | |
| HCV-1 sE1E2.Cut$_{1+2}$.SPYΔN-10GS-FR SApNP | 173.3 | 272 | 673.6 | 544.8 | 538.2 | 614.5 | 74.19 | 245.6 | 317.9 |
| HCV-1 sE1E2.Cut$_{1+2}$.SPYΔN-10GS-FR SApNP (Kif-treated) | <40 | 88.65 | 254.9 | 143.8 | 159.1 | 176.1 | 1616 | 356.7 | 195.0 |

##### Statistical analysis

| Unpaired t test (w11) | Statistics | P Value |
|---|---|---|
| HCV-1 sE1E2.Cut$_{1+2}$.SPYΔN-10GS-FR SApNP vs. HCV-1 sE1E2.Cut$_{1+2}$.SPYΔN-10GS-FR SApNP (Kif-treated) | ns | 0.8535 |

**Fig. S6. Impact of protein expression system, sex-specific differences, and injection route on the immunogenicity of HCV-1 sE1E2.Cut$_{1+2}$.SPYΔN vaccines in mice.** (**a**) Neutralization curves of mouse sera from glycan-modified HCV-1 sE1E2.Cut$_{1+2}$.SPYΔN-His$_6$ dimer (produced in HEK293F cells) vaccine groups (n = 8 mice/group) against H77 HCVpp. The administered dose was 200 µl of antigen/AH + CpG adjuvant mix containing 10 µg of immunogen and 100 µl of adjuvant via the intraperitoneal route. Mice were immunized at weeks 0, 3, 6 and 9 with 3-week intervals. (**b**) (Top) Summary of geometric mean ID$_{50}$ titers measured for glycan-modified HCV-1 sE1E2.Cut$_{1+2}$. SPYΔN-His$_6$ dimer (produced in HEK293F cells) vaccine groups against H77 HCVpp. Color coding indicates the level of ID$_{50}$ titers (white: no neutralization; green to red: low to high neutralization). ID$_{50}$ values were derived by setting the minimum/maximum % neutralization to 0.0/100.0 as constraints. (Bottom) Summary of statistical analysis performed for each timepoint. (**c**) Neutralization curves of mouse sera from glycan-modified HCV-1 sE1E2.Cut$_{1+2}$.SPYΔN-His$_6$ dimer (produced in HEK293F cells) vaccine groups at week 11 against heterologous 3.1.2, 4.2.2, ED43, and 5.2.1 HCVpps. (**d**) Summary of geometric mean ID$_{50}$ values and statistical analysis. (**e**) Neutralization curves of mouse sera from glycan-modified HCV-1 sE1E2.Cut$_{1+2}$.SPYΔN-His$_6$ dimer vaccine group against H77 HCVpp from an independent study. Summary of geometric mean ID$_{50}$ values and statistical analysis. (**f**) Neutralization curves of mouse sera from glycan-modified HCV-1 sE1E2.Cut$_{1+2}$.SPYΔN-His$_6$ dimer vaccine groups at week 11 against heterologous 3.1.2, 4.2.2, ED43, and 5.2.1 HCVpps. Summary of geometric mean ID$_{50}$ values and statistical analysis. (**g**) Neutralization curves of mouse sera from glycan-modified HCV-1 sE1E2.Cut$_{1+2}$.SPYΔN-10GS-FR SApNP vaccine group against H77 HCVpp from an independent study. Summary of geometric mean ID$_{50}$ values and statistical analysis. (**h**) Neutralization curves of mouse sera from glycan-modified HCV-1 sE1E2.Cut$_{1+2}$.SPYΔN-10GS-FR SApNP vaccine groups at week 11 against heterologous 3.1.2, 4.2.2, ED43, and 5.2.1 HCVpps. Summary of geometric mean ID$_{50}$ values and statistical analysis. Neutralization curves of mouse sera from glycan-modified HCV-1 sE1E2.Cut$_{1+2}$.SPYΔN-His$_6$ dimer (produced in ExpiCHO cells) vaccine group in (**i**) female and (**j**) male mice against H77 HCVpp. The administered dose was 80 µl of antigen/AH adjuvant mix containing 10 µg of antigen and 40 µl of adjuvant via the footpad intradermal (i.d.) route. (**k**) Neutralization curves of mouse sera from glycan-modified HCV-1 sE1E2.Cut$_{1+2}$.SPYΔN-10GS-FR SApNP (produced in ExpiCHO cells) vaccine group in female mice against H77 HCVpp. Summary of geometric mean ID$_{50}$ values and statistical analysis. Error bars represent the difference between duplicate values at each concentration tested for each sample. ID$_{50}$ values were calculated in GraphPad Prism 10.3.1. Data were analyzed using one-way ANOVA, followed by Tukey's multiple comparison post hoc test for each timepoint. Two-tailed unpaired t-tests were used to compare the geometric means of two groups. Statistical significance is indicated as follows: ns (not significant) and $*p < 0.05$.

**Table S1. Antibodies and ligand used for antigenic profiling of HCV-1 sE1E2.Cut$_{1+2}$.SPYΔN antigens and nanoparticles.** [a]

| Antibody name | Antigenic regions (AR) or antigenic sites (AS) | Interacting residues | References (PMID) |
|---|---|---|---|
| A4 (mouse) | AS197 (E1 residues 197-207) | S198, G199, Y201, H202, T204 | Dubuisson et al. 1994 (8083956); Gopel et al. 2017 (29253863) |
| HCV1 | AS412 (E2 residues 412-423) | L413, N415, T416, G418, W420 | Kong et al., 2012 (22623528); Gopel et al. 2017 (29253863) |
| AR1B | AR1, E2 Ig scaffold β strands 6 and 7 (E2 residues 536-555) | V536, F537, V538, L539, N540, T542, R543, P544, P545, G547, N548, W549, F550, G551, C552, T553, W554, M555 | Law et al. 2008 (18064037); Gopel et al. 2017 (29253863) |
| AR2A | AR2, E2 back layer β strand 11 (E2 residues 625-633) | T625, I626, F627, K628, V629, R630, M631, Y632, V633 | Law et al. 2008 (18064037); Gopel et al. 2017 (29253863) |
| AR3C | AR3, E2 neutralizing face involving front layer (E2 residues 421-459) and CD81 binding loop (E2 residues 519-535) | S424, T425, L427, N428, C429, N430, S432, N434, G436, W437, L438, G440, L441, Y443, S450, L459, T519, D520, R521, S522, G523, P525, T526, Y527, S528, W529, G530, N532, D533, D535 | Law et al. 2008 (18064037); Gopel et al. 2017 (29253863); Tzarum et al. 2019 (30613781) |
| HEPC74 | AR3 | H421, L427, D431, S432, H434, G436, L438, A439, F442, Y443, K446, F447, N448, W529, E531 | Chen et al. 2021 (33675683) |
| HC1AM | AR3 | W420, I422, N423, L427, N428, C429, K500, S528, W529, A531, D533, Y527, W616 | Tzarum et al. 2020 (32754640) |
| RM2-01 | AR3 | I422, L441, F442, Y443, I438, T439, W529, E531, Y613 | Chen et al. 2021 (33675683) |
| RM11-43 | AR3 | I422, L427, F442, Y443, Q444, I438, T439, P505, W529, P612 | Chen et al. 2021 (33675683) |
| AR4A | AR4, E1E2 interface involving E2 residue 698 | Y201, N205, L654, R657, D658. R659, P676, F679, L689, I690, L692, D698, V699, Q700 | Giang et al. 2012 (22492964); Gopel et al. 2017 (29253863) |
| AR5A | AR5, E1E2 interface involving E2 residues 639 and 665 | Y201, N205, R639, L654, R657, D658, R659, L665, P676, F679, L689, I690, L692, D698, V699, Q700 | Giang et al. 2012 (22492964); Gopel et al. 2017 (29253863) |
| CD81-Fc [b] | CD81 binding loop (E2 residues 519-535) | T519, D520, R521, S522, G523, A524, P525, T526, Y527, S528, W529, G530, A531, D533, D535 | Gopel et al. 2017 (29253863) |

[a] All antibodies and CD81-Fc were included in the ELISA analysis of HCV-1 sE1E2.Cut1+2.SPYΔN antigens and nanoparticles; however, A4, a murine antibody targeting E1, was excluded from the BLI analysis.

**Table S2. Cryo-EM data collection.**

| Data Collection | HEPC74-AR4A-sE1E2.$Cut_{1+2}$.SPYΔN-$His_6$ Vitrojet | AR3C-AR4A-sE1E2.$Cut_{1+2}$.SPYΔN-$His_6$ Vitrojet | AR3C-AR4A-sE1E2.$Cut_{1+2}$.SPYΔN-$His_6$ Vitrobot |
|---|---|---|---|
| Magnification (nominal) | 81,000 | 81,000 | 81,000 |
| Voltage (kV) | 300 | 300 | 300 |
| Electron exposure (e–/Å$^2$) | 50.5 | 50.5 | 50.03 |
| Defocus range (µm) | 1 - 2 | 1 - 2 | 1 - 2 |
| Pixel size (Å) | 1.09 | 1.09 | 1.09 |
| Symmetry imposed | C1 | C1 | |
| Initial particle images (no.) | 14,430,734 | 2,446,658 | 8,699,347 |
| Final particle images (no.) | 36,896 | 89,013 | 85,859 |
| Map resolution (Å) | 6.30 | 4.97 | |
| FSC threshold | 0.143 | 0.143 | |
| Map resolution range (Å) | 4.188 - 12.77 | 2.281 - 13.161 | |
| EMDB ID | EMD-70622 | EMD-70623 | — |

**Table S3. Epitope information derived from mutagenesis vs. nsEM-based epitope mapping.** [a]

| HCV mAb | Antigenic region | Binding residues on HCV-1 E1E2 from publish structural analysis[a] | Binding residues on H77 E1E2 from mutagenesis information[b] | References |
|---|---|---|---|---|
| AR1B | AR1, E2 Ig scaffold β strands 6 and 7 (E2 residues 536-555) | No structure available. | V536, F537, V538, L539, N540, T542, R543, P544, P545, G547, N548, W549, F550, G551, C552, T553, W554, M555, T556, G559, T561 | Law et al. 2008; Gopel et al. 2017 |
| AR2A | AR2, E2 back layer β strand 11 (E2 residues 625-633) | No structure available. | T625, I626, F627, K628, V629, R630, M631, Y632, V633 | Law et al. 2008; Gopel et al. 2017 |
| AR3C | AR3, E2 neutralizing face involving front layer (E2 residues 421-459) and CD81 binding loop (E2 residues 519-535) | L427, C429, E431, S432, Y443 | S424, T425, L427, N428, C429, N430, S432, N434, G436, W437, L438, G440, L441, Y443, S450, L459, T519, D520, R521, S522, G523, P525, T526, Y527, S528, W529, G530, N532, D533, D535 | Law et al. 2008; Gopel et al. 2017; Kong et al. 2013; Tzarum et al. 2019 |
| AR4A | AR4, E1E2 interface involving E2 residue 698 | W646, R648, G649, L667, Q671, P676, I696, D698 | Y201, N205, L654, R657, D658. R659, P676, F679, L689, I690, L692, D698, V699, Q700 | Giang et al. 2012; Gopel et al. 2017; Alba Torrents de la Peña et al. 2022. |
| AR5A | AR5, E1E2 interface involving E2 residues 639 and 665 | No structure available. | Y201, N205, R639, L654, R657, D658, R659, L665, P676, F679, L689, I690, L692, D698, V699, Q700 | Giang et al. 2012; Gopel et al. 2017 |

[a] The involved residues on E2 were analyzed using structures 4MWF and 7T6X for AR3C and AR4A, respectively.

[b] Residues overlapping with the epitope footprint and facing the density-fitted antibody model (indicative of direct interactions) are labeled in red, while those located at the E1E2 interface but may not be involved in direct interactions were labeled in orange.
